# Network Analysis and Transcriptome Profiling Identify Autophagic and Mitochondrial Dysfunctions in SARS-CoV-2 Infection

**DOI:** 10.1101/2020.05.13.092536

**Authors:** Komudi Singh, Yun-Ching Chen, Jennifer T Judy, Fayaz Seifuddin, Ilker Tunc, Mehdi Pirooznia

**Author notes:** Correspondence: Mehdi Pirooznia.

## Abstract

Analyzing host transcriptional changes in response to SARS-CoV-2 infection will help delineate biological processes underlying viral pathogenesis. Comparison of expression profiles of lung cell lines A549 (infected with either SARS-CoV-2 (with ACE2 expression)) or Influenza A virus (IAV)) and Calu3 (infected with SARS-CoV-2 or MERS-CoV) revealed upregulation of the antiviral interferon signaling in all three viral infections. However, perturbations in inflammatory, mitochondrial, and autophagy processes were specifically observed in SARS-CoV-2 infected cells. Validation of findings from cell line data revealed perturbations in autophagy and mitochondrial processes in the infected human nasopharyngeal samples. Specifically, downregulation of mTOR expression, mitochondrial ribosomal, mitochondrial complex I, and lysosome acidification genes were concurrently observed in both infected cell lines and human datasets. Furthermore, SARS-CoV-2 infection impedes autophagic flux by upregulating GSK3B in lung cell lines, or by downregulating autophagy genes, SNAP29 and lysosome acidification genes in human samples, contributing to increased viral replication. Therefore, drugs targeting lysosome acidification or autophagic flux could be tested as intervention strategies. Additionally, downregulation of MTFP1 (in cell lines) or SOCS6 (in human samples) results in hyperfused mitochondria and impede proper interferon response. Coexpression networks analysis identifies correlated clusters of genes annotated to inflammation and mitochondrial processes that are misregulated in SARS-CoV-2 infected cells. Finally, comparison of age stratified human gene expression data revealed impaired upregulation of chemokines, interferon stimulated and tripartite motif genes that are critical for antiviral signaling. Together, this analysis has revealed specific aspects of autophagic and mitochondrial function that are uniquely perturbed in SARS-CoV-2 infection.

## INTRODUCTION

### Epidemiology

Severe Acute Respiratory Syndrome Coronavirus 2 (SARS-CoV-2) is the virus that causes the current global pandemic, coronavirus disease (COVID-19). COVID-19 presents as a wide range of clinical manifestations, ranging from asymptomatic to respiratory failure or multiorgan and systemic manifestations (1–3). The viral pneumonia outbreak caused by SARS-CoV-2 was first identified in Wuhan, China in December 2019 (4). Since then, the virus has continued to spread globally, with a current transmissibility estimate (R_0_) between 3-4 (5, 6). According to the World Health Organization (7) as of late April 2020, there have been over 2.8 million confirmed cases and more than 196,000 confirmed deaths across 213 countries/areas/territories. No treatments currently exist, and management strategies include supportive medical care for existing cases and social distancing for prevention. Understanding this novel pathogen and the host response it elicits is crucial to combatting the emerging threat to public health.

### Human Coronavirus (hCoV) Phylogeny

SARS-CoV-2 is the 7^th^ and most recent addition to human coronaviruses (hCoVs), which include four globally endemic hCoVs that cause a substantial portion of upper respiratory infections (229E, OC43, HKU1, and NL63), as well as two other highly pathogenic strains that have also caused recent pandemics (SARS-CoV and MERS-CoV (5, 8) in 2002-2003 and 2012, respectively (9)). All seven hCoVs are single-stranded, positive-sense RNA viruses. They all have zoonotic origins, with bats as the evolutionary reservoir host of five viruses (229E, NL63, SARS-CoV, MERS-CoV, and SARS-CoV-2). In some cases, there are intermediate and amplifying host species as well (5). Although SARS-CoV-2 is phylogenetically similar to both MERS-CoV, and SARS-CoV (10), there are biological differences. Notably, although SARS-CoV-2 has a lower, but yet undetermined mortality rate, it is distinctly more contagious than these other highly pathogenic hCoVs, causing vastly different epidemiological dynamics. In fact, MERS-CoV was largely propagated by camel-to-human transmissions, as the virus was never able to fully adapt to optimal human-to-human transmission (11).

### Pathogenicity

As obligate parasites, the viruses, while evading the host cell immune response, should encode enough proteins to ensure replication, and spread, by relying on the host cell’s machinery. These processes require an intricate series of interaction between the virus and host (12). While many of the elicited responses are common across pathogens, each virus also creates a unique transcriptional profile (13). Functional distinctions across viruses may arise due to distinct processes utilized by a virus for cellular entry, or for host immune system evasion, or for replication and dissemination (14). While the immune response is essential to resolving the infection, dysregulation of the immune system can result in immunopathogenesis (15, 16). A dysregulated immune response is caused by rapid viral replication, cytokine storms, delayed interferon response, and macrophage infiltration and excessive proinflammatory cytokines (15). This immunopathogenesis mechanism is supported by the observation of decreased viral loads occurring with increased disease severity (9). Severity of illness for SARS-CoV-2 infections is likely impacted by both the direct cytotoxic effects of the virus, and the effectiveness of the complex host response (17, 18). However, efforts to understand the molecular mechanisms require further study, as the cause of unusually high morbidity and mortality of hCoVs remain unclear.

### Cell/Tissue Tropism of SARS-CoV-2

The hCoVs differentially infect the human respiratory tract. The low pathogenic hCoVs infect the upper respiratory tract, and the highly pathogenic hCoVs infect the lower respiratory tract (15). Consistent with this, SARS-CoV, SARS-CoV-2, and MERS-CoV were shown to differentially infect the lung alveolar cell subtypes in cynomolgus macaques (19) and SARS-CoV elicited distinct immune response in different tissues (20). Furthermore, cell tropism study of the SARS-CoV and SARS-CoV-2 in different cell type cultures could partially explain the symptomatic differences of these two virus infections (21). Single cell (sc) transcriptomic data of the COVID-19 lung tissue have been analyzed to identify the subset of cells most prone to the SARS-CoV-2 infection and the marker genes associated with the infected cells. One such study intriguingly identified upregulation of the receptor-angiotensin-converting enzyme 2 (ACE2) in the SARS-CoV-2 infected type II pneumocyte population of the lung cells as a potential mechanism facilitating virus infection (22). Several studies have linked the expression of ACE2 and TMPRSS2 with increased susceptibility to viral entry (23–25). Another study utilized the ACE2 and TMPRSS2 expression information at the single cell level to rank the cells based on their susceptibility to the SARS-CoV-2 infection (26). Consistent with the sc RNA seq data, dual inhibition of the host cell cysteine and serine proteases impeded viral entry into the cell (24). In addition to ACE2 and TMPRSS2, another study has shown the neuropilin-1 (NRP1) could act as host co factor and facilitate viral entry (27).

To delineate the host cell transcriptional response to the viral infection and potentially identify genes and biological processes (BPs) specifically impacted by SARS-CoV-2 infection, we have utilized gene expression information from several datasets across cell lines (28), and from human nasopharyngeal samples (29) classified into young and old groups that were positive for high or low viral loads (See Fig 1, study schema). Since A549 cells do not express the requisite ACE2 or TMPRSS2 receptors for viral entry, A549 cells transduced with human ACE2 (hACE2) infected with either 0.2 multiplicity of infection (MOI, low), or 2 MOI (high) of SARS-CoV-2 were utilized. These transformed cells revealed viral transcript reads at either MOIs indicating viral entry in A549 cells (28). Calu3 cells also revealed viral reads when infected with SARS-CoV-2 at 2 MOI (28). Viral infection in the transduced A549 cells and Calu3 cells presented in this dataset was confirmed by the evaluating the percent reads that aligned with the viral genome for each of the infected samples and has been published by Blanco-Melo et al., (28). Comparing pathway enrichment results of SARS-CoV-2, MERS-CoV, and influenza A virus (IAV) comparisons revealed upregulation of interferon signaling in all infected cell lines. However, the SARS-CoV-2 infection elicited a differential gene expression response that is unique to coronavirus infection, which included perturbations in inflammation, autophagy, and mitochondrial processes. To validate the findings from cell lines, expression profile of infected human nasopharyngeal samples was analyzed. To delineate age and viral load specific host cell response, the human data was classified into young (<40 years) and old (>60 years) that were positive with either high or low viral loads. For comparison with A549 cell data, expression profile of control vs. high viral load positive old age human samples were used. Consistent with the cell line data, the differentially expressed (DE) genes from control vs. infected human samples also annotated to inflammation, autophagy, and mitochondrial processes. It is likely that perturbation of the mitochondrial function and autophagy could negatively impact the host cells’ immune response against the viral infection leading to systemic inflammation (30, 31). Notably, we found downregulation of mTOR, mitochondrial ribosomal, mitochondrial complex I, and several lysosome acidification genes in both cell line and human datasets. However, specific genes regulating autophagic flux were differently expressed in both datasets. Consistent with the function of mTOR (32, 33), the autophagy genes were upregulated in infected cell lines. However, the autophagy flux impeding GSK3B was also upregulated. In the infected human samples, several autophagy genes, p62 and SNAP29 were downregulated. Together, these gene expression changes support that the autophagic flux is likely decreased in SARS-CoV-2 infected cells, which may contribute to viral propagation. Therefore, drugs increasing autophagic flux, or lysosome acidification could be tested as treatment strategies. Additionally, mitochondrial fission promoting MFTP1 and SOCS6 were downregulated in infected cell lines and human samples, respectively. This may contribute to hyperfused mitochondria and impaired interferon response. Furthermore, the gene expression profile of A549 cell line strongly correlated with the lung epithelial lineage basal and ionocyte cell types from the lung single cell (sc) RNA seq data. Using the age stratified human nasopharyngeal expression data, we have also delineated some age specific changes in antiviral signaling which may provide more insight into the age dependent differences in viral pathogenicity. Therefore, from this analysis we have identified some key aspects of autophagy and mitochondrial processes that are uniquely impacted in SARS-CoV-2 infection and are likely representative of host cells’ response to the infection.

**Figure 1.**
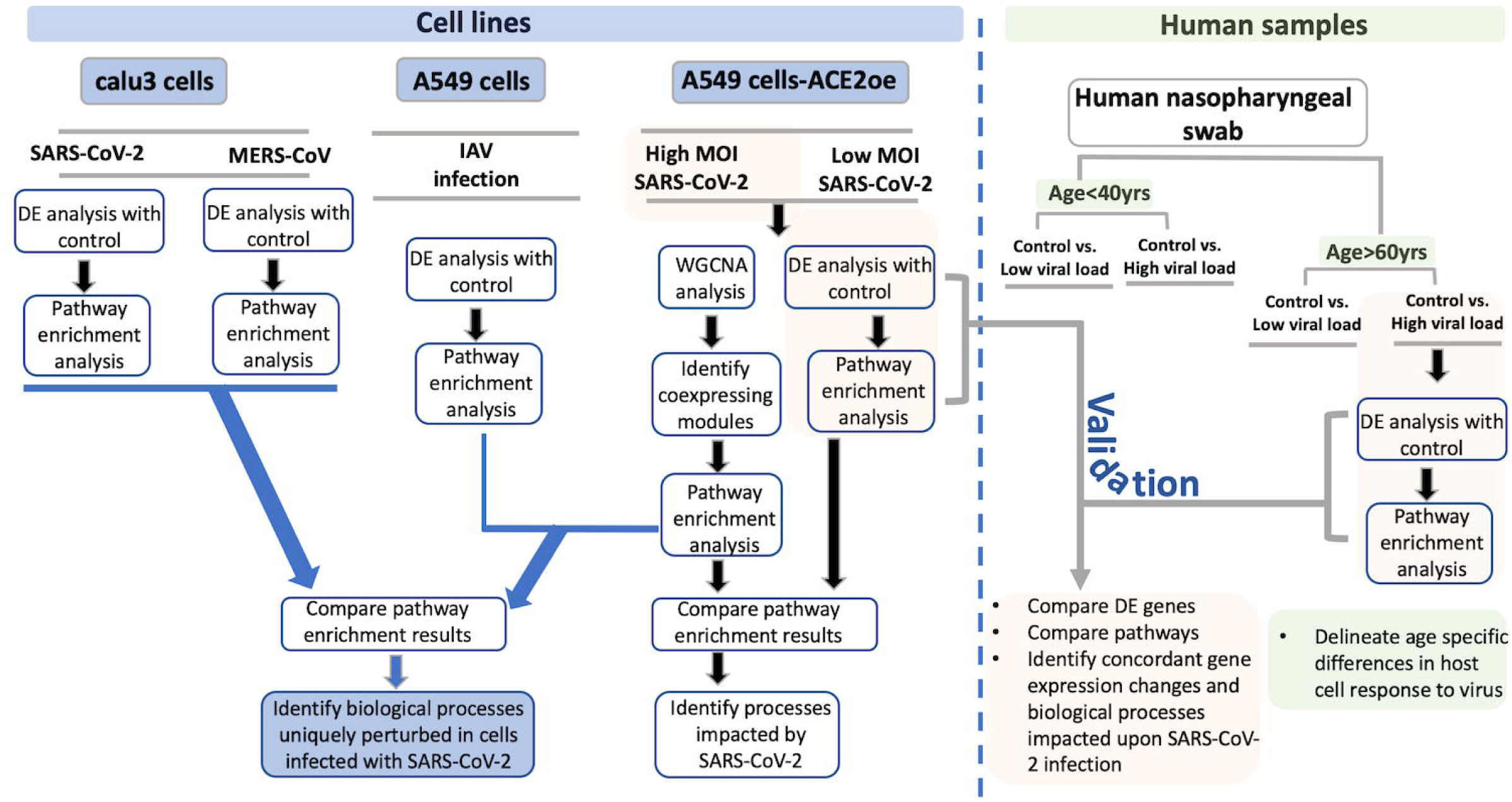
Study schema. A schema highlighting the various datasets used in the study and the downstream analysis performed for each dataset is shown as a flow-chart. The datasets utilized in this study is divided into two groups: Blue color boxes highlight the cell lines used in the study. The viruses used for infection are indicated between grey horizontal lines. The green box represents the human nasopharyngeal datasets and the age and viral load criteria used to classify these samples. For each pair of infected and control samples, DE analysis was performed to identify DE genes and pathway enrichment analysis was performed to reveal the biological processes to which the DE genes annotated to (shown in the flow chart). From the cell line data, the DE genes and pathway enrichment results were compared to identify BPs that were either uniquely perturbed in SARS-CoV-2 infection or were commonly perturbed in all viral infection (blue color lines with arrowhead leading to the blue color box). For validating the findings from the cell line data, the DE results from infected human nasopharyngeal samples were analyzed. Specifically, the DE results from A549 dataset (highlighted in orange box) was compared with the DE results from old age control vs. high viral load positive human nasopharyngeal samples (highlighted in orange box). This comparison will help identify concordant gene expression changes and BPs impacted in both datasets (grey color line with arrowhead leading to orange color box). Finally, using the age stratified human nasopharyngeal datasets, age-specific gene expression changes were delineated (green color boxes). DE: differential expression; BPs: biological processes; MOI: multiplicity of infection.

## RESULTS

To perform an in-depth analysis of the host cells’ transcriptional response to SARS-CoV-2 infection, several gene expression datasets from different cell lines and human samples were used. An overview of the datasets analyzed and compared are depicted in Figure 1 (study schema).

### Interferon autophagy, and mitochondrial processes are impacted in A549 cells infected with SARS-CoV-2

#### SARS-CoV-2 (high viral titer) vs Mock

Differential gene expression analysis of hACE2 receptor transduced A549 lung epithelial cell line that were either mock or infected with SARS-CoV-2 at higher viral titer (2MOI) (see methods, (28)) identified ∼8000 DE genes. The volcano plot profiles both up-regulated and down-regulated genes in the SARS-CoV-2 infected cells (Supp Fig S1A, https://doi.org/10.6084/m9.figshare.12272351 [Table S3]). Pathway enrichment analysis of the DE genes showed enrichment in a wide range of biological processes (Supp Fig S1B, https://doi.org/10.6084/m9.figshare.12272351 [Table S4]). These DE genes were classified into upregulated or downregulated following SARS-CoV-2 infection and analyzed by pathway enrichment analysis. Upregulated DE genes annotated to a wide range of pathways, notably including the interferon signaling, NFkB/cytokine signaling processes, and proteasomal degradation (Fig 2A, https://doi.org/10.6084/m9.figshare.12272351 [Table S4]). Heatmaps highlight the upregulation of genes in interferon, and cytokine processes, and perturbation of genes in the autophagy pathways (Fig 2B,2C,2D respectively). DE genes downregulated in the SARS-CoV-2 infected cells annotated to pathways primarily involving cell cycle and mitochondrial processes (Fig 2E, https://doi.org/10.6084/m9.figshare.12272351 [Table S4]). A heatmap shows that the expression of the genes in mitochondria-related processes, electron transport chain and respiration were mostly downregulated in SARS-CoV-2 infected cells (Fig 2F).

**Figure 2:**
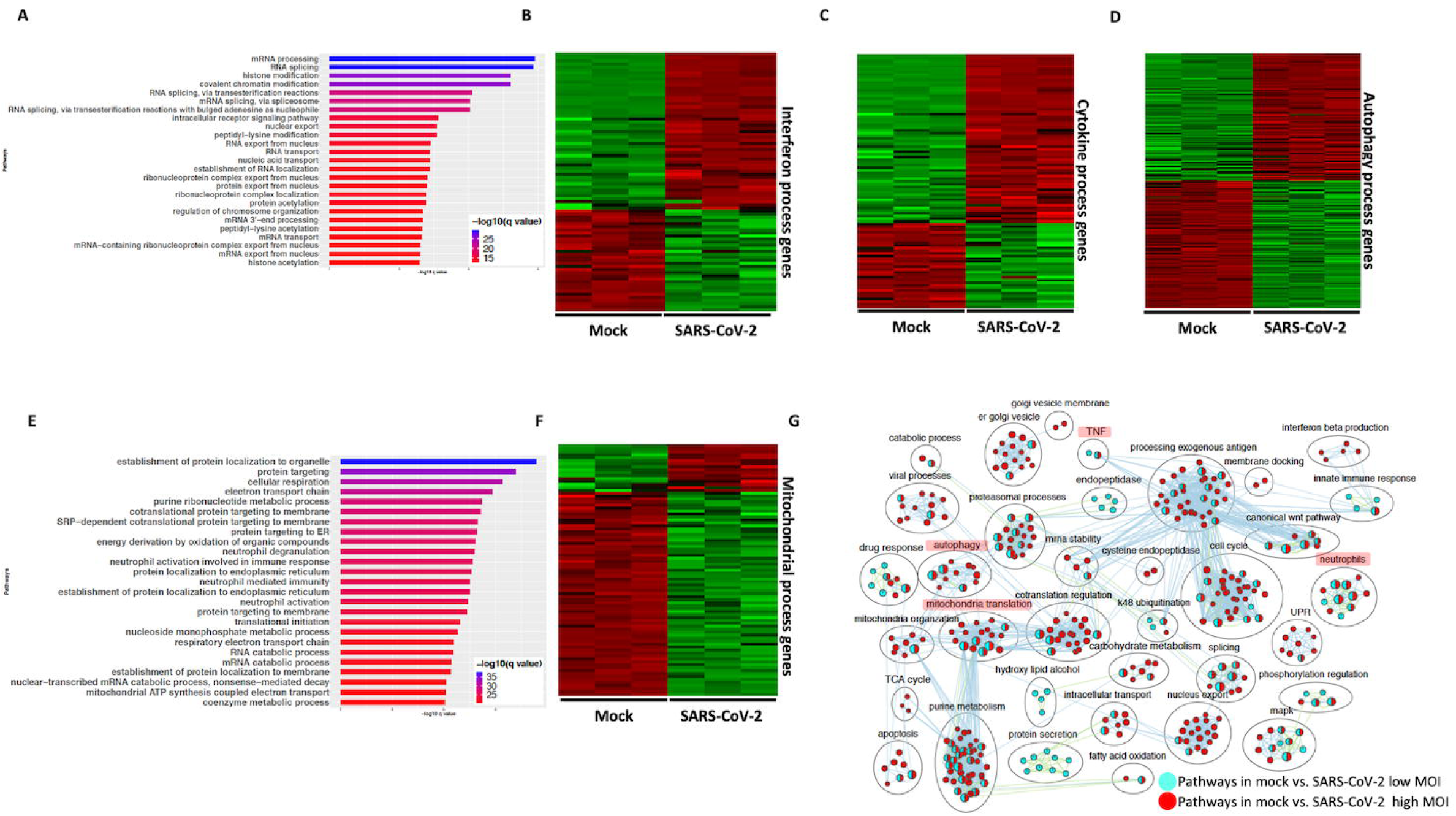
SARS-CoV-2 infection of hACE2 transduced lung epithelial A549 cells impacts expression of genes in interferon, cytokine and autophagic processes. **(A)** Top 25 pathways from the pathway enrichment analysis of gene upregulated in SARS-CoV-2 infected A549 cell is presented as a horizontal bar plot, where x axis represents the −log10 transformed q value and the color of the horizontal bar is scaled blue to red representing low to high q values, respectively. **(B)** Heatmap highlighting the expression of genes in the interferon processes in mock and infected cells. The red and green color bands represent up and downregulated genes, respectively. This heatmap shows that cytokine related genes were predominantly upregulated in infected cells. **(C)** Heatmap highlighting the expression of genes in the cytokine processes in mock and infected cells. The red and green color bands represent up and downregulated genes, respectively. This heatmap shows that interferon related genes were predominantly upregulated in infected cells. **(D)** Heatmap highlighting the expression of genes in the autophagy related processes in mock and infected cells. The red and green color bands represent up and downregulated genes, respectively. This heatmap shows that autophagy related genes were perturbed in infected cells. **(E)** Top 25 pathways from the pathway enrichment analysis of gene upregulated in SARS-CoV-2 infection is presented as a horizontal bar plot, where x axis represents the −log10 transformed q value and the color of the horizontal bar is scaled blue to red representing low to high q values, respectively. **(F)** Heatmap highlighting the expression of genes in the mitochondrial organization and translation in mock and infected cells. The red and green color bands represent up and downregulated genes, respectively. This heatmap shows that mitochondrial processes related genes were predominantly downregulated in infected cells. (G) Pathway enrichment summary map for mock vs. SARS-CoV-2 at high MOI (blue nodes) and low MOI (red nodes) comparisons. Each node represents a pathway/biological process (BP). The node size is proportional to the number of DE genes overlapping with the BP. The nodes that share genes are connected with edges. Single color nodes are pathways that are distinctly enriched by DE genes from one comparison. Two colored nodes are pathways enriched by DE genes from both comparisons. The label above each black circle summarizes the gene ontology (GO) terms of similar BPs present inside the circle. Notable groups of BPs associated with antigen processing, autophagy and mitochondria that were predominantly enriched by DE genes from Mock vs. SARS-CoV-2 (high MOI) comparison and are highlighted in red. BP associated interferon and UPRprocesses were also predominantly enriched by DE genes from Mock vs. SARS-CoV-2 (high MOI) comparison. MOI: multiplicity of infection; DE: differentially expressed; UPR: unfolded protein response.

#### SARS-CoV-2: Low viral titer vs high viral titer

Differential gene expression analysis of hACE2 transduced A549 cells infected with mock and a 10-fold lower viral titer of SARS-CoV-2 (see methods, (28)) was also performed. The resulting DE genes could be compared to the DE genes from mock vs. SARS-CoV-2 infection at higher viral titer (Fig 2). Given the exposure of cells to a lower viral titer, the number of DE genes from this comparison was smaller (4494 genes) vs. the comparison of high titer SARS-CoV-2 against mock (∼8000 genes) (Supp Fig S1C, https://doi.org/10.6084/m9.figshare.12272351 [Table S3]). Analysis of the 4494 DE genes showed significant enrichment in inflammation, autophagy and mitochondrial processes (Supp Fig S1D, https://doi.org/10.6084/m9.figshare.12272351 [Table S4]). To further assess the extent of overlap biological processes between low titer and high titer A549 cell datasets, the pathway enrichment results were graphically summarized and presented in a single map. This pathway summary map overlays the pathway enrichment results of mock vs. high titer SARS-CoV-2 infected cells on top of the mock vs. low virus titer infected cells to show processes exclusively (single color nodes) or commonly (double colored nodes) enriched between the two datasets. This analysis confirmed that perturbation in autophagy, inflammation, and mitochondrial processes were enriched by DE genes from both datasets (i.e. high and low MOI infected A549 cells) (Fig 1G).

#### Correlation of expression profiles between cell lines and lung cell types

The results presented in the section above and in subsequent sections used gene expression profiles of A549 (that were transduced with hACE2) and Calu3 lung cell lines. The biological significance of using these cell lines was assessed by evaluating the correlation of the gene expression profiles of cell lines with lung cells using the single cell (sc) RNA seq information from lung. First, using the scRNA seq data, cell type expression profile was computed as the mean expression across cells within each cell type. The top 1000 genes with the highest variance among the 57 cell type expression profiles were selected as highly variable genes which were presumably informative for differentiating the 57 cell types. Next, the expression profiles of lung cell lines were compared with the expression profile of (hACE2 transduced) A549 and Calu3 cell lines. This analysis revealed that the hACE2 transduced A549 cells gene expression strongly correlated with the basal and ionocyte lung cell subpopulations, which both represent lung epithelial cell lineage (34, 35). Correlation between the highly variable genes from lung scRNA seq data and either A549 or Calu3 cells were calculated and plotted (Supp Fig S1E). Notably, the Calu3 cells showed similar pattern but lower correlation with the lung cell types analyzed (Supp Fig S1E).

It is likely that the biological processes impacted in a SARS-CoV-2 infected A549 cells is likely impacted in the SARS-CoV-2 infected lung epithelial cells too. However, given the limitations of analyzing lung cell-lines data, gene expression analysis of lung samples from patient with severe or mild COVID-19 will help test if these processes are differently impacted depending on the severity of the disease. Together, these results support that SARS-CoV-2 infection impacts the expression of genes involved in the cytokine signaling, autophagy and mitochondria/respiration.

#### Comparing SARS-CoV-2 infection in hACE2 transduced A549 and Calu3 cell lines

A number of studies have been published that focused on identifying receptors used by SARS-CoV-2 to delineate viral entry mechanisms. Several of these studies have identified Angiotensin-Converting Enzyme 2 (ACE2) as the receptor that interacts with the SARS-CoV-2 spike protein to mediate viral entry (36, 37). Furthermore, TMPRSS2 and TMPRSS4, which are two membrane-bound serine proteases, were found to facilitate viral entry into the cells (24, 38, 39). Analysis of the gene expression datasets of A549 and Calu3 cells revealed that ACE2 and TMPRSS2 genes are highly expressed in the latter (https://doi.org/10.6084/m9.figshare.12272351 [Table S3]). To facilitate SARS-CoV-2 infection, A549 cells were transduced with hACE2 vector (28). The gene expression profile of SARS-CoV-2 infected A549 and Calu3 cells were compared and the correlation of gene expression between infected A549 and Calu3 cells was determined. We found significant correlation (R=0.68, p value<2.2e-16) between gene expression of SARS-CoV-2 infected A549 and Calu3 cells (Supp Fig S2A). Furthermore, 65% of DE genes from mock vs. SARS-CoV-2 in Calu3 comparison overlapped with DE genes from the respective A549 comparison (Supp Fig S2B). Finally, we plotted a pathway enrichment summary map by using the pathway enrichment results from mock vs. SARS-CoV-2 comparison in A549 and Calu3 cells (Supp Fig S2C). Overlaying the pathway analysis results from A549 over Calu3 revealed overlap of a wide range of biological processes including the interferon, neutrophils, mitochondrial, and autophagy processes between the two datasets (Supp Fig S2C). Together, these results show that, upon ACE2 expression, the gene expression changes in infected A549 cells is highly correlated with infected Calu3 cells.

### Network analysis identified protein-protein interaction subnetworks of genes involved in interferon, inflammation and mitochondrial translation

#### SARS-CoV-2 vs Mock: network analysis

To further understand the potential biological processes in play during SARS-CoV-2 infection, we performed a consensus weighted gene coexpression network analysis (WGCNA) (40) on combined, batch corrected (see methods), gene expression values of mock and SARS-CoV-2 infected at low and high titer cells, to identify clusters/modules of correlated gene. WGCNA identified more than 47 coexpression modules. The overlap of genes in each of these modules with significant DE genes from mock vs. SARS-CoV-2 infected at high titer is presented in the cluster dendrogram where each correlated module is represented by a color and their overlap with DE genes is shown in horizontal bars (Fig 3A, Supp Table 1).

**Figure 3:**
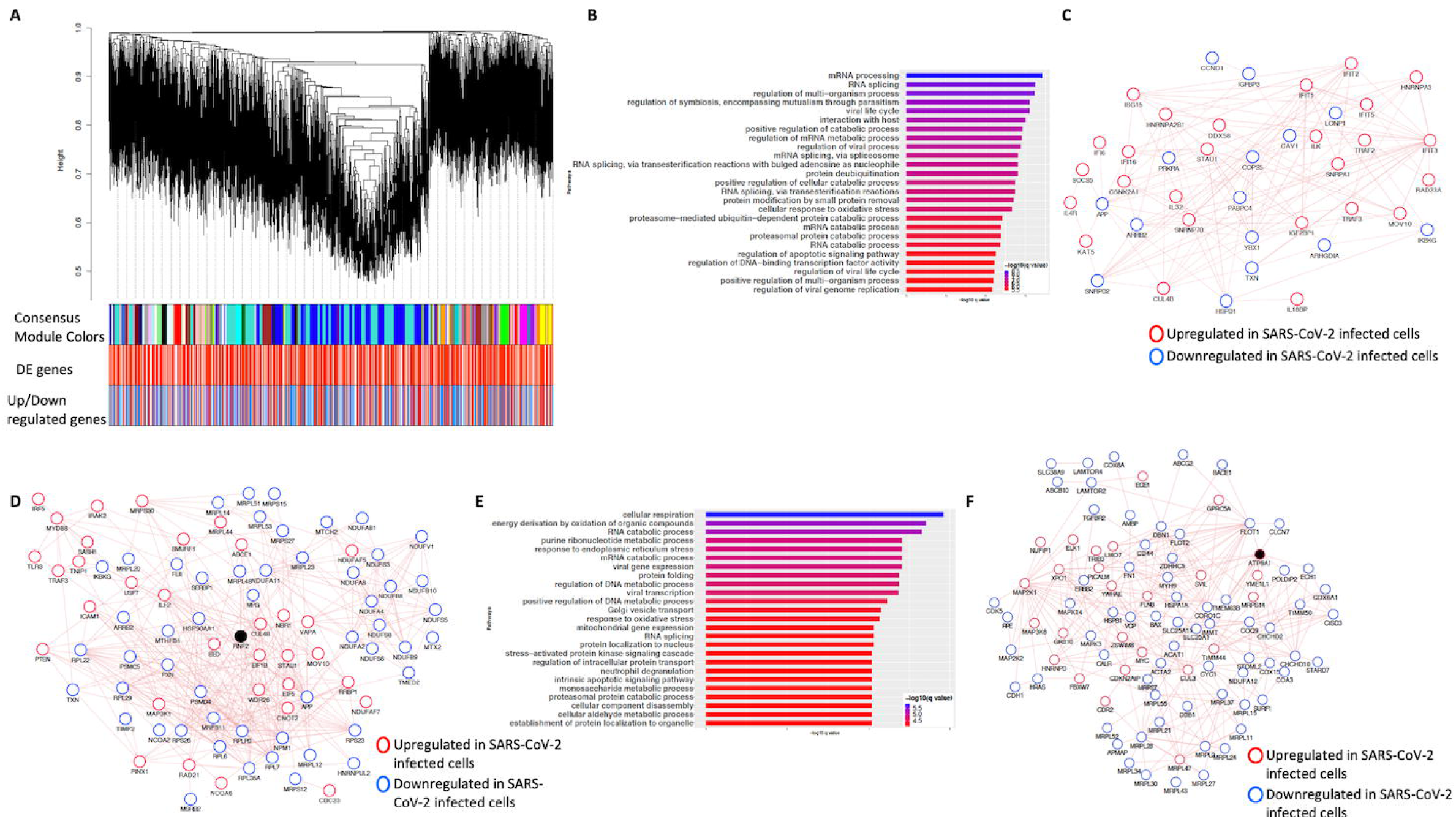
Consensus network analysis of hACE2 transduced mock and SARS-CoV-2 infected (high and low MOI) A549 cells. **(A)** Cluster dendrogram showing correlated genes grouped into clusters marked by different colors on the horizontal block labeled “Consensus Module Colors”. The DE genes in each cluster is marked as red color vertical line in the horizontal block labeled “DE genes”. The up and down regulated genes are shown as red and blue color vertical lines in a block labeled “Upregulated/downregulated genes”, respectively. **(B)** Top pathways from the pathway enrichment analysis of correlated DE genes in the blue module is presented as a horizontal bar plot, where x axis represents the −log10 transformed q value and the color of the horizontal bar is scaled blue to red representing low to high q values, respectively. **(C)** Protein-protein interaction (PPI) subnetworks in the blue module is presented where each node represents a gene and the border color of the nodes indicate up (red color) and downregulation (blue) in SARS-CoV-2 infected A549 cells (high MOI) compared to mock infected cells. The edge between the nodes indicate interaction based on the GeneMANIA database information. The network shows a highly connected interactome of interferon stimulated genes (ISGs) that are coordinately upregulated in infected cells. **(D)** Another PPI subnetwork identified in the blue module shows several highly interconnected mitochondrial ribosomal (MRP) and complex I (NDUF) genes. Each node represents a gene and the border color of the nodes indicate up (red color) and downregulation (blue) in SARS-CoV-2 infected A549 cells (high MOI) compared to mock infected cells. Additional interactome of MYD88 TRAF3 and TLR3 genes that were coordinately upregulated in infected cells can be seen (top left) **(E)** Top pathways from the pathway enrichment analysis of correlated DE genes in the turquoise module is presented as a horizontal bar plot, where x axis represents the −log10 transformed q value and the color of the horizontal bar is scaled blue to red representing low to high q values, respectively. (F) A PPI subnetwork of correlated DE genes in the turquoise module shows a well-connected interactome of genes encoding mitochondrial ribosomal proteins, mitochondrial coiled-coil-helix-coiled-coil-helix domain proteins and cytochrome oxidase. Each node represents a gene and the border color of the nodes indicate up (red color) and downregulation (blue) in SARS-CoV-2 infected cells (high MOI) compared to mock infected cells. DE: differentially expressed. MOI: multiplicity of infection.

First, pathway enrichment analysis of the correlated DE genes in the blue module showed significant annotation to the mitochondria, immunity, and mRNA/transcription processes (Fig 3B, https://doi.org/10.6084/m9.figshare.12272351 [Table S4]). Using the GeneMANIA (41) database, protein-protein interaction (PPI) subnetworks for the DE genes in this module/cluster were identified. This analysis identified two PPI subnetworks of genes involved in interferon signaling (Fig 3C), TLR3 and MYD88 (Fig 3D), and mitochondrial translation (Fig 3D). After incorporating the gene expression fold change information, we concluded that the interferon signaling and inflammation genes were upregulated, and mitochondrial genes were downregulated in SARS-CoV-2 infected cells.

Next, pathway enrichment analysis of DE genes from the turquoise module revealed significant annotation to viral gene expression and apoptosis processes (Fig 3E, https://doi.org/10.6084/m9.figshare.12272351 [Table S4]). Using GeneMANIA database, a PPI subnetwork of genes encoding the mitochondrial ribosomal proteins that were mostly downregulated in SARS-CoV-2 infected cells was also identified (Fig 3F). Together, these data suggest that SARS-CoV-2 infection results in a coordinated changed in the interferon signaling, inflammation, and mitochondrial processes.

### Gene expression changes associated with SARS-CoV-2 infection is distinct from Influenza A virus infection with minor overlaps

#### SARS-CoV-2 vs Influenza A virus (IAV)

To compare the expression profile of SARS-CoV-2 infected cells with another virus infected cells, DE analysis of mock vs. influenza A virus (IAV) infected cells was performed and the up and downregulated genes are presented in a volcano plot (Supp Fig S3A, https://doi.org/10.6084/m9.figshare.12272351 [Table S3]). The pathway analysis of the DE genes from this comparison showed enrichment in protein translation, localization and anti-viral responses (Supp Fig S3B, https://doi.org/10.6084/m9.figshare.12272351 [Table S4]). Additionally, genes upregulated in the IAV infected cells annotated to pathways for virus response, protein trafficking, and unfolded protein response (UPR) (Fig 4A, https://doi.org/10.6084/m9.figshare.12272351 [Table S4]). Genes that were downregulated in IAV infected cells enriched in vacuole and lysosome related processes (Fig 4B, https://doi.org/10.6084/m9.figshare.12272351 [Table S4]). Interestingly, few DE genes from the mock vs. SARS-CoV-2 overlapped with the DE genes from mock vs. IAV comparison (Supp Fig 3C).

**Figure 4:**
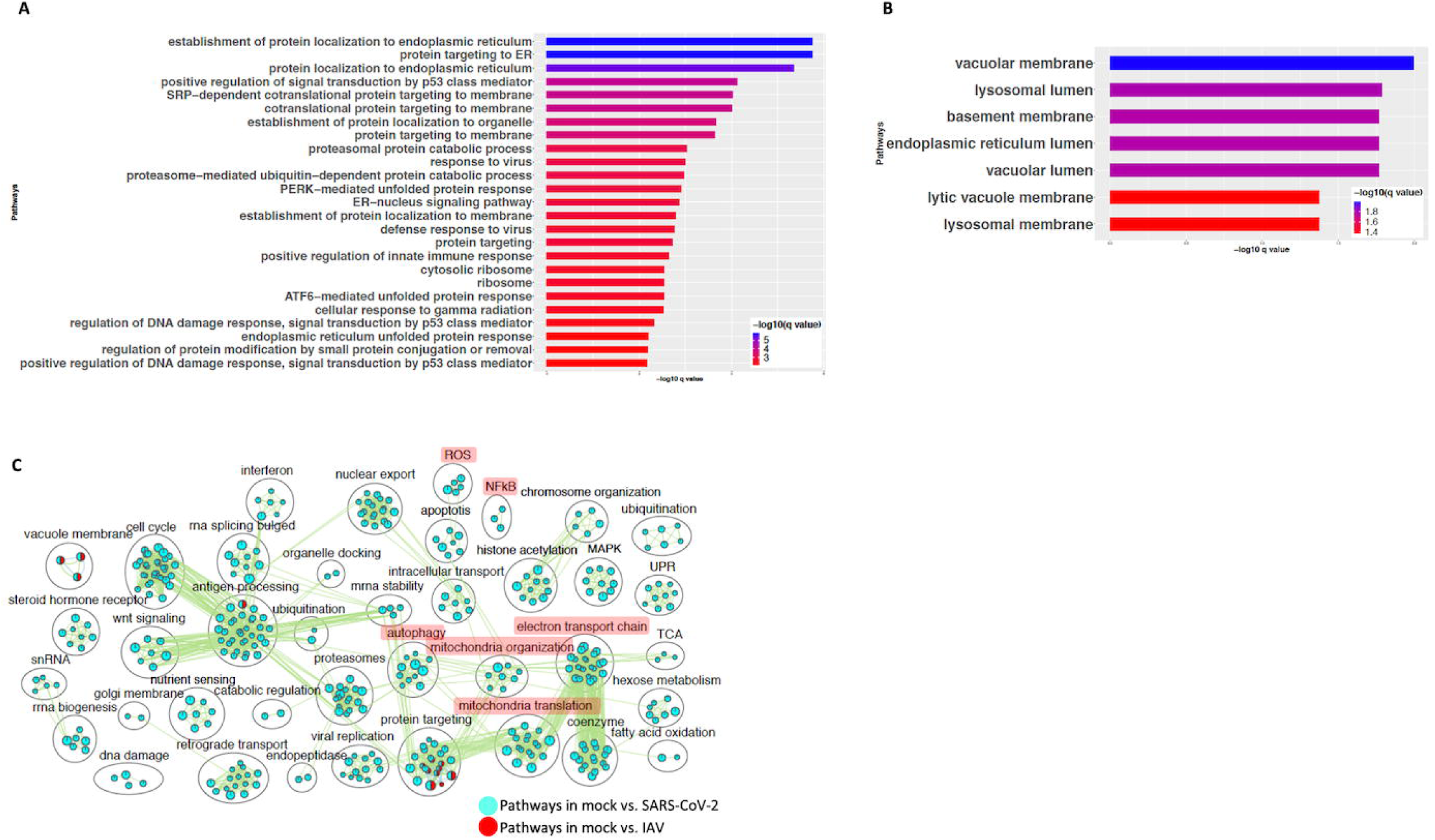
SARS-CoV-2 infection of A549 lung epithelial cells results in distinct gene expression changes that are not seen in IAV infection. **(A)** Top 25 pathways from the pathway enrichment analysis of DE genes upregulated in IAV infected cells is presented as a horizontal bar plot, where x axis represents the −log10 transformed q value and the color of the horizontal bar is scaled blue to red representing low to high q values, respectively. **(B)** Top 25 pathways from the pathway enrichment analysis of DE genes downregulated in IAV infected cells is presented as a horizontal bar plot, where x axis represents the −log10 transformed q value and the color of the horizontal bar is scaled blue to red representing low to high q values, respectively. **(C)** Pathway enrichment summary map for mock vs. SARS-CoV-2 (blue nodes) and mock vs. IAV (red nodes) comparisons. Each node represents a pathway/biological process (BP). The node size is proportional to the number of DE genes overlapping with the BP. The nodes that share genes are connected with edges. The label above each black circle summarizes the gene ontology (GO) terms of similar BPs present inside the circle. Single color nodes are pathways that are distinctly enriched by DE genes from one comparison. Two colored nodes are pathways enriched by DE genes from both comparisons. Notable groups of BPs associated with NFkB, ROS, autophagy and mitochondria are highlighted in red. DE genes from Mock vs. SARS-CoV-2 comparison exclusively enriched in highlighted BPs.

A pathway enrichment summary map was created by overlaying the pathway enrichment results of the mock vs. IAV comparison on top of the mock vs. SARS-CoV-2 comparison. Consistent with the DE genes comparison (Supp Fig 3C), the enrichment map also highlighted little overlap of pathways between the two comparisons. DE genes from both comparisons commonly enriched in a subset of pathways associated with protein trafficking (Fig 4C). Furthermore, only a subset of the interferon pathway genes and a few chemokine genes that were upregulated in SARS-CoV-2 infected cells were also upregulated in IAV infected cells, while the autophagy and inflammation genes remained mostly unchanged in the latter (Table S3 and S4). Therefore, upregulation of cytokine/inflammation, changes in autophagy, and downregulation of the mitochondrial processes were uniquely observed in SARS-CoV-2 infected cells. Upregulation of DE genes involved in the cytokine/inflammation processes is consistent with cytokine storm observed in severe cases COVID-19 patients. Since these observations were made by analyzing the gene expression changes in a lung cell line, future studies profiling gene expression changes in severe COVID-19 patients’ lung samples will be needed to confirm these findings.

### SARS-CoV-2 infected cells share some gene expression signature with MERS-CoV infected cells with few exceptions

#### SARS-CoV-2 vs MERS-CoV

Comparison of gene expression profiles revealed that a SARS-CoV-2 infected cells are distinct from those of IAV infected cells (Fig 4). Although these are both viruses, they are not phylogenetically close. Therefore, we next compared the gene expression profiles of SARS-CoV-2 and MERS-CoV infected cells, since both are hCoVs. DE analysis of the mock vs. SARS-CoV-2 infected Calu3 lung carcinoma cells identified several up and down genes (Supp Fig S4A, https://doi.org/10.6084/m9.figshare.12272351 [Table S3]). Pathway enrichment analysis showed annotation of the DE genes to cell cycle, inflammation, apoptosis processes (Supp Fig S4B, https://doi.org/10.6084/m9.figshare.12272351 [Table S4]). A pathway enrichment summary map for mock vs. SARS-CoV-2 and mock vs. MERS-CoV comparisons was generated to assess the extent of overlap of pathways between the two datasets (Fig 5A). Notably, the DE genes from both comparisons enriched in the mitochondria, autophagy, cell cycle, and UPR processes. However, DE genes from mock vs. SARS-CoV-2 comparison predominantly enriched in inflammation, cytokine signaling, and immunity related processes (Fig 5A). Consistently, genes upregulated in the SARS-CoV-2 infected Calu3 cells enriched in inflammation, nuclear factor kappaB (NFkB) processes (Fig 5B, https://doi.org/10.6084/m9.figshare.12272351 [Table S4]), while upregulated genes from both hCoV infected cells annotated to protein trafficking and small GTPase signaling (Fig 5B, 5C,, https://doi.org/10.6084/m9.figshare.12272351 [Table S4]). On the other hand, genes downregulated in both comparisons commonly annotated to mitochondrial processes (Fig 5D, 5E, https://doi.org/10.6084/m9.figshare.12272351 [Table S4]). These findings suggest that perturbation of autophagy, mitochondrial genes are common gene expression signatures associated with hCoVs infection, but the SARS-CoV-2 virus almost exclusively impacts the cytokine/inflammatory processes in the lung cells. It is likely that perturbation of mitochondrial processes and autophagy may lead to a dysfunctional immune response (30, 31). Further studies will be required to understand if and how these processes may contribute to inflammation during SARS-CoV-2 infection.

**Figure 5:**
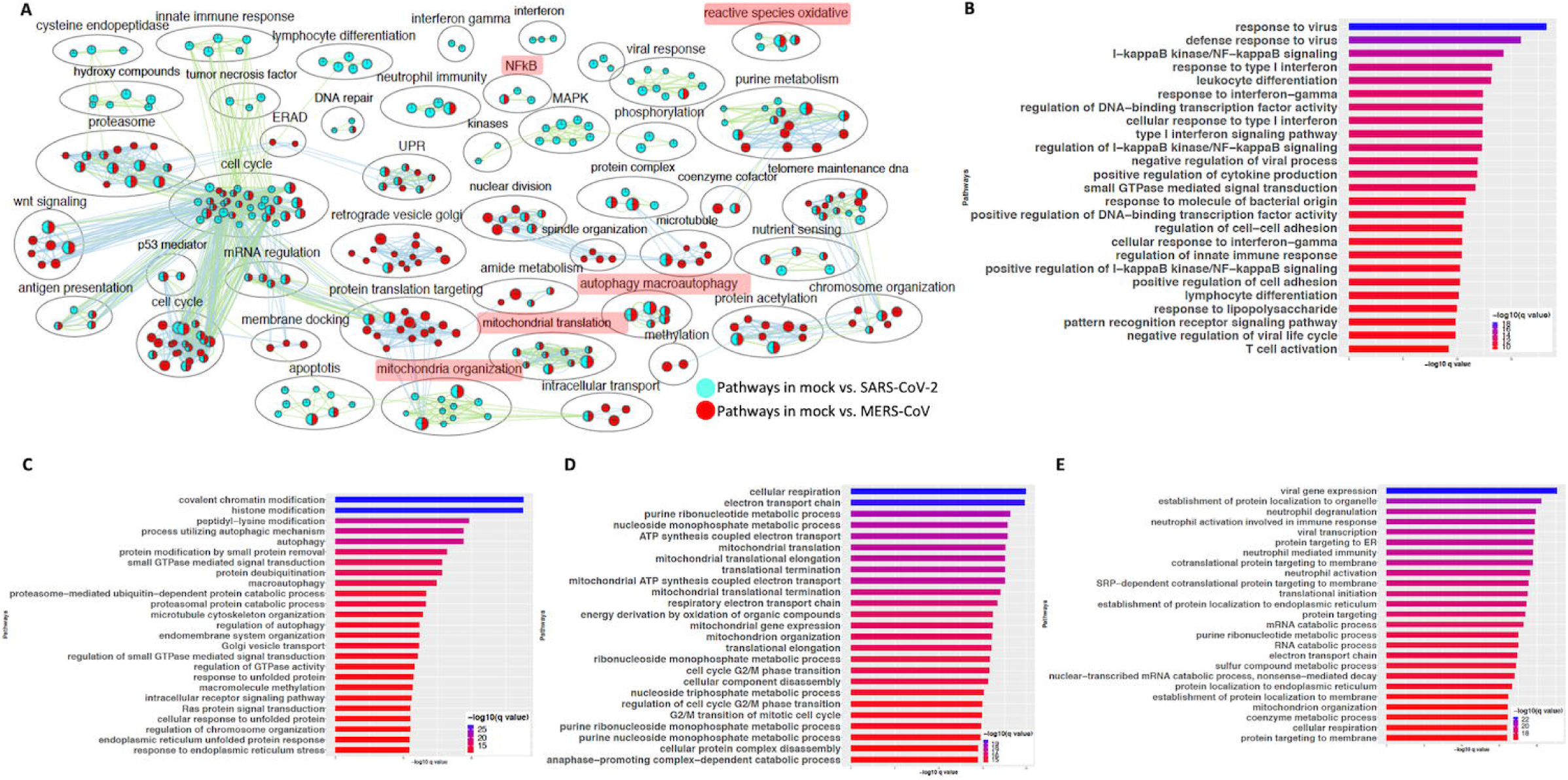
SARS-CoV-2 and MERS-CoV infection have some common and some distinct gene expression signatures. **(A)** Pathway enrichment summary map for mock vs. SARS-CoV-2 (blue nodes) and mock vs. MERS-CoV (red nodes) comparisons in Calu 3 cells. Each node represents a pathway/biological process (BP). The node size is proportional to the number of DE genes overlapping with the BP. The nodes that share genes are connected with edges. The label above each black circle summarizes the gene ontology (GO) terms of similar BPs present inside the circle. Single color nodes are pathways that are distinctly enriched by DE genes from one comparison. Two colored nodes are pathways enriched by DE genes from both comparisons. Notable groups of BPs associated with immunity, ROS, autophagy and mitochondria are highlighted in red. These notable groups of BPs were commonly enriched by DE genes from both SARS-CoV-2 and MERS-CoV comparisons. The DE genes from mock vs. SARS-CoV-2 comparison predominantly enriched in inflammation and immunity related processes. (B) Top 25 pathways from the pathway enrichment analysis of DE genes upregulated in SARS-CoV-2 infected Calu3 cells is presented as a horizontal bar plot, where x axis represents the −log10 transformed q value and the color of the horizontal bar is scaled blue to red representing low to high q values, respectively. (C) Top 25 pathways from the pathway enrichment analysis of DE genes upregulated in MERS-CoV infected Calu3 cells is presented as a horizontal bar plot, where x axis represents the −log10 transformed q value and the color of the horizontal bar is scaled blue to red representing low to high q values, respectively. (D) Top 25 pathways from the pathway enrichment analysis of DE genes downregulated in SARS-CoV-2 infected Calu3 cells is presented as a horizontal bar plot, where x axis represents the −log10 transformed q value and the color of the horizontal bar is scaled blue to red representing low to high q values, respectively. (E) Top 25 pathways from the pathway enrichment analysis of DE genes downregulated in MERS-CoV infected Calu3 cells is presented as a horizontal bar plot, where x axis represents the −log10 transformed q value and the color of the horizontal bar is scaled blue to red representing low to high q values, respectively. DE: differentially expressed.

### Validation of findings from cell line data in SARS-CoV-2 positive human samples

#### Comparing SARS-CoV-2 positive vs. negative human nasopharyngeal expression profile with A549 dataset

The gene expression profiling of the A549 and Calu3 cell line data revealed that perturbations in inflammatory, autophagy and mitochondrial processes were unique to coronavirus infections. To validate the findings from SARS-CoV-2 infected lung cell lines, we analyzed the gene expression profiles of SARS-CoV-2 positive human nasopharyngeal samples. DE analysis of SARS-CoV-2 positive vs. negative samples revealed up and downregulated genes (https://doi.org/10.6084/m9.figshare.12272351 [Table S3]). Concurrent with the cell line data, the pathway enrichment analysis of the DE genes revealed significant annotation to inflammation, autophagy and mitochondrial processes (Fig S5A, https://doi.org/10.6084/m9.figshare.12272351 [Table S4]). ∼60% of the DE genes from the SARS-CoV-2 infected human samples comparison overlapped with the infected A549 cells data (Fig S5B). Of this, ∼50% of the DE genes were concurrently downregulated in both datasets (i.e. human samples and A549 cells) and significantly annotated to the autophagy, immunity, and mitochondrial processes (Fig S5C). Since age (42) and viral load (43) can determine the severity of the COVID-19 outcome, the human samples were further classified into young (<40 years) and old (>60 years) with low or high viral load positive samples. Consistent with the SAR-CoV-2 positive vs. negative human data, the DE genes from the control vs. high viral load in old subjects significantly annotated to inflammation, autophagy, and mitochondrial processes (Fig 6A, https://doi.org/10.6084/m9.figshare.12272351 [Table S3 & S4]). When the pathway enrichment result from the old human subjects (that were high viral load positive) was overlaid on the pathway enrichment results from A549 cells transduced hACE2 infected with high MOI of SARS-CoV-2, autophagy, NFkB, oxidative stress, mitochondrial processes were commonly perturbed in both datasets (Fig 6B). Together, these observations suggest that across cell types, SARS-CoV-2 infection alters the cells’ autophagy and mitochondrial processes and that perturbations in these processes may impede an effective immune response leading to severe outcomes.

**Figure 6:**
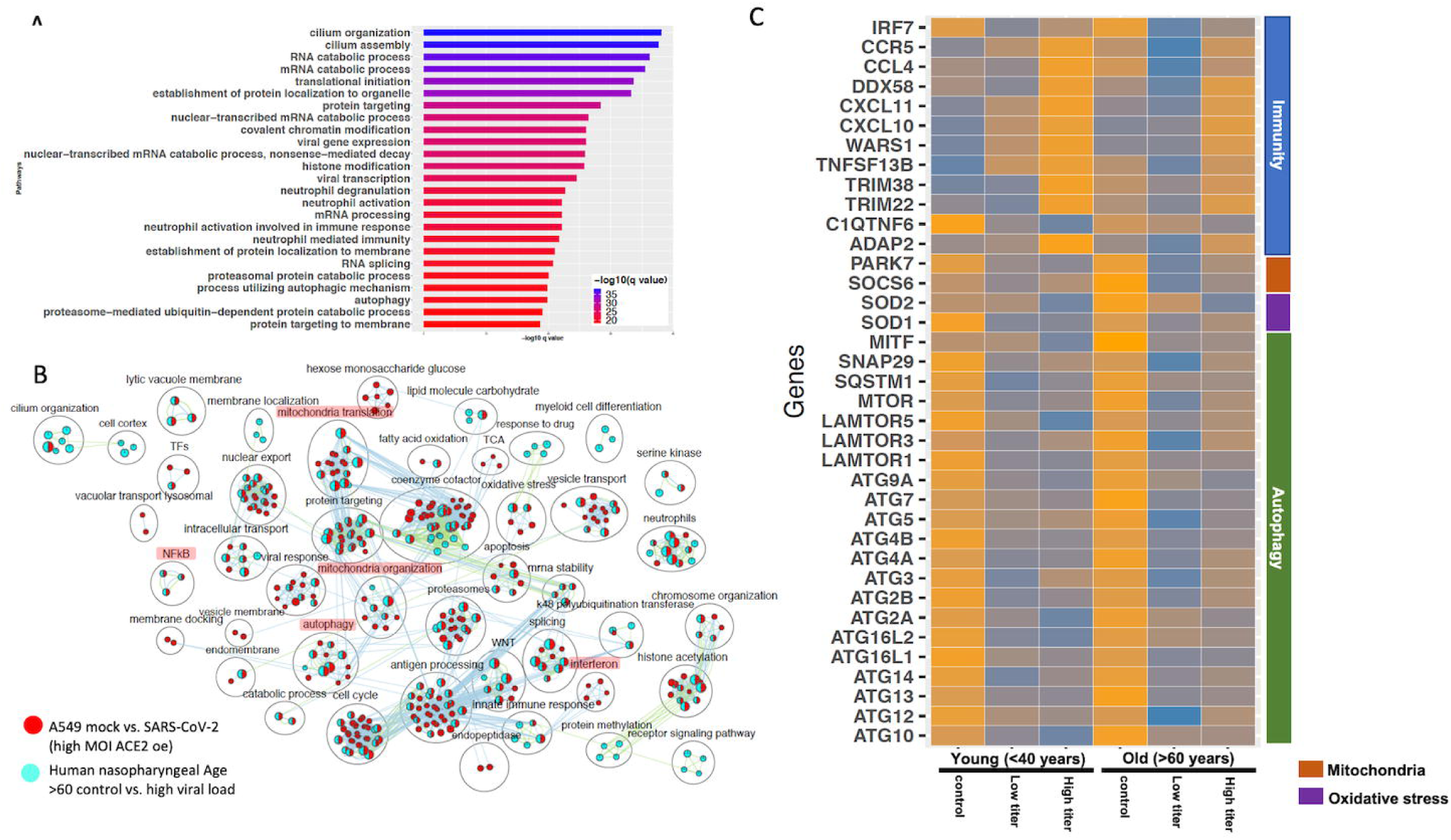
Gene expression profiling of human nasopharyngeal samples. (**A**) Top 25 pathways from the pathway enrichment analysis of DE genes from control vs. high viral load positive old age human samples is presented as a horizontal bar plot, where x axis represents the −log10 transformed q value and the color of the horizontal bar is scaled blue to red representing low to high q values, respectively. (**B**) Pathway enrichment summary map for control vs. high viral load positive human samples (blue nodes) and mock vs. SASR-CoV-2 (red nodes) comparisons in hACE2 transduced A549 cells. Each node represents a pathway/biological process (BP). The node size is proportional to the number of DE genes overlapping with the BP. The nodes that share genes are connected with edges. The label above each black circle summarizes the gene ontology (GO) terms of similar BPs present inside the circle. Single color nodes are pathways that are distinctly enriched by DE genes from one comparison. Two colored nodes are pathways enriched by DE genes from both comparisons. Notable groups of BPs associated with immunity, autophagy and mitochondria are highlighted in red. These notable groups of BPs were commonly enriched by DE genes from both comparisons. (**C**) Heatmap of the mean expression values of the indicated genes in young and old human samples that were negative (control) or positive with either high or low viral loads of SARS-CoV-2 virus. The indicated genes are broadly grouped into 4 different processes using a vertical bar present on the right side of the heatmap.

#### Delineating the age and viral load impact on SARS-CoV-2 response in human nasopharyngeal samples

The human nasopharyngeal samples were classified into young and old samples with low or high viral load to delineate either age specific or viral load specific gene expression changes in response to infection. The analysis in control vs. infected (with either low or high viral load) young or old samples helped identify DE genes that significantly annotated to autophagy, neutrophils, and mitochondrial processes (https://doi.org/10.6084/m9.figshare.12272351 [Table S3]), which is consistent with the pathway enrichment results from SARS-CoV-2 infected A549 cell line data (Fig 2). However in-depth gene expression analysis of the human data revealed some concurrent gene expression changes between cell line and human datasets, and some sample specific changes. Genes encoding the mTOR (Fig 6C), mitochondrial ribosomal genes, mitochondrial complex I genes, and lysosome acidification genes were downregulated in all of the SARS-CoV-2 positive human samples as well as in SARS-CoV-2 infected A549 cells https://doi.org/10.6084/m9.figshare.12272351 [Table S3]). Additionally, autophagy initiation, nucleation genes, p62 (SQSTM1), SNAP29, and MITF were specifically downregulated in SARS-CoV-2 positive human samples, which indicates decreased autophagic flux in infected samples (Fig 6C & Fig S6D). Furthermore, antioxidant encoding SOD1 (44) gene, the oxidative stress sensor PARK7 (DJ-1) (45), and mitochondria fission promoting SOCS6 (46) were also downregulated in infected human samples (Fig 6C). Finally, several cytokine/inflammation process genes were downregulated in infected human samples, which was opposite to the upregulation of these genes in SARS-CoV-2 infected A549 cells (Table S3).

There were several age and viral load dependent gene expression changes observed in the human nasopharyngeal samples, which are described below. Several interferon pathway genes were upregulated in high viral infected young and old samples. However, the upregulation of these gene in high viral load positive old samples was muted compared to the corresponding young samples (Fig S6A). The difference in interferon genes induction was most prominent in low viral load positive samples, as unlike the young samples where most of these genes were upregulated, in old samples, these interferon genes were significantly downregulated (Fig S6A). Similarly, ADAP2 (which is an interferon stimulated genes (ISG)), antiviral tripartite motif family E3 liage TRIM38, chemokine CCL4 and its receptor CCR5, and WARS1encoding the Tryptophanyl-tRNA Synthetase were all robustly down in old, low viral dose positive samples compared to the age matched control (Fig 6C). Together, these data suggest age-specific differences in host transcriptional response to SARS-CoV-2 infection. However, further analysis of the infected lung samples from old and young patients will be required to test if these differences may be causing more severe outcomes older patients.

### Gene expression analysis of a severe COVID-19 lung sample shows exaggerated immune/inflammation response

#### Healthy vs. COVID-19 lung biopsy samples

The gene expression analysis of the SAR-CoV-2 infected A549 and Calu3 cell lines revealed upregulation of the cytokine/inflammatory processes. However, this observation was inconsistent with the human nasopharyngeal expression profile where inflammatory process genes were downregulated. This difference could arise due to disparate cells type being compared, or that severe inflammatory symptoms may arise in severe case of COVID-19. Since the human nasopharyngeal data did not include clinical symptoms indicating the severity of COVID-19 in the samples whose sequence were analyzed, we could not stratify the samples based on disease severity. However, to test if the cytokine/inflammatory processes were also impacted in COVID-19 lungs, RNA seq data from the healthy and COVID-19 lung biopsy was analyzed. It should be noted that COVID-19 lung biopsy were technical replicates and therefore statistical significance of this analysis is limiting. Future studies involving bigger sample size will be required to confirm these observations. Nevertheless, the gene expression profile of a SARS-CoV-2 infected lung was distinct from healthy lungs with up and downregulated genes highlighted in the volcano plot (Supp Fig S7A, https://doi.org/10.6084/m9.figshare.12272351 [Table S3]). The pathway enrichment summary map and the plot showed that the DE genes predominantly annotated to the inflammation, ROS, leukocyte/monocyte related pathways (Fig S7B). Furthermore, the DE genes upregulated in the COVID-19 lungs enriched in the anti-viral response processes, cytokine secretion, immune cell proliferation/migration, and inflammation (Fig S7C, https://doi.org/10.6084/m9.figshare.12272351 [Table S4]). The downregulated genes were significantly enriched in protein trafficking, RNA metabolism, and oxygen sensing processes (Fig S7D, https://doi.org/10.6084/m9.figshare.12272351 [Table S4]). It is likely that perturbations in the oxygen sensing processes are reflective of the severe respiratory distress often seen in severe COVID-19 patients to due reduced oxygenation ability of the failing lungs.

## DISCUSSION

Highly pathogenic human coronaviruses (hCoV) are known to infect the lower respiratory airways and cause severe acute respiratory syndrome (SARS) (15). The recently discovered SARS-CoV-2 virus is the cause of COVID-19 (47). The clinical manifestations of this virus infection include fever, cough, fatigue, respiratory distress, and cardiac injury (48–50). While some patients with COVID-19 suffered from mild symptoms, other patients had increasingly life-threatening symptoms (50). Age and underlying medical conditions such as diabetes, hypertension, are likely to determine the severity of the symptoms (42). However, the underlying biological processes and mechanisms impacted by this viral infection of the host is still not clear. Analyzing the gene expression profiles of host cells infected with SARS-CoV-2 will be necessary to decipher the subcellular functions perturbed by this virus and to inform drug development strategies.

Here we present an in-depth differential expression analysis of A549 and Calu3 cell lines, comparing mock to infection with either SARS-CoV-2, or IAV, or MERS-CoV. Since A549 cells lacked expression of ACE2, TMPRSS2, or TMPRSS4 that are required for SARS-CoV-2 viral entry into the cells (24, 38, 39), A549 cells transduced with hACE2 were used. Upon SARS-CoV-2 infection at low and high MOI, viral transcripts were detected in these cells indicating infection (28). Concurrently, we also observed strong correlation between SARS-CoV-2 infected Calu3 and hACE2 transduced A549 cells. Furthermore, we report that (i) SARS-CoV-2 infection impacted the expression of genes in inflammation, cell cycle, reactive oxygen species (ROS), autophagy, and mitochondrial processes, which were absent in IAV infected cells; (ii) while perturbation in autophagy and mitochondrial processes is common in hCoV infections (SARS-CoV-2 and MERS-CoV), we found that increased expression of the inflammatory/cytokine signaling genes were exclusively observed in SARS-CoV-2 infected lung cells; (iii) Coexpression network analysis helped identify a cluster of genes involved in inflammation and mitochondrial translation process that were either coordinately up or downregulated in SARS-CoV-2 infected cells, respectively. Together, these data suggest that perturbation in the autophagy, mitochondrial processes in SARS-CoV-2 infected lung cells could hinder an effective immune response (30, 31) and increase inflammation, which is often seen in severe COVID-19 patients suffering from cytokine storm (15, 51). Since these conclusions were made using the data from virus infected lung cell lines, the correlation between these cells’ expression profiles and marker genes expression from different lung cell types were determined. While the A549 cells showed robust correlation with lung epithelial lineage basal and ionocyte cells, Calu3 cells showed similar pattern but lower correlation with these cell types. Therefore, the processes delineated in SARS-CoV-2 A549 cells likely represent the lung epithelial cells response to SARS-CoV-2 infection. To further substantiate the cell line findings, the gene expression profile of SARS-CoV-2 positive human nasopharyngeal samples were used for validation. Analysis of control vs. positive samples helped identify DE genes that significantly annotated to the autophagy, NFkB, oxidative stress, and mitochondrial processes. Using the patient information available from this dataset, the samples were grouped into young (<40 years) and old (>60 years) that were either positive with low or high viral load. Comparing the gene expression and pathway enrichment results of old age control vs. infected samples with A549 high MOI data revealed a wide range of BPs that were commonly perturbed in both datasets. Therefore, this analysis has delineated several biological processes, discussed in more detail below, that are impacted in the SARS-CoV-2 infected host cells.

Some complement genes (C1S, C1R) were specifically upregulated in high viral titer SARS-CoV-2 infected cell lines. Consistently, the C1q/TNF-related protein 6, a glycoprotein that regulates complement activation, was downregulated in both SARS-CoV-2 infected cells and human samples. This gene is implicated in arthritis, and intra-articular injection of the recombinant C1qTNF6 protein was shown as an effective strategy in improving arthritis and inflammation in C1qtnf6-/- mice (52). An elevated complement response could likely lead to excessive inflammation, which was also observed in MERS-CoV infection of the hDPP4-transgenic mouse model (53). Additionally, several past studies have highlighted the interplay between the complement and coagulation systems (54, 55). It is likely that the increased thrombosis in COVID-19 patients (56) is a result of excessive complement activation. Further assessment of complement activation in COVID-19 patients will be required to confirm this. Together, these observations suggest that inhibition of the complement system as potential treatment strategies could be tested.

Infection of A549 cells with SARS-CoV-2 at higher viral titer perturbed autophagy, upregulated genes in the interferon, cytokine, nuclear factor kappaB (NFkB), reactive oxygen species (ROS) processes, while downregulated genes in the mitochondrial, electron transport chain processes. Consistently, analysis of DE genes in one of the correlated clusters from WGCNA showed significant enrichment in the interferon signaling processes. Additionally, GeneMANIA analysis of the correlated DE genes in two modules revealed PPI subnetworks of genes involved in interferon stimulated genes (ISGs) and NFkB, which were both mostly upregulated in the infected cells. In addition to the ISGs, the JAK-STAT signal transduction genes, which play critical role in type I cytokine (such as IL6) signaling and inflammation (57–59), were also upregulated in the SARS-CoV-2 infected A549 cells (Fig 4C). IL6, a pleotropic cytokine, was shown to be elevated in critically ill COVID-19 patients (60). Consistently, IL6 was upregulated in the SARS-CoV-2 infected cells. IL6 acts via the JAK-STAT signaling through SOCS3 protein kinase (also upregulated in SARS-CoV-2 infected cells) to activate the immune response (61). Excessive IL6 causes excessive inflammation as seen in arthritis (62). However, the inflammatory/cytokine gene expression profile in the virus infected human nasopharyngeal samples was distinct from the cell line data. In the infected nasopharyngeal samples, most of the cytokine/inflammatory process genes were significantly downregulated compared to the control. This discordant inflammatory gene expression profiles between two datasets may be due to different cell types being compared (A549 is a lung cell line, and human nasopharyngeal samples are predominantly squamous epithelial cells). Or, the upregulation of cytokine/inflammatory processes genes in SARS-CoV-2 infected A549 cells may represent a severe COVID-19 infection state. Due to lack of clinical information describing the disease state of subjects whose nasopharyngeal samples were analyzed, we could not test this possibility in the nasopharyngeal dataset. However, analysis of the COVID-19 lung biopsy samples revealed significant upregulation of genes enriched in cytokine/inflammatory processes. Therefore, upregulation of IL6 and NFkB genes may be contribute to the inflammatory symptoms observed in severe COVID-19 patients (15, 51). These data support a central role for cytokine signaling in COVID-19 pathogenesis. Treatment strategies aimed at mitigating the cytokine effects or complement system could be tested in treatment of COVID-19. One such clinical trial aimed at mitigating the IL6 effects is already underway (NCT04322773). Furthermore, another study showed decreased mortality in patients treated with tocilizumab, which blocks IL6 (63).

Analysis of young (<40 years) and old (>60 years) nasopharyngeal samples also revealed some age-specific changes in the gene expression profile. Notably, the ISGs (IFIT1, IFIT2, and IFIT3) were upregulated in both low and high viral load SARS-CoV-2 positive young samples. Upregulation of these genes in high viral load infected old age samples were however less robust. Furthermore, in the low viral load positive old samples, most of the interferon genes (except IFIT3) were downregulated. In addition to the interferon genes, ADAP2, which is an ISG (64), TRIM5, which is a retroviral restriction factor (65), tripartite motif TRIM22 and TRIM38 were significantly downregulated in low viral load positive old samples compared to age matched control. The latter two genes are involved in innate immunity and in restricting viral infections (66, 67). Moreover, the Tryptophanyl-tRNA Synthetase encoding WARS1, which stimulates immunity against viral infection (68), chemokines CXCL11, CCL4 and CCL4 receptor CCR5 were also significantly downregulated in low viral load positive old samples compared to age matched control. Chemokine CXCL10 expression was unchanged in the low viral load positive old samples. All these chemokines and the receptor were upregulated in the low viral load positive young samples. It should be noted that upregulation of CXCL10, CXCL11, and IFIT2 in the nasopharyngeal samples has been proposed to accurately predict the presence of respiratory virus infection (69). Lack of induction of these genes in old age infected subjects may indicate an inability of the host cells to detect viral entry. This in combination with downregulation of genes involved in antiviral immunity likely contributes to severe disease outcomes, which is consistent with severe COVID-19 manifestations in older patients. However, more studies using the lung biopsy samples of SARS-CoV-2 infected young and old age patients will be required to confirm these findings.

What processes may be causing/contributing to defective immune response in the SARS-CoV-2 infected cells? Autophagy and mitochondria related processes were two other prominent categories of the biological processes that were exclusively impacted in the hCoV infected cells.

Gene expression analysis of SARS-CoV-2 infected A549 and Calu3 cells revealed perturbation of genes involved in autophagic processes. In contrast, most of the DE genes from human nasopharyngeal sample comparison that annotated to autophagy processes were downregulated. SARS-CoV-2 infected A549 and Calu3 cell lines and human nasopharyngeal samples expressed significantly low levels of mTOR and LAMTOR genes. The regulatory associated protein of mTOR (RAPTOR) expression was also significantly decreased in all SARS-CoV-2 infected samples. mTOR inhibits autophagy (70, 71). Consistent with the mTOR function the autophagy inducing microphthalamia-associated transcription factors (MiTF/TFE) and unc-51-like autophagy activating kinase 1 (ULK1) were upregulated in the infected A549 cells. Concurrent with this data, inhibition of mTORC1 in SARS-CoV-2 infected human bronchial epithelial cells NCI-H1299 and monkey kidney cells (VeroFM) has also been reported (72). These observations suggest that downregulation of mTOR may result in autophagy induction upon SARS-CoV-2 infection in A549 cells. However, contrary to the expression profiles of infected A549 cells, the autophagy initiation and nucleation genes were downregulated in infected human nasopharyngeal cells compared to control samples. It is worth noting that a past study in mouse embryonic cells showed that coronavirus infection induced autophagy and that coronavirus mouse hepatitis virus (MHV) replication was impaired in atg5-/- cells (73). While the gene expression data from SARS-CoV-2 infected cell lines and human samples indicate changes in autophagy genes expression, very little can be inferred about the changes to the autophagic flux, which is the capacity of the cells to degrade the autophagosome by fusing with the lysosomes. Studies have shown that Dengue and enteroviral infections inhibit autophagic flux in host cells by decreasing p62 or cleaving SNAP29, respectively, to facilitate infection (74, 75). Consistent with this observation, we found that the autophagic flux inducing p62 (SQSTM1) and SNAP29 genes expression level were down in SARS-CoV-2 positive human samples. Moreover, inhibition of S-phase kinase-associated protein 2 (SKP2) as E3 ligase decreased Beclin1 (BECN1) degradation, and increased autophagic flux, which in turn decreased MERS-CoV (76, 77). However, SKP2 expression in both SARS-CoV-2 infected cell lines and human samples was significantly low. Nevertheless, drugs known to increase autophagic flux has been shown to impede SARS-CoV-2 infection (72). It is likely that autophagic flux is decreased in human samples by downregulation of p62 and SNAP29, and in SARS-CoV-2 infected A549 cells potentially through upregulation of glycogen synthase kinase 3 beta (GSK3B), which has been shown to impair lysosome acidification (78). Consistently, GSK3B inhibition has been shown to increase autophagy flux in mice liver and human pancreatic cancer cells (79, 80). Additionally, several lysosome acidification genes were concordantly downregulated in both SARS-CoV-2 infected A549 cells and human samples. Since impaired lysosome acidification has been associated with impaired autophagic flux (81), drugs targeting lysosome reacidification or increasing autophagic flux could potentially be tested as a therapeutic intervention to SARS-CoV-2 infection. Since these inferences were made by analyzing the gene expression datasets, further studies will be required to assay the protein levels of genes involved in autophagy and the impact SARS-CoV-2 infection on autophagic flux in host cells.

In addition to changes in expression of genes involved the autophagic processes, several genes involved in the mitochondrial processes were downregulated in SARS-CoV-2 infected A549 cells and human samples. This is consistent with our current understanding that viruses either induce or inhibit various mitochondrial processes as part of their replication and dissemination efforts (82). Infection of cells with SARS-CoV-2 at higher viral titer downregulated the genes in the mitochondrial processes. Consistently, WGCNA of SARS-CoV-2 infected A549 datasets also identified a DE gene cluster that annotated to the mitochondrial organization and translation processes. Subsequent GeneMANIA analysis identified a PPI subnetwork of genes involved in mitochondrial translation which were coordinately downregulated in SARS-CoV-2 infected cells. Since mitochondrial import and translation are inter-linked (83, 84), we found that several mitochondrial complex I and translocase genes were downregulated in the SARS-CoV-2 infected cells. Given the extensive crosstalk between autophagy and mitochondrial function (85, 86), it is likely that perturbations in autophagy and mitochondrial processes observed in SARS-CoV-2 infected cells are interlinked. It is worth noting that mTORC1 complex which comprises the mTOR and RAPTOR stimulates synthesis of mitochondrial ribosomal, complex I proteins (32, 33) and mitochondrial fission process 1 (MTFP1) (87). Consistently, we found decreased expression mitochondrial ribosomal and complex I genes, which is likely a result of decreased mTOR and RAPTOR in SARS-CoV-2 infected cells. Additionally, decreased expression of MTFP1 that may impede mitochondrial fission resulting in hyperfused mitochondria (88) was specifically observed in SARS-CoV-2 infected A549 cells. While MTFP1 gene expression was undetectable in human nasopharyngeal samples, another mitochondrial fission promoting SOCS6 (46) was downregulated in infected samples. Past studies have shown that hyperfused or elongated mitochondria in Dengue and SARS infected cells can suppress interferon signaling and innate immunity (89–91). Furthermore, reduced complex I expression has been found in many cancer cells and is shown to affect the oxidative phosphorylation, which also impacts the immune cell function (31, 92–95). Together, these data highlight that SARS-CoV-2 infected cells have decreased mTOR, perturbation in autophagy and mitochondrial processes, which in turn could impede proper the immune response to infection. Further studies will however be required to delineate which of these perturbations are the direct result of SARS-CoV-2 infection or contribute to pathogenesis/severe clinical manifestations.

## SUMMARY

In summary, we have presented a detailed DE and coexpression network analysis of the RNA seq data from SARS-CoV-2 infected A549 cells. Using the gene expression profiles of A549 and Calu3 cells infected with IAV or MERS-CoV, we concluded that perturbations in cytokine signaling and inflammation processes, downregulation of genes in the mitochondrial processes, and perturbation of autophagy were uniquely observed in novel coronavirus infected cells. To validate the findings from the cell line data, gene expression analysis of control and SARS-CoV-2 positive human nasopharyngeal samples was performed. Consistent with the cell line data, DE genes from human data also significantly annotated to inflammation, autophagy, and mitochondrial processes. It is likely that perturbation of autophagy and mitochondrial processes may impede an effective immune response leading to severe outcomes. Furthermore, age stratified human nasopharyngeal was used to analyze gene expression changes in control vs. high viral load positive old age (>60 years) samples. This analysis also revealed several BPs that were concordantly impacted in both cell line and human datasets, with few differences. Specifically, gene encoding mTOR were downregulated in infected cells, which likely caused downregulation of mitochondrial ribosomal genes in both cell line and human datasets. Additionally, genes encoding mitochondrial complex I, lysosome acidification were also concurrently downregulated in infected cells from both datasets. Several inflammation process genes and autophagy genes were discordantly regulated in both datasets. It is likely that the autophagic flux is impeded in infected cells lines and human samples due to increased expression of GSK3B gene or downregulation of p62, SNAP29, respectively, which may further promote viral propagation. These data suggest that drugs that enhances autophagic flux or increases lysosome acidification could be tested as intervention strategies. Increased expression of inflammation process genes in A549 cells is likely due to these cells representing severe COVID-19 infection state. Consistently, we found upregulation of inflammatory process genes in expression profiles of COVID-19 lung biopsy sample. These data support a central role for cytokine/inflammation processes in COVID-19 pathogenesis. Finally, using the age stratified expression profiles of infected human nasopharyngeal samples, we identified muted or downregulation of several chemokine, ISGs, and tripartite motif genes that are critical for innate immunity and antiviral signaling. It is likely that defective antiviral response in old age patients in combination with perturbations in autophagy and mitochondrial processes could result in severe COVID-19 disease state often seen in older population. In summary, using gene expression data of SARS-CoV-2 infected cells, we show that viral infection of host cells results in perturbations in specific aspects of autophagy and mitochondrial processes. Future studies focusing on how these perturbations contribute either to viral propagation or impede an effective immune response will be required to gain more understanding of the viral pathogenesis.

## MATERIALS AND METHODS

### Data collection

Raw gene count matrix for bulk RNA-seq was obtained from GEO (accession number GSE147507) (28). The data contained gene expression count matrix of two lung carcinoma cell lines A549 (96, 97) and Calu3 (98, 99). In this dataset, the A549 treatment conditions included mock, infection with influenza A virus (IAV) (N=2 per group), and infection with SARS-CoV-2 at 2 (high titer, N=3 per group) and 0.2 (low titer, N=3 per group) multiplicity of infection (MOI), after transduction with a vector expressing human ACE2 (hACE2) (N=3 per group). The plasmid Ad-GFP-h-ACE2 from Vector Biolabs was used for transduction of A549 cells at 500 MOI (28). Subsequently, cells were infected with SARS-CoV-2 (Isolate USA-WA1/2020 (NR-52281)) at 0.2 or 2 MOI as indicated (28). From this dataset the raw gene count matrix for 2 healthy human lung biopsy and one COVID-19 samples (2 technical replicates), and Calu3 cells infected with SARS-CoV-2 at 2 MOI (N=3 per group) were also analyzed. Since the A549 cells infected with SARS-CoV-2 at high and low MOI were used in the network analysis, and DE genes from A549 and Calu3 comparisons were used, a boxplot of the normalized counts was created to show that all samples were comparable with no outlier (Supp Fig S8). Additionally, gene expression data in FPKM was downloaded from GEO (accession number GSE139516) (100) for Calu3 cell line infected with MERS-CoV and mock (N=3 per group). Differential expression (DE) analysis was performed on mock and MERS-CoV infected cells for 24 hr. Human lung single-cell RNA-seq (scRNA-seq) data with 57 annotated cell types was downloaded from Synapse (accession syn21041850) (101). For validation study, human nasopharyngeal gene expression matrix was obtained from GEO (accession number GSE152075) (29). Using the patient age, viral load information, the samples were further classified into young (<40 years) and old age (>60 years) that were either negative, or positive with low or high viral loads.

### Data analysis

#### RNA-seq analysis and network analysis

Differential expression analysis was performed using limma-voom and limma trends (102, 103). Genes with adjusted p value <0.05 were considered differentially expressed (DE). The DE genes were tested for pathway enrichment using clusterProfiler and pathways with q values (i.e. p values corrected for multiple comparison) < 0.05 were considered significant (104). To perform the consensus weighted gene coexpression network analysis (WGCNA) (40), first the pooled control and hACE2 transduced SARS-CoV-2 infected A549 cells at low and high titer expression data were batch corrected using combat (from the sva R package) (105), and the low expressing genes with count less than 5 in 4/6 samples were removed. The batch corrected normalized count were analyzed using the WGCNA R package (40) with default parameters. A total of 50 coexpressing modules were identified, of which, DE genes in correlated modules > 50 genes in size were selected for downstream analysis. For subnetwork analysis, GeneMANIA database (106) was used to identify potential protein-protein interaction (PPI) between the DE genes from the correlated modules. The PPI networks were then overlaid with the fold-change information using Cytoscape (107). Prior to DE analysis of the human nasopharyngeal gene expression data (29), the gene count matrix was subset to only include the SARS-CoV-2 positive and negative samples that were either <40 years (young) or >60 years (old) and exclude samples that were positive with medium load viral titer. Using the sva R package and batch information, the count matrix was batch corrected across three experimental groups: control (or negative), low viral load, and high viral load in both young and old groups. DE analysis of this dataset was performed as described above. DE analysis was also performed on batch corrected negative (control) vs. SARS-CoV-2 positive human nasopharyngeal samples.

#### Generating pathway enrichment summary map

The pathway enrichment summary map was generated using the indicated pathway enrichment results presented in supplemental table S4. To compare the pathway enrichment results from two different comparisons, the “Description”, “qvalue”, and “GeneID” information from each pathway enrichment table were used to compute similarity of the pathways and overlap of genes between the pathways. Using this information, the enrichment Map (108) app in Cytoscape creates a graphical network, where the nodes represent a pathway. If the genes annotated to pathway is shared with another pathway (which arises due to genes usually getting annotated to more than one pathway), an edge is drawn between the pathway nodes to show this information. Additionally, if DE genes from two different comparisons were significantly annotated to the same pathway/node, that node is highlighted with two colors. The common terms in the pathway description was then used to annotate/label a group of nodes using the Enrichment Map (108) and AutoAnnotate (109) apps in Cytoscape. R (Foundation for Statistical Computing, Vienna, Austria. URL https://www.R-project.org/) was used for data visualization.

The data sets supporting the results of this article are available from Figshare (https://doi.org/10.6084/m9.figshare.12272351) (110).

#### Single-cell RNA-seq analysis

Analysis of human lung single-cell RNA-seq (scRNA-seq) data with 57 annotated cell types was performed in R (v3.6) using Seurat (v3.1.1) (111). The UMI (Unique Molecular Identifier) count matrix was filtered for genes expressed in less than 3 cells and normalized using *SCTransform* implemented in Seurat. Differentially expressed genes were computed for 57 cell types using *FindAllMarkers* implemented in Seurat with default parameters. The UMAP plot was plotted using the top 50 principal components computed from the expression of highly variable genes selected by *SCTransform*.

## Acknowledgements

We thank Dr. Mark Knepper, head of the Epithelial Systems Biology Laboratory at the NHLBI, for providing his input in data analysis. This work utilized the computational resources of the NIH HPC Biowulf cluster (http://hpc.nih.gov). This work was supported by the Intramural Research Programs (IRPs) of the National Heart, Lung, and Blood Institute (NHLBI) grant number: 1ZICHL006228-03 to M.P.

## Supplemental Figure Legend

**Supp Fig S1**. (**A**) Volcano plot showing DE genes that were up (red color dots) and down regulated (blue color dots) in hACE2 transduced A549 cells infected with SARS-CoV-2 (high MOI). (**B**) Top 25 pathways from the pathway enrichment analysis of the DE genes from the mock vs. SARS-CoV-2 (high MOI) comparison is presented as a horizontal bar plot, where x axis represents the −log10 transformed q value and the color of the horizontal bar is scaled blue to red representing low to high q values, respectively. (**C**) Volcano plot showing DE genes that were up (red color dots) and down regulated (blue color dots) in SARS-CoV-2 (low MOI) infected A549 cells that were transduced with hACE2. (**D**) Top 25 pathways from the pathway enrichment analysis of the DE genes from the mock vs. SARS-CoV-2 (low MOI) comparison is presented as a horizontal bar plot, where x axis represents the −log10 transformed q value and the color of the horizontal bar is scaled blue to red representing low to high q values, respectively. DE: differentially expressed; MOI: multiplicity of infection. (**E**) Plot showing correlation between marker genes from different lung subpopulations (on x-axis) and hACE2 transduced A549 and Calu3 cells lines (color coded independent samples with legend at the bottom of the plot).

**Supp Fig S2**. (**A**) Correlation plot between mean gene expression from SARS-CoV-2 infected hACE2 transduced A549 (x axis) and Calu3 (y axis) cells. (**B**) Venn diagram showing overlap between DE genes from mock vs. SARS-CoV-2 A549 and Calu3 cell comparisons. (**C**) Pathway enrichment summary map for mock vs. SARS-CoV-2 comparisons in Calu3 (blue nodes) and hACE2 transduced A549 (red nodes) cells. Each node represents a pathway/biological process (BP). The node size is proportional to the number of DE genes overlapping with the BP. The nodes that share genes are connected with edges. The black circle outlines group the gene ontology (GO) terms of similar BPs. Single color nodes are pathways that are distinctly enriched by DE genes from one comparison. Two colored nodes are pathways enriched by DE genes from both comparisons. The DE genes from both comparisons enriched in inflammation, ROS, mitochondria and autophagy processes.

**Supp Fig S3**. (**A**) Volcano plot showing DE genes that were up (red color dots) and down regulated (blue color dots) in IAV infected A549 cells. (**B**) Top 25 pathways from the pathway enrichment analysis of the DE genes from the mock vs. IAV comparison is presented as a horizontal bar plot, where x axis represents the −log10 transformed q value and the color of the horizontal bar is scaled blue to red representing low to high q values, respectively. (**C**) Venn diagram showing DE genes overlap between mock vs. SARS-CoV-2 (High MOI) and mock vs. IAV comparisons. DE: differentially expressed; MOI: multiplicity of infection.

**Supp Fig S4**. (**A**) Volcano plot showing DE genes that were up (red color dots) and down regulated (blue color dots) in SARS-CoV-2 infected Calu3 cells. (**B**) Top 25 pathways from the pathway enrichment analysis of the DE genes from the mock vs. SARS-CoV-2 comparison in Calu3 is presented as a horizontal bar plot, where x axis represents the −log10 transformed q value and the color of the horizontal bar is scaled blue to red representing low to high q values, respectively.

**Supp Fig S5**. (**A**) Top 25 pathways from the pathway enrichment analysis of the DE genes from the positive (infected) vs negative human nasopharyngeal samples comparison is presented as a horizontal bar plot, where x axis represents the −log10 transformed q value and the color of the horizontal bar is scaled blue to red representing low to high q values, respectively. (**B**) Venn diagram showing DE genes overlap between control vs. high viral load old age human samples and mock vs. SARS-CoV-2 infected A549 cells comparisons. (**C**) Pathway enrichment result of common DE genes indicates in figure (B), that were concordantly downregulated in both datasets is presented as a horizontal bar plot, where x axis represents the −log10 transformed q value and the color of the horizontal bar is scaled blue to red representing low to high q values, respectively. DE: differentially expressed.

**Supp Fig S6**. Heatmap of the mean expression values of the indicated genes in young and old human samples that were negative (control) or positive with either high or low viral loads of SARS-CoV-2 virus is presentes. (**A**) Heatmap of interferon signaling genes. (**B**) Heatmap of mitochondrial ribosomal genes. (**C**) Heatmap of mitochondrial complex I genes. (**D**) Heatmap of lysosome acidification genes.

**Supp Fig S7: DE genes from COVID-19 lung compared to healthy lungs show robust upregulation of immunity, cytokines, and inflammatory processes.** (**A**) Volcano plot showing DE genes that were up (red color dots) and down regulated (blue color dots) in COVID-19 lung biopsy samples compared to healthy samples. (**B**) Pathway enrichment summary map for healthy vs. COVID-19 lungs (technical replicates) (blue nodes). Each node represents a pathway/biological process (BP). The node size is proportional to the number of DE genes overlapping with the BP. The nodes that share genes are connected with edges. The black circle summarizes the gene ontology (GO) terms of similar BPs. The DE genes from healthy vs. COVID-19 lung comparison predominantly enriched in inflammation and immunity related processes. (**C**) Pathway enrichment result of DE genes upregulated in COVID-19 lung vs. healthy lung biopsy samples is presented as a horizontal bar plot, where x axis represents the −log10 transformed q value and the color of the horizontal bar is scaled blue to red representing low to high q values, respectively. (**D**) Pathway enrichment result of DE genes downregulated in COVID-19 lung vs. healthy lung biopsy samples is presented as a horizontal bar plot, where x axis represents the −log10 transformed q value and the color of the horizontal bar is scaled blue to red representing low to high q values, respectively

## Supplemental Tables

**Supp Table 1:** Consensus module name and size (gene numbers) and its overlap with significant genes

**Supp Table 2:** List of marker genes of lung cell types from single cell RNA-seq data

## REFERENCES

1. Cascella M, Rajnik M, Cuomo A, Dulebohn SC, Di Napoli R. 2020. Features, Evaluation and Treatment Coronavirus (COVID-19), StatPearls, Treasure Island (FL).

2. Wang C, Horby PW, Hayden FG, Gao GF. 2020. A novel coronavirus outbreak of global health concern. Lancet 395:470–473.

3. Zhu N, Zhang D, Wang W, Li X, Yang B, Song J, Zhao X, Huang B, Shi W, Lu R, Niu P, Zhan F, Ma X, Wang D, Xu W, Wu G, Gao GF, Tan W, China Novel Coronavirus I, Research T. 2020. A Novel Coronavirus from Patients with Pneumonia in China, 2019. N Engl J Med 382:727-733.

4. Chan JF, Yuan S, Kok KH, To KK, Chu H, Yang J, Xing F, Liu J, Yip CC, Poon RW, Tsoi HW, Lo SK, Chan KH, Poon VK, Chan WM, Ip JD, Cai JP, Cheng VC, Chen H, Hui CK, Yuen KY. 2020. A familial cluster of pneumonia associated with the 2019 novel coronavirus indicating person-to-person transmission: a study of a family cluster. Lancet 395:514–523.

5. Fung SY, Yuen KS, Ye ZW, Chan CP, Jin DY. 2020. A tug-of-war between severe acute respiratory syndrome coronavirus 2 and host antiviral defence: lessons from other pathogenic viruses. Emerg Microbes Infect 9:558–570.

6. Yuen KS, Ye ZW, Fung SY, Chan CP, Jin DY. 2020. SARS-CoV-2 and COVID-19: The most important research questions. Cell Biosci 10:40.

7. World Health Organization. 2020. Coronavirus disease (COVID-19) Pandemic. https://www.who.int/emergencies/diseases/novel-coronavirus-2019. Accessed April 27, 2020.

8. Raoult D, Zumla A, Locatelli F, Ippolito G, Kroemer G. 2020. Coronavirus infections: Epidemiological, clinical and immunological features and hypotheses. Cell Stress 4:66–75.

9. de Wit E, van Doremalen N, Falzarano D, Munster VJ. 2016. SARS and MERS: recent insights into emerging coronaviruses. Nat Rev Microbiol 14:523–34.

10. Wu A, Peng Y, Huang B, Ding X, Wang X, Niu P, Meng J, Zhu Z, Zhang Z, Wang J, Sheng J, Quan L, Xia Z, Tan W, Cheng G, Jiang T. 2020. Genome Composition and Divergence of the Novel Coronavirus (2019-nCoV) Originating in China. Cell Host Microbe 27:325–328.

11. Zhang YZ, Holmes EC. 2020. A Genomic Perspective on the Origin and Emergence of SARS-CoV-2. Cell 181:223–227.

12. Katze MG, Fornek JL, Palermo RE, Walters KA, Korth MJ. 2008. Innate immune modulation by RNA viruses: emerging insights from functional genomics. Nat Rev Immunol 8:644–54.

13. Blanco-Melo D, Nilsson-Payant BE, Liu W-C, Møller R, Panis M, Sachs D, Albrecht RA, tenOever BR. 2020. SARS-CoV-2 launches a unique transcriptional signature from in vitro, ex vivo, and in vivo systems. bioRxiv.

14. Grove J, Marsh M. 2011. The cell biology of receptor-mediated virus entry. J Cell Biol 195:1071–82.

15. Channappanavar R, Perlman S. 2017. Pathogenic human coronavirus infections: causes and consequences of cytokine storm and immunopathology. Semin Immunopathol 39:529–539.

16. Liao M, Liu Y, Yuan J, Wen Y, Xu G, Zhao J, Chen L, Li J, Wang X, Wang F, Liu L, Zhang S, Zhang Z. 2020. The landscape of lung bronchoalveolar immune cells in COVID-19 revealed by single-cell RNA sequencing. medRxiv.

17. Mar KB, Rinkenberger NR, Boys IN, Eitson JL, McDougal MB, Richardson RB, Schoggins JW. 2018. LY6E mediates an evolutionarily conserved enhancement of virus infection by targeting a late entry step. Nat Commun 9:3603.

18. Chen G, Wu D, Guo W, Cao Y, Huang D, Wang H, Wang T, Zhang X, Chen H, Yu H, Zhang X, Zhang M, Wu S, Song J, Chen T, Han M, Li S, Luo X, Zhao J, Ning Q. 2020. Clinical and immunological features of severe and moderate coronavirus disease 2019. J Clin Invest doi:10.1172/JCI137244.

19. Rockx B, Kuiken T, Herfst S, Bestebroer T, Lamers MM, de Meulder D, van Amerongen G, van den Brand J, Okba NMA, Schipper D, van Run P, Leijten L, Verschoor E, Verstrepen B, Langermans J, Drosten C, van Vlissingen MF, Fouchier R, de Swart R, Koopmans M, Haagmans BL. 2020. Comparative Pathogenesis Of COVID-19, MERS And SARS In A Non-Human Primate Model. bioRxiv.

20. To KF, Tong JH, Chan PK, Au FW, Chim SS, Chan KC, Cheung JL, Liu EY, Tse GM, Lo AW, Lo YM, Ng HK. 2004. Tissue and cellular tropism of the coronavirus associated with severe acute respiratory syndrome: an in-situ hybridization study of fatal cases. J Pathol 202:157–63.

21. Chu H, Chan JF-W, Yuen TT-T, Shuai H, Yuan S, Wang Y, Hu B, Yip CC-Y, Tsang JO-L, Huang X, Chai Y, Yang D, Hou Y, Chik KK-H, Zhang X, Fung AY-F, Tsoi H-W, Cai J-P, Chan W-M, Ip JD, Chu AW-H, Zhou J, Lung DC, Kok K-H, To KK-W, Tsang OT-Y, Chan K-H, Yuen K-Y. 2020. Comparative tropism, replication kinetics, and cell damage profiling of SARS-CoV-2 and SARS-CoV with implications for clinical manifestations, transmissibility, and laboratory studies of COVID-19: an observational study. The Lancet Microbe doi:https://doi.org/10.1016/S2666-5247(20)30004-5.

22. Ziegler CGK, Allon SJ, Nyquist SK, Mbano IM, Miao VN, Tzouanas CN, Cao Y, Yousif AS, Bals J, Hauser BM, Feldman J, Muus C, Wadsworth MH, Kazer SW, Hughes TK, Doran B, Gatter GJ, Vukovic M, Taliaferro F, Mead BE, Guo Z, Wang JP, Gras D, Plaisant M, Ansari M, Angelidis I, Adler H, Sucre JMS, Taylor CJ, Lin B, Waghray A, Mitsialis V, Dwyer DF, Buchheit KM, Boyce JA, Barrett NA, Laidlaw TM, Carroll SL, Colonna L, Tkachev V, Peterson CW, Yu A, Zheng HB, Gideon HP, Winchell CG, Lin PL, Bingle CD, Snapper SB, Kropski JA, Theis FJ, et al. 2020. SARS-CoV-2 receptor ACE2 is an interferon-stimulated gene in human airway epithelial cells and is detected in specific cell subsets across tissues. Cell doi:https://doi.org/10.1016/j.cell.2020.04.035.

23. Walls AC, Park YJ, Tortorici MA, Wall A, McGuire AT, Veesler D. 2020. Structure, Function, and Antigenicity of the SARS-CoV-2 Spike Glycoprotein. Cell 181:281-292 e6.

24. Hoffmann M, Kleine-Weber H, Schroeder S, Kruger N, Herrler T, Erichsen S, Schiergens TS, Herrler G, Wu NH, Nitsche A, Muller MA, Drosten C, Pohlmann S. 2020. SARS-CoV-2 Cell Entry Depends on ACE2 and TMPRSS2 and Is Blocked by a Clinically Proven Protease Inhibitor. Cell 181:271–280 e8.

25. Yan R, Zhang Y, Li Y, Xia L, Guo Y, Zhou Q. 2020. Structural basis for the recognition of SARS-CoV-2 by full-length human ACE2. Science 367:1444–1448.

26. Zhou L, Niu Z, Jiang X, Zhang Z, Zheng Y, Wang Z, Zhu Y, Gao L, Wang X, Sun Q. 2020. Systemic analysis of tissue cells potentially vulnerable to SARS-CoV-2 infection by the protein-proofed single-cell RNA profiling of ACE2, TMPRSS2 and Furin proteases. bioRxiv doi:10.1101/2020.04.06.028522:2020.04.06.028522.

27. Daly JL, Simonetti B, Antón-Plágaro C, Kavanagh Williamson M, Shoemark DK, Simón-Gracia L, Klein K, Bauer M, Hollandi R, Greber UF, Horvath P, Sessions RB, Helenius A, Hiscox JA, Teesalu T, Matthews DA, Davidson AD, Cullen PJ, Yamauchi Y. 2020. Neuropilin-1 is a host factor for SARS-CoV-2 infection. bioRxiv doi:10.1101/2020.06.05.134114:2020.06.05.134114.

28. Blanco-Melo D, Nilsson-Payant BE, Liu WC, Uhl S, Hoagland D, Moller R, Jordan TX, Oishi K, Panis M, Sachs D, Wang TT, Schwartz RE, Lim JK, Albrecht RA, tenOever BR. 2020. Imbalanced Host Response to SARS-CoV-2 Drives Development of COVID-19. Cell 181:1036–1045 e9.

29. Lieberman NAP, Peddu V, Xie H, Shrestha L, Huang ML, Mears MC, Cajimat MN, Bente DA, Shi PY, Bovier F, Roychoudhury P, Jerome KR, Moscona A, Porotto M, Greninger AL. 2020. In vivo antiviral host response to SARS-CoV-2 by viral load, sex, and age. bioRxiv doi:10.1101/2020.06.22.165225.

30. Jang YJ, Kim JH, Byun S. 2019. Modulation of Autophagy for Controlling Immunity. Cells 8.

31. Won JH, Park S, Hong S, Son S, Yu JW. 2015. Rotenone-induced Impairment of Mitochondrial Electron Transport Chain Confers a Selective Priming Signal for NLRP3 Inflammasome Activation. J Biol Chem 290:27425–37.

32. Morita M, Gravel SP, Chenard V, Sikstrom K, Zheng L, Alain T, Gandin V, Avizonis D, Arguello M, Zakaria C, McLaughlan S, Nouet Y, Pause A, Pollak M, Gottlieb E, Larsson O, St-Pierre J, Topisirovic I, Sonenberg N. 2013. mTORC1 controls mitochondrial activity and biogenesis through 4E-BP-dependent translational regulation. Cell Metab 18:698–711.

33. Morita M, Gravel SP, Hulea L, Larsson O, Pollak M, St-Pierre J, Topisirovic I. 2015. mTOR coordinates protein synthesis, mitochondrial activity and proliferation. Cell Cycle 14:473–80.

34. Schiller HB, Montoro DT, Simon LM, Rawlins EL, Meyer KB, Strunz M, Vieira Braga FA, Timens W, Koppelman GH, Budinger GRS, Burgess JK, Waghray A, van den Berge M, Theis FJ, Regev A, Kaminski N, Rajagopal J, Teichmann SA, Misharin AV, Nawijn MC. 2019. The Human Lung Cell Atlas: A High-Resolution Reference Map of the Human Lung in Health and Disease. Am J Respir Cell Mol Biol 61:31–41.

35. Morrisey EE. 2018. Basal Cells in Lung Development and Repair. Dev Cell 44:653–654.

36. Shang J, Ye G, Shi K, Wan Y, Luo C, Aihara H, Geng Q, Auerbach A, Li F. 2020. Structural basis of receptor recognition by SARS-CoV-2. Nature 581:221-224.

37. Li W, Sui J, Huang IC, Kuhn JH, Radoshitzky SR, Marasco WA, Choe H, Farzan M. 2007. The S proteins of human coronavirus NL63 and severe acute respiratory syndrome coronavirus bind overlapping regions of ACE2. Virology 367:367–74.

38. Zang R, Gomez Castro MF, McCune BT, Zeng Q, Rothlauf PW, Sonnek NM, Liu Z, Brulois KF, Wang X, Greenberg HB, Diamond MS, Ciorba MA, Whelan SPJ, Ding S. 2020. TMPRSS2 and TMPRSS4 promote SARS-CoV-2 infection of human small intestinal enterocytes. Sci Immunol 5.

39. Iwata-Yoshikawa N, Okamura T, Shimizu Y, Hasegawa H, Takeda M, Nagata N. 2019. TMPRSS2 Contributes to Virus Spread and Immunopathology in the Airways of Murine Models after Coronavirus Infection. J Virol 93.

40. Langfelder P, Horvath S. 2008. WGCNA: an R package for weighted correlation network analysis. BMC Bioinformatics 9:559.

41. Franz M, Rodriguez H, Lopes C, Zuberi K, Montojo J, Bader GD, Morris Q. 2018. GeneMANIA update 2018. Nucleic Acids Res 46:W60–W64.

42. Wang D, Hu B, Hu C, Zhu F, Liu X, Zhang J, Wang B, Xiang H, Cheng Z, Xiong Y, Zhao Y, Li Y, Wang X, Peng Z. 2020. Clinical Characteristics of 138 Hospitalized Patients With 2019 Novel Coronavirus-Infected Pneumonia in Wuhan, China. JAMA doi:10.1001/jama.2020.1585.

43. Liu Y, Yan LM, Wan L, Xiang TX, Le A, Liu JM, Peiris M, Poon LLM, Zhang W. 2020. Viral dynamics in mild and severe cases of COVID-19. Lancet Infect Dis 20:656–657.

44. Fukai T, Ushio-Fukai M. 2011. Superoxide dismutases: role in redox signaling, vascular function, and diseases. Antioxid Redox Signal 15:1583–606.

45. Wang B, Cai Z, Tao K, Zeng W, Lu F, Yang R, Feng D, Gao G, Yang Q. 2016. Essential control of mitochondrial morphology and function by chaperone-mediated autophagy through degradation of PARK7. Autophagy 12:1215–28.

46. Lin HY, Lai RH, Lin ST, Lin RC, Wang MJ, Lin CC, Lee HC, Wang FF, Chen JY. 2013. Suppressor of cytokine signaling 6 (SOCS6) promotes mitochondrial fission via regulating DRP1 translocation. Cell Death Differ 20:139–53.

47. Tammaro A, Adebanjo GAR, Parisella FR, Pezzuto A, Rello J. 2020. Cutaneous manifestations in COVID-19: the experiences of Barcelona and Rome. J Eur Acad Dermatol Venereol doi:10.1111/jdv.16530.

48. Chen N, Zhou M, Dong X, Qu J, Gong F, Han Y, Qiu Y, Wang J, Liu Y, Wei Y, Xia J, Yu T, Zhang X, Zhang L. 2020. Epidemiological and clinical characteristics of 99 cases of 2019 novel coronavirus pneumonia in Wuhan, China: a descriptive study. Lancet 395:507-513.

49. Huang C, Wang Y, Li X, Ren L, Zhao J, Hu Y, Zhang L, Fan G, Xu J, Gu X, Cheng Z, Yu T, Xia J, Wei Y, Wu W, Xie X, Yin W, Li H, Liu M, Xiao Y, Gao H, Guo L, Xie J, Wang G, Jiang R, Gao Z, Jin Q, Wang J, Cao B. 2020. Clinical features of patients infected with 2019 novel coronavirus in Wuhan, China. Lancet 395:497–506.

50. Guan W-j, Ni Z-y, Hu Y, Liang W-h, Ou C-q, He J-x, Liu L, Shan H, Lei C-l, Hui DSC, Du B, Li L-j, Zeng G, Yuen K-Y, Chen R-c, Tang C-l, Wang T, Chen P-y, Xiang J, Li S-y, Wang J-l, Liang Z-j, Peng Y-x, Wei L, Liu Y, Hu Y-h, Peng P, Wang J-m, Liu J-y, Chen Z, Li G, Zheng Z-j, Qiu S-q, Luo J, Ye C-j, Zhu S-y, Zhong N-s. 2020. Clinical Characteristics of Coronavirus Disease 2019 in China. New England Journal of Medicine doi:10.1056/NEJMoa2002032.

51. Mehta P, McAuley DF, Brown M, Sanchez E, Tattersall RS, Manson JJ, Hlh Across Speciality Collaboration UK. 2020. COVID-19: consider cytokine storm syndromes and immunosuppression. Lancet 395:1033-1034.

52. Murayama MA, Kakuta S, Inoue A, Umeda N, Yonezawa T, Maruhashi T, Tateishi K, Ishigame H, Yabe R, Ikeda S, Seno A, Chi HH, Hashiguchi Y, Kurata R, Tada T, Kubo S, Sato N, Liu Y, Hattori M, Saijo S, Matsushita M, Fujita T, Sumida T, Iwakura Y. 2015. CTRP6 is an endogenous complement regulator that can effectively treat induced arthritis. Nat Commun 6:8483.

53. Jiang Y, Zhao G, Song N, Li P, Chen Y, Guo Y, Li J, Du L, Jiang S, Guo R, Sun S, Zhou Y. 2018. Blockade of the C5a-C5aR axis alleviates lung damage in hDPP4-transgenic mice infected with MERS-CoV. Emerg Microbes Infect 7:77.

54. Oikonomopoulou K, Ricklin D, Ward PA, Lambris JD. 2012. Interactions between coagulation and complement--their role in inflammation. Semin Immunopathol 34:151–65.

55. Skoglund C, Wettero J, Tengvall P, Bengtsson T. 2010. C1q induces a rapid up-regulation of P-selectin and modulates collagen-and collagen-related peptide-triggered activation in human platelets. Immunobiology 215:987–95.

56. Bikdeli B, Madhavan MV, Jimenez D, Chuich T, Dreyfus I, Driggin E, Nigoghossian CD, Ageno W, Madjid M, Guo Y, Tang LV, Hu Y, Giri J, Cushman M, Quéré I, Dimakakos EP, Gibson CM, Lippi G, Favaloro EJ, Fareed J, Caprini JA, Tafur AJ, Burton JR, Francese DP, Wang EY, Falanga A, McLintock C, Hunt BJ, Spyropoulos AC, Barnes GD, Eikelboom JW, Weinberg I, Schulman S, Carrier M, Piazza G, Beckman JA, Steg PG, Stone GW, Rosenkranz S, Goldhaber SZ, Parikh SA, Monreal M, Krumholz HM, Konstantinides SV, Weitz JI, Lip GYH. 2020. COVID-19 and Thrombotic or Thromboembolic Disease: Implications for Prevention, Antithrombotic Therapy, and Follow-up. Journal of the American College of Cardiology doi:10.1016/j.jacc.2020.04.031:27284.

57. O’Brown ZK, Van Nostrand EL, Higgins JP, Kim SK. 2015. The Inflammatory Transcription Factors NFkappaB, STAT1 and STAT3 Drive Age-Associated Transcriptional Changes in the Human Kidney. PLoS Genet 11:e1005734.

58. Leonard WJ, O’Shea JJ. 1998. Jaks and STATs: biological implications. Annu Rev Immunol 16:293–322.

59. Banerjee S, Biehl A, Gadina M, Hasni S, Schwartz DM. 2017. JAK-STAT Signaling as a Target for Inflammatory and Autoimmune Diseases: Current and Future Prospects. Drugs 77:521–546.

60. Chen X, Zhao B, Qu Y, Chen Y, Xiong J, Feng Y, Men D, Huang Q, Liu Y, Yang B, Ding J, Li F. 2020. Detectable serum SARS-CoV-2 viral load (RNAaemia) is closely associated with drastically elevated interleukin 6 (IL-6) level in critically ill COVID-19 patients. medRxiv.

61. Brocke-Heidrich K, Kretzschmar AK, Pfeifer G, Henze C, Loffler D, Koczan D, Thiesen HJ, Burger R, Gramatzki M, Horn F. 2004. Interleukin-6-dependent gene expression profiles in multiple myeloma INA-6 cells reveal a Bcl-2 family-independent survival pathway closely associated with Stat3 activation. Blood 103:242–51.

62. Srirangan S, Choy EH. 2010. The role of interleukin 6 in the pathophysiology of rheumatoid arthritis. Ther Adv Musculoskelet Dis 2:247–56.

63. Somers EC, Eschenauer GA, Troost JP, Golob JL, Gandhi TN, Wang L, Zhou N, Petty LA, Baang JH, Dillman NO, Frame D, Gregg KS, Kaul DR, Nagel J, Patel TS, Zhou S, Lauring AS, Hanauer DA, Martin E, Sharma P, Fung CM, Pogue JM. 2020. Tocilizumab for treatment of mechanically ventilated patients with COVID-19. medRxiv doi:10.1101/2020.05.29.20117358.

64. Shu Q, Lennemann NJ, Sarkar SN, Sadovsky Y, Coyne CB. 2015. ADAP2 Is an Interferon Stimulated Gene That Restricts RNA Virus Entry. PLoS Pathog 11:e1005150.

65. de Silva S, Wu L. 2011. TRIM5 acts as more than a retroviral restriction factor. Viruses 3:1204–9.

66. Lian Q, Sun B. 2017. Interferons command Trim22 to fight against viruses. Cell Mol Immunol 14:794–796.

67. Barr SD, Smiley JR, Bushman FD. 2008. The interferon response inhibits HIV particle production by induction of TRIM22. PLoS Pathog 4:e1000007.

68. Lee HC, Lee ES, Uddin MB, Kim TH, Kim JH, Chathuranga K, Chathuranga WAG, Jin M, Kim S, Kim CJ, Lee JS. 2019. Released Tryptophanyl-tRNA Synthetase Stimulates Innate Immune Responses against Viral Infection. J Virol 93.

69. Landry ML, Foxman EF. 2018. Antiviral Response in the Nasopharynx Identifies Patients With Respiratory Virus Infection. J Infect Dis 217:897–905.

70. Rabanal-Ruiz Y, Korolchuk VI. 2018. mTORC1 and Nutrient Homeostasis: The Central Role of the Lysosome. Int J Mol Sci 19.

71. Kapuy O, Vinod PK, Banhegyi G. 2014. mTOR inhibition increases cell viability via autophagy induction during endoplasmic reticulum stress-An experimental and modeling study. FEBS Open Bio 4:704–13.

72. Gassen NC, Papies J, Bajaj T, Dethloff F, Emanuel J, Weckmann K, Heinz DE, Heinemann N, Lennarz M, Richter A, Niemeyer D, Corman VM, Giavalisco P, Drosten C, Müller MA. 2020. Analysis of SARS-CoV-2-controlled autophagy reveals spermidine, MK-2206, and niclosamide as putative antiviral therapeutics. bioRxiv doi:10.1101/2020.04.15.997254:2020.04.15.997254.

73. Prentice E, Jerome WG, Yoshimori T, Mizushima N, Denison MR. 2004. Coronavirus replication complex formation utilizes components of cellular autophagy. J Biol Chem 279:10136–41.

74. Mohamud Y, Shi J, Qu J, Poon T, Xue YC, Deng H, Zhang J, Luo H. 2018. Enteroviral Infection Inhibits Autophagic Flux via Disruption of the SNARE Complex to Enhance Viral Replication. Cell Rep 22:3292–3303.

75. Metz P, Chiramel A, Chatel-Chaix L, Alvisi G, Bankhead P, Mora-Rodriguez R, Long G, Hamacher-Brady A, Brady NR, Bartenschlager R. 2015. Dengue Virus Inhibition of Autophagic Flux and Dependency of Viral Replication on Proteasomal Degradation of the Autophagy Receptor p62. J Virol 89:8026–41.

76. Yang N, Shen HM. 2020. Targeting the Endocytic Pathway and Autophagy Process as a Novel Therapeutic Strategy in COVID-19. Int J Biol Sci 16:1724–1731.

77. Gassen NC, Niemeyer D, Muth D, Corman VM, Martinelli S, Gassen A, Hafner K, Papies J, Mosbauer K, Zellner A, Zannas AS, Herrmann A, Holsboer F, Brack-Werner R, Boshart M, Muller-Myhsok B, Drosten C, Muller MA, Rein T. 2019. SKP2 attenuates autophagy through Beclin1-ubiquitination and its inhibition reduces MERS-Coronavirus infection. Nat Commun 10:5770.

78. Weikel KA, Cacicedo JM, Ruderman NB, Ido Y. 2016. Knockdown of GSK3beta increases basal autophagy and AMPK signalling in nutrient-laden human aortic endothelial cells. Biosci Rep 36.

79. Ren F, Zhang L, Zhang X, Shi H, Wen T, Bai L, Zheng S, Chen Y, Chen D, Li L, Duan Z. 2016. Inhibition of glycogen synthase kinase 3beta promotes autophagy to protect mice from acute liver failure mediated by peroxisome proliferator-activated receptor alpha. Cell Death Dis 7:e2151.

80. Marchand B, Arsenault D, Raymond-Fleury A, Boisvert FM, Boucher MJ. 2015. Glycogen synthase kinase-3 (GSK3) inhibition induces prosurvival autophagic signals in human pancreatic cancer cells. J Biol Chem 290:5592–605.

81. Yim WW, Mizushima N. 2020. Lysosome biology in autophagy. Cell Discov 6:6.

82. Anand SK, Tikoo SK. 2013. Viruses as modulators of mitochondrial functions. Adv Virol 2013:738794.

83. Sanchez-Caballero L, Guerrero-Castillo S, Nijtmans L. 2016. Unraveling the complexity of mitochondrial complex I assembly: A dynamic process. Biochim Biophys Acta 1857:980–90.

84. Mokranjac D, Neupert W. 2010. The many faces of the mitochondrial TIM23 complex. Biochim Biophys Acta 1797:1045–54.

85. Rambold AS, Lippincott-Schwartz J. 2011. Mechanisms of mitochondria and autophagy crosstalk. Cell Cycle 10:4032–8.

86. Graef M, Nunnari J. 2011. Mitochondria regulate autophagy by conserved signalling pathways. EMBO J 30:2101–14.

87. Morita M, Prudent J, Basu K, Goyon V, Katsumura S, Hulea L, Pearl D, Siddiqui N, Strack S, McGuirk S, St-Pierre J, Larsson O, Topisirovic I, Vali H, McBride HM, Bergeron JJ, Sonenberg N. 2017. mTOR Controls Mitochondrial Dynamics and Cell Survival via MTFP1. Mol Cell 67:922–935 e5.

88. Tondera D, Czauderna F, Paulick K, Schwarzer R, Kaufmann J, Santel A. 2005. The mitochondrial protein MTP18 contributes to mitochondrial fission in mammalian cells. J Cell Sci 118:3049–59.

89. Das R, Chakrabarti O. 2020. Mitochondrial hyperfusion: a friend or a foe. Biochem Soc Trans 48:631–644.

90. Barbier V, Lang D, Valois S, Rothman AL, Medin CL. 2017. Dengue virus induces mitochondrial elongation through impairment of Drp1-triggered mitochondrial fission. Virology 500:149–160.

91. Shi CS, Qi HY, Boularan C, Huang NN, Abu-Asab M, Shelhamer JH, Kehrl JH. 2014. SARS-coronavirus open reading frame-9b suppresses innate immunity by targeting mitochondria and the MAVS/TRAF3/TRAF6 signalosome. J Immunol 193:3080–9.

92. Baracca A, Chiaradonna F, Sgarbi G, Solaini G, Alberghina L, Lenaz G. 2010. Mitochondrial Complex I decrease is responsible for bioenergetic dysfunction in K-ras transformed cells. Biochim Biophys Acta 1797:314–23.

93. Simonnet H, Demont J, Pfeiffer K, Guenaneche L, Bouvier R, Brandt U, Schagger H, Godinot C. 2003. Mitochondrial complex I is deficient in renal oncocytomas. Carcinogenesis 24:1461–6.

94. Angajala A, Lim S, Phillips JB, Kim JH, Yates C, You Z, Tan M. 2018. Diverse Roles of Mitochondria in Immune Responses: Novel Insights Into Immuno-Metabolism. Front Immunol 9:1605.

95. Bonora E, Porcelli AM, Gasparre G, Biondi A, Ghelli A, Carelli V, Baracca A, Tallini G, Martinuzzi A, Lenaz G, Rugolo M, Romeo G. 2006. Defective oxidative phosphorylation in thyroid oncocytic carcinoma is associated with pathogenic mitochondrial DNA mutations affecting complexes I and III. Cancer Res 66:6087–96.

96. Holownia A, Wielgat P, Rysiak E, Braszko JJ. 2016. Intracellular and Extracellular Cytokines in A549 Cells and THP1 Cells Exposed to Cigarette Smoke. Adv Exp Med Biol 910:39–45.

97. Lieber M, Smith B, Szakal A, Nelson-Rees W, Todaro G. 1976. A continuous tumor-cell line from a human lung carcinoma with properties of type II alveolar epithelial cells. Int J Cancer 17:62–70.

98. Martens K, Hellings PW, Steelant B. 2018. Calu-3 epithelial cells exhibit different immune and epithelial barrier responses from freshly isolated primary nasal epithelial cells in vitro. Clin Transl Allergy 8:40.

99. Shen BQ, Finkbeiner WE, Wine JJ, Mrsny RJ, Widdicombe JH. 1994. Calu-3: a human airway epithelial cell line that shows cAMP-dependent Cl-secretion. Am J Physiol 266:L493–501.

100. Zhang X, Chu H, Wen L, Shuai H, Yang D, Wang Y, Hou Y, Zhu Z, Yuan S, Yin F, Chan JF, Yuen KY. 2020. Competing endogenous RNA network profiling reveals novel host dependency factors required for MERS-CoV propagation. Emerg Microbes Infect 9:733–746.

101. Travaglini KJ, Nabhan AN, Penland L, Sinha R, Gillich A, Sit RV, Chang S, Conley SD, Mori Y, Seita J, Berry GJ, Shrager JB, Metzger RJ, Kuo CS, Neff N, Weissman IL, Quake SR, Krasnow MA. 2020. A molecular cell atlas of the human lung from single cell RNA sequencing. bioRxiv doi:10.1101/742320:742320.

102. Ritchie ME, Phipson B, Wu D, Hu Y, Law CW, Shi W, Smyth GK. 2015. limma powers differential expression analyses for RNA-sequencing and microarray studies. Nucleic Acids Res 43:e47.

103. Law CW, Chen Y, Shi W, Smyth GK. 2014. voom: Precision weights unlock linear model analysis tools for RNA-seq read counts. Genome Biol 15:R29.

104. Yu G, Wang LG, Han Y, He QY. 2012. clusterProfiler: an R package for comparing biological themes among gene clusters. OMICS 16:284–7.

105. Leek JT, Johnson WE, Parker HS, Jaffe AE, Storey JD. 2012. The sva package for removing batch effects and other unwanted variation in high-throughput experiments. Bioinformatics 28:882–3.

106. Mostafavi S, Ray D, Warde-Farley D, Grouios C, Morris Q. 2008. GeneMANIA: a real-time multiple association network integration algorithm for predicting gene function. Genome Biol 9 Suppl 1:S4.

107. Shannon P, Markiel A, Ozier O, Baliga NS, Wang JT, Ramage D, Amin N, Schwikowski B, Ideker T. 2003. Cytoscape: a software environment for integrated models of biomolecular interaction networks. Genome Res 13:2498–504.

108. Merico D, Isserlin R, Stueker O, Emili A, Bader GD. 2010. Enrichment map: a network-based method for gene-set enrichment visualization and interpretation. PLoS One 5:e13984.

109. Kucera M, Isserlin R, Arkhangorodsky A, Bader GD. 2016. AutoAnnotate: A Cytoscape app for summarizing networks with semantic annotations. F1000Res 5:1717.

110. Mehdi P. 2020. Supplemental material for Singh et al. 2020 mBio doi:10.6084/m9.figshare.12272351.v1.

111. Stuart T, Butler A, Hoffman P, Hafemeister C, Papalexi E, Mauck WM, 3rd, Hao Y, Stoeckius M, Smibert P, Satija R. 2019. Comprehensive Integration of Single-Cell Data. Cell 177:1888-1902 e21.

