## Supplemental Figures 1-8 and Tables 1-2 for "Network Analysis and Transcriptome Profiling Identify Autophagic and Mitochondrial Dysfunctions in SARS-CoV-2 Infection"

Supp Figure S1

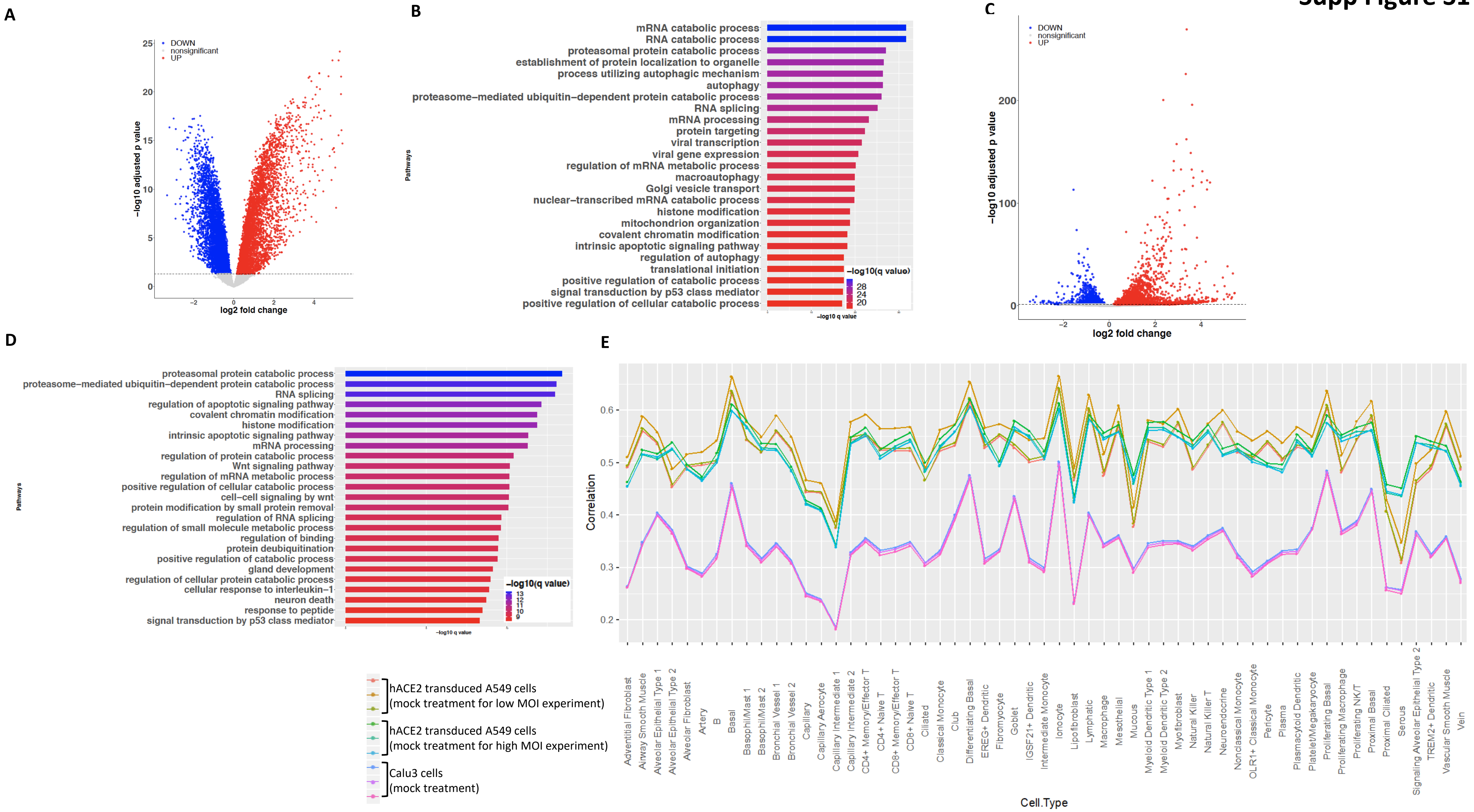

#### Supp Figure S2

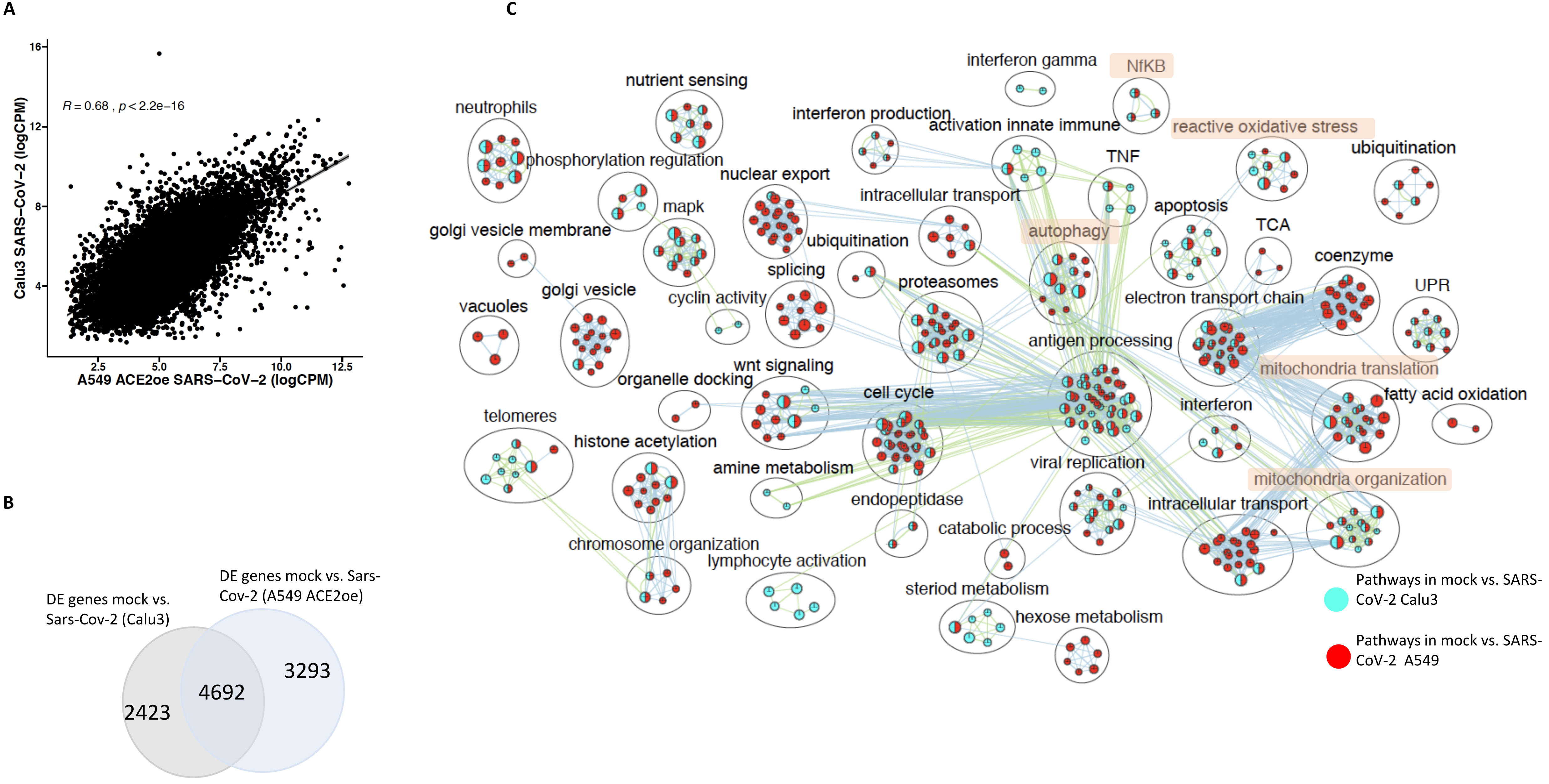

Supp Figure S3

A

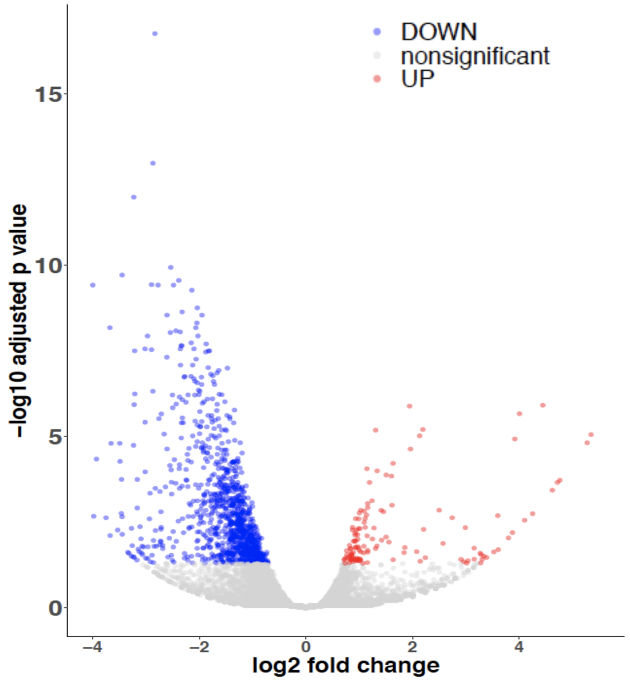


B

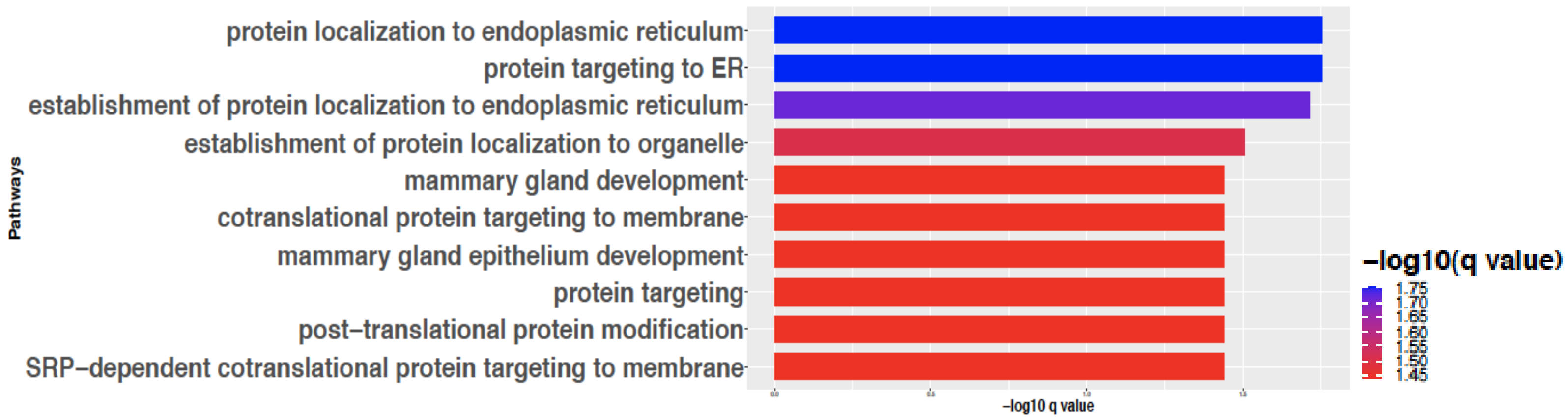


C

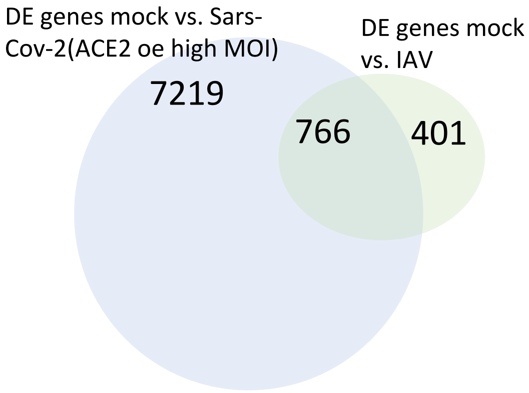

Supp Figure S4

A

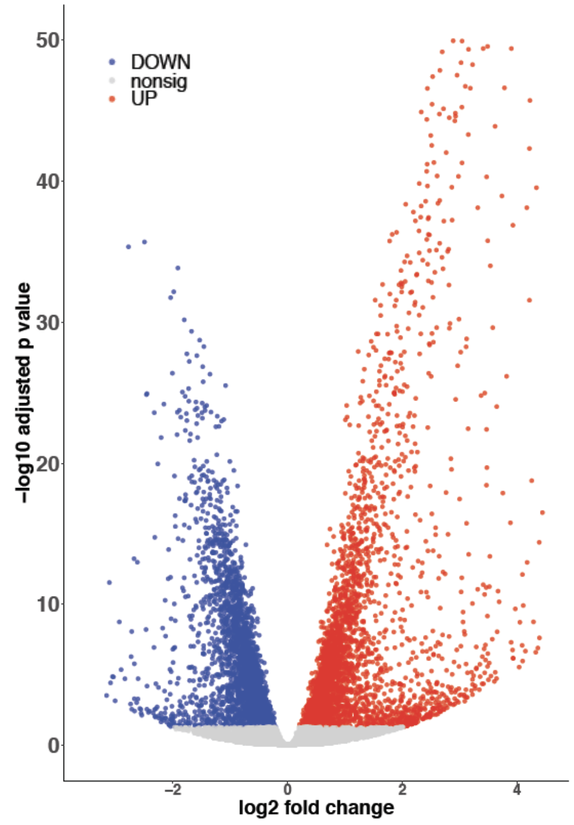


B

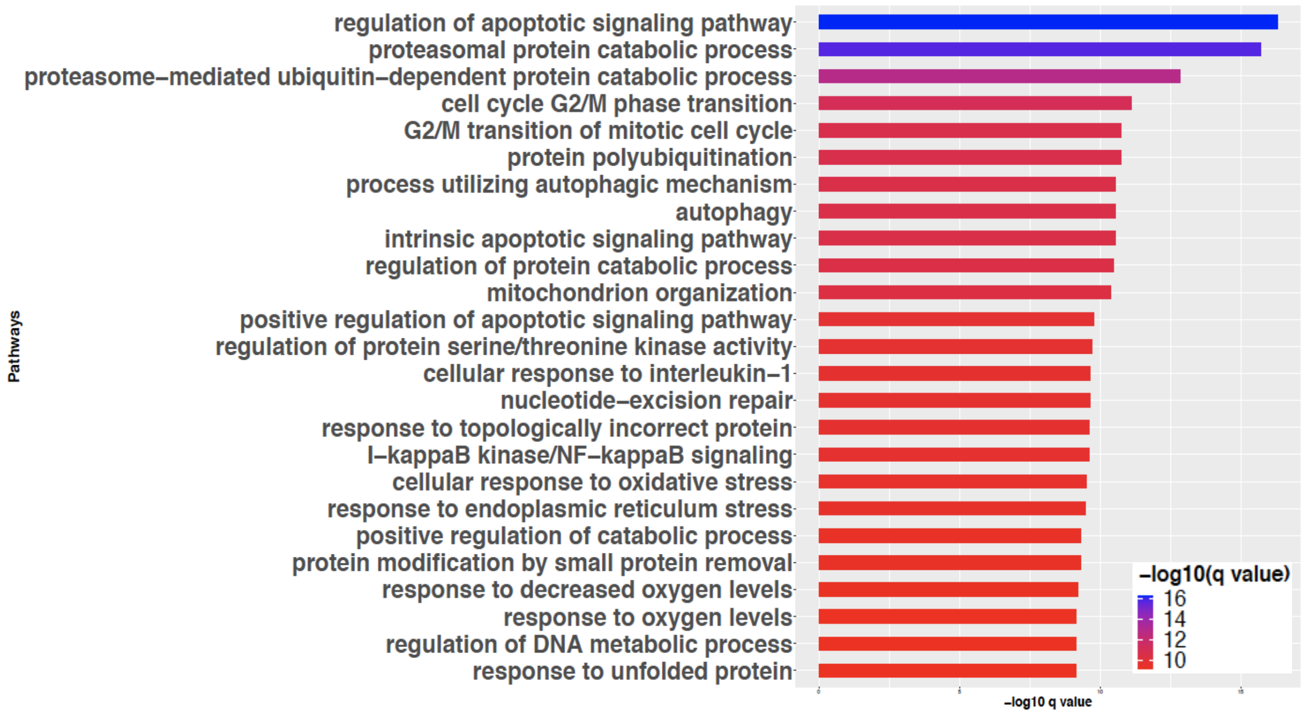

### Supp Figure S5

A

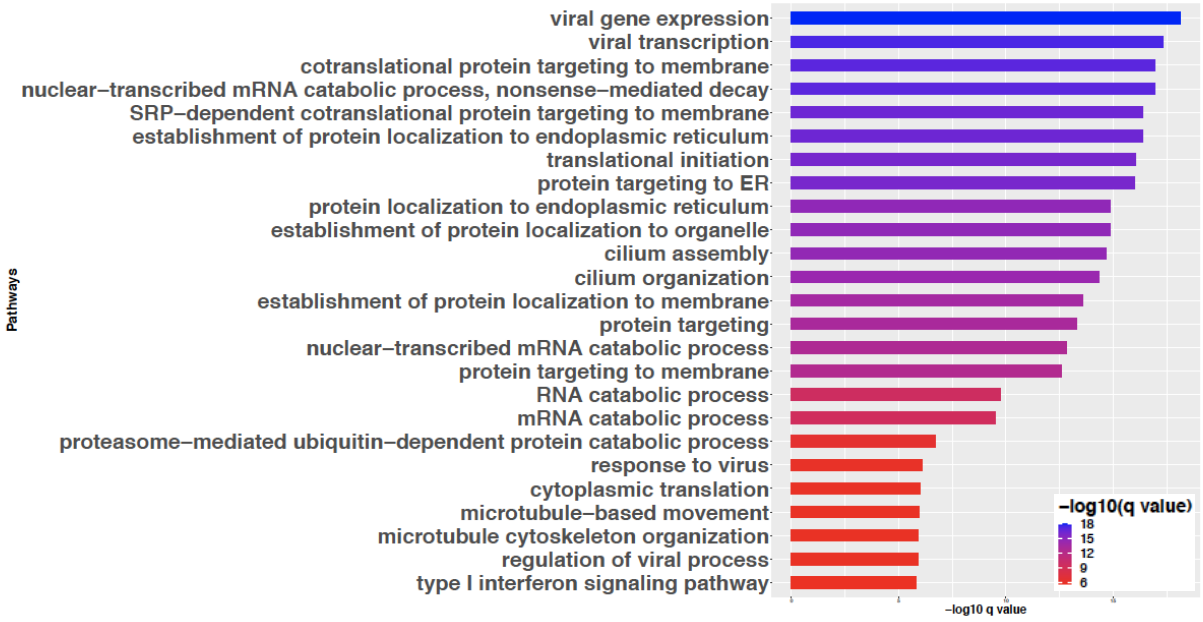


B

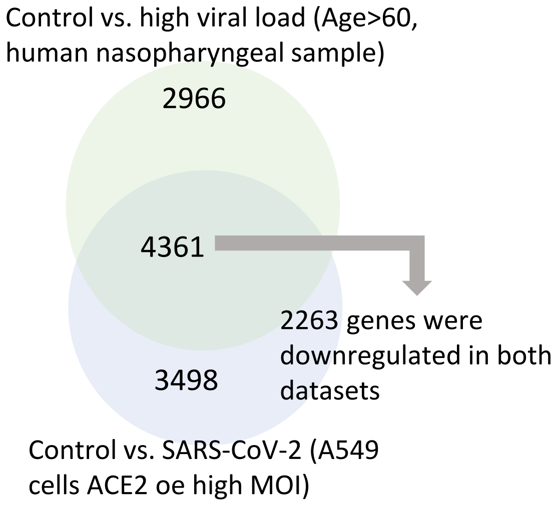


C

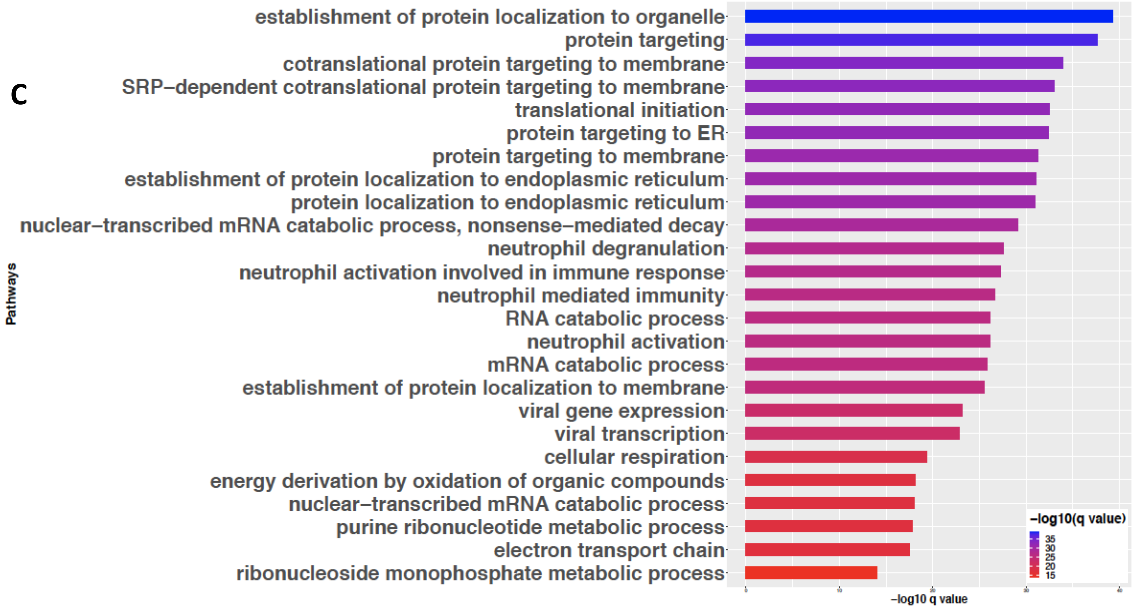

### Supp Figure S6

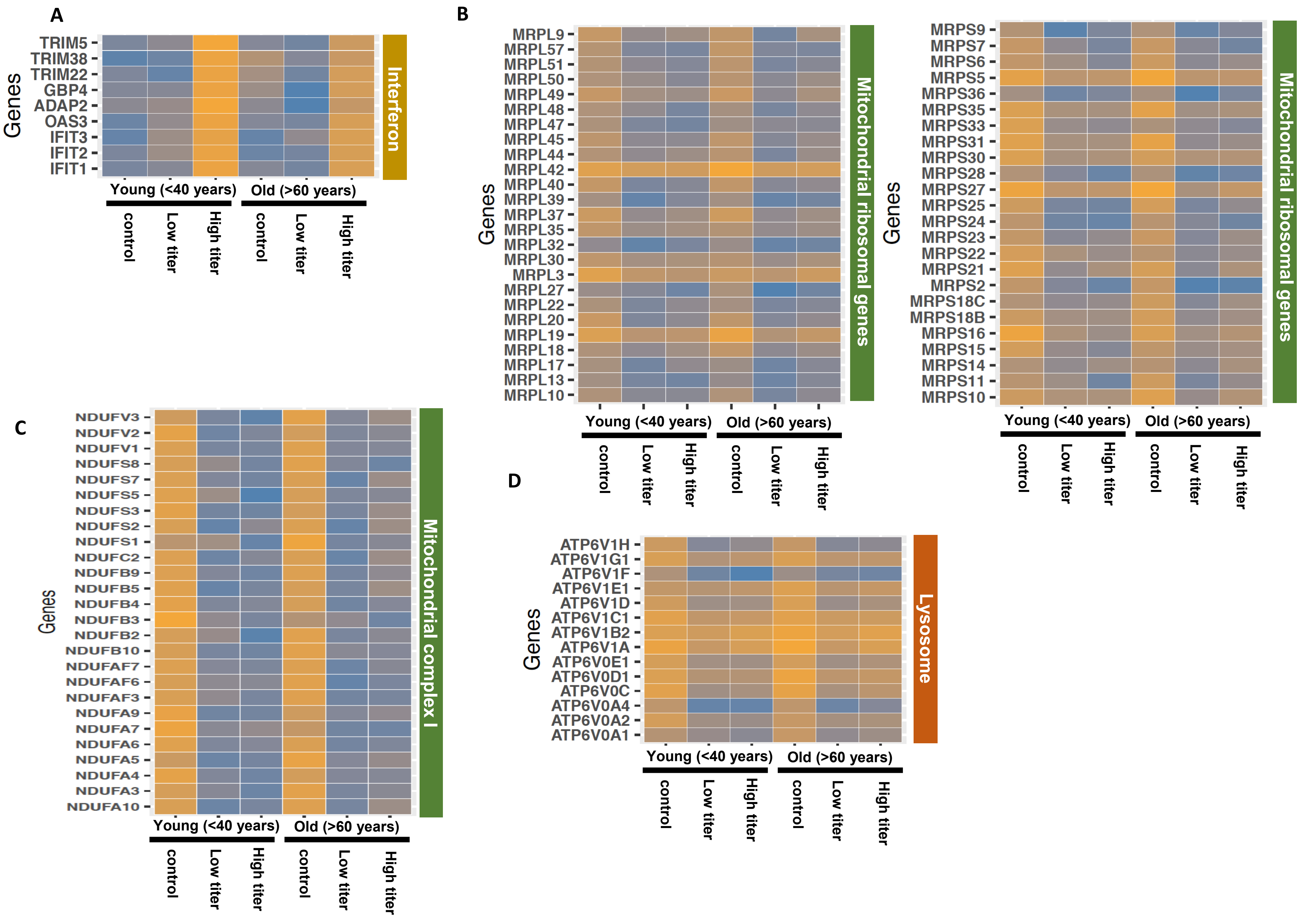

**Supp Figure S7**

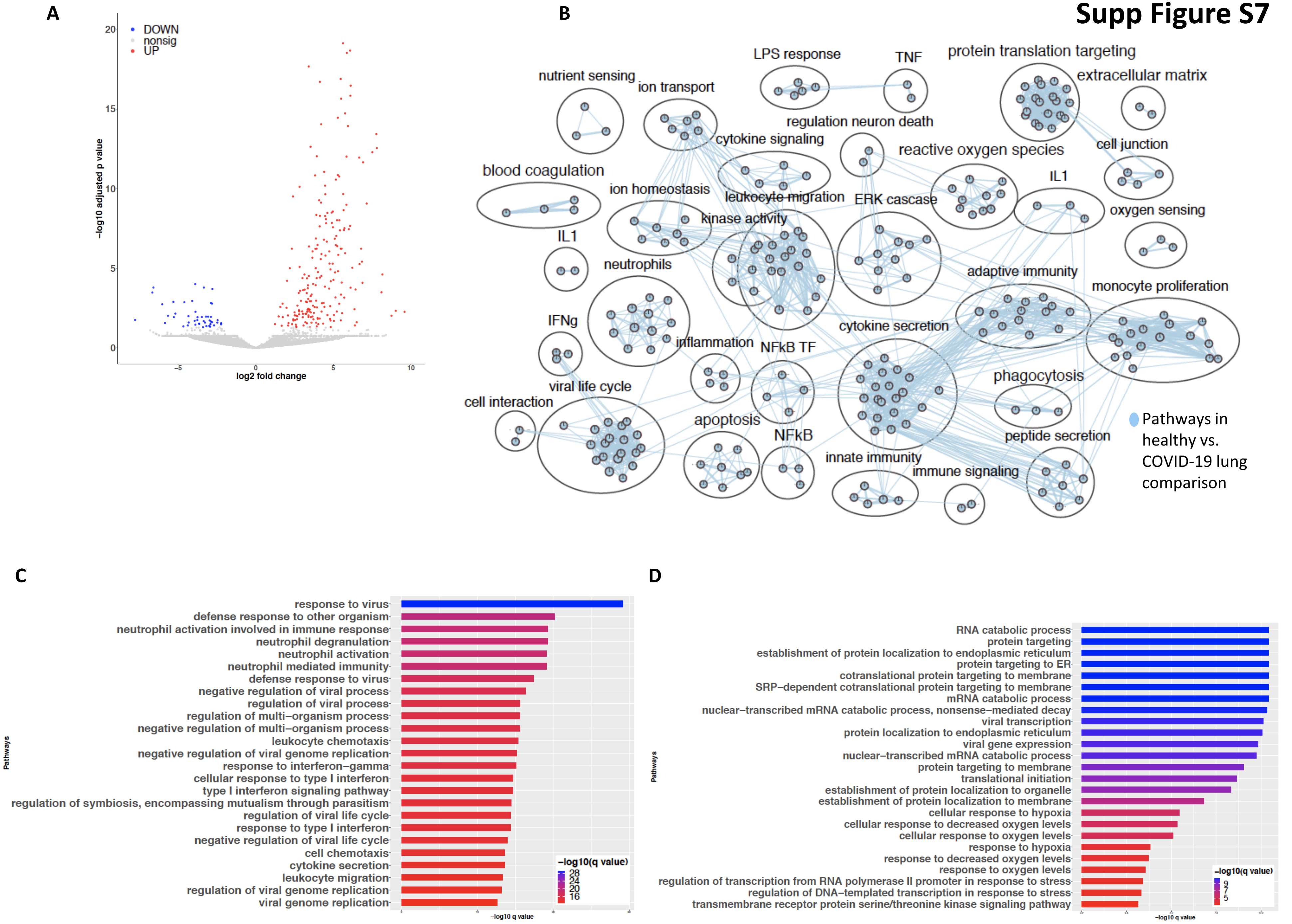

| Table S1: Consensus module name and | | | | | |
|---|---|---|---|---|---|
| Module Name | Module Size | Significant Genes Overlap | Module Name | Module Size | Significant Genes Overlap |
| bisque4 | 51 | 27 | magenta | 158 | 77 |
| black | 176 | 118 | mediumpurp | 56 | 28 |
| blue | 2093 | 1806 | midnightblue | 123 | 78 |
| brown | 431 | 308 | orange | 83 | 42 |
| brown4 | 52 | 36 | orangered4 | 58 | 36 |
| cyan | 125 | 92 | paleturquois | 77 | 60 |
| darkgreen | 89 | 55 | pink | 162 | 128 |
| darkgrey | 84 | 55 | plum1 | 62 | 43 |
| darkmagenta | 64 | 54 | purple | 144 | 74 |
| darkolivegre | 66 | 46 | red | 183 | 116 |
| darkorange | 81 | 44 | royalblue | 106 | 54 |
| darkorange2 | 53 | 27 | saddlebrown | 78 | 36 |
| darkred | 94 | 67 | salmon | 127 | 68 |
| darkturquois | 86 | 43 | sienna3 | 63 | 32 |
| floralwhite | 53 | 36 | skyblue | 78 | 48 |
| green | 186 | 87 | skyblue3 | 62 | 42 |
| greenyellow | 139 | 101 | steelblue | 78 | 31 |
| grey60 | 113 | 70 | tan | 139 | 89 |
| ivory | 53 | 39 | turquoise | 2096 | 1804 |
| lightcyan | 114 | 79 | violet | 69 | 36 |
| lightcyan1 | 53 | 29 | white | 79 | 47 |
| lightgreen | 108 | 59 | yellow | 314 | 171 |
| lightsteelblu | 55 | 38 | yellowgreen | 62 | 43 |
| lightyellow | 107 | 66 | | | |

| Table S2: Marker genes in each cluster | | | | | | |
|---|---|---|---|---|---|---|
| gene | cluster | p_val | avg_logFC | pct.1 | pct.2 | p_val_adj |
| LUM | Adventitial Fibroblast | 0 | 2.786590029 | 0.996 | 0.046 | 0 |
| C1R | Adventitial Fibroblast | 0 | 2.230799138 | 0.957 | 0.104 | 0 |
| C1S | Adventitial Fibroblast | 0 | 2.12550214 | 0.964 | 0.067 | 0 |
| COL6A2 | Adventitial Fibroblast | 0 | 1.949707642 | 0.969 | 0.095 | 0 |
| C3 | Adventitial Fibroblast | 0 | 1.625204903 | 0.806 | 0.095 | 0 |
| EFEMP1 | Adventitial Fibroblast | 0 | 1.579792258 | 0.763 | 0.108 | 0 |
| MYC | Adventitial Fibroblast | 0 | 1.533787556 | 0.826 | 0.13 | 0 |
| RARRES1 | Adventitial Fibroblast | 0 | 1.387014784 | 0.601 | 0.064 | 0 |
| LTBP4 | Adventitial Fibroblast | 0 | 1.34734777 | 0.903 | 0.094 | 0 |
| SERPINA3 | Adventitial Fibroblast | 0 | 1.321580056 | 0.637 | 0.027 | 0 |
| MEG3 | Adventitial Fibroblast | 0 | 1.266829057 | 0.596 | 0.026 | 0 |
| FSTL1 | Adventitial Fibroblast | 0 | 1.239723918 | 0.749 | 0.075 | 0 |
| PLTP | Adventitial Fibroblast | 0 | 1.16386539 | 0.801 | 0.079 | 0 |
| UAP1 | Adventitial Fibroblast | 0 | 1.077900945 | 0.614 | 0.12 | 0 |
| C11orf96 | Adventitial Fibroblast | 1.50E-291 | 1.114677374 | 0.598 | 0.112 | 3.06E-287 |
| FHL1 | Adventitial Fibroblast | 1.30E-281 | 1.117944403 | 0.862 | 0.29 | 2.65E-277 |
| CCL2 | Adventitial Fibroblast | 2.11E-279 | 2.330734285 | 0.668 | 0.156 | 4.30E-275 |
| IGFBP7 | Adventitial Fibroblast | 7.64E-207 | 1.17851979 | 0.878 | 0.334 | 1.56E-202 |
| JUNB | Adventitial Fibroblast | 6.55E-196 | 1.554021352 | 0.955 | 0.75 | 1.33E-191 |
| SOCS3 | Adventitial Fibroblast | 3.69E-170 | 1.268265343 | 0.811 | 0.375 | 7.51E-166 |
| DUSP1 | Adventitial Fibroblast | 7.44E-166 | 1.093371235 | 0.982 | 0.866 | 1.51E-161 |
| MT1A | Adventitial Fibroblast | 8.50E-155 | 1.263691019 | 0.506 | 0.133 | 1.73E-150 |
| FOSB | Adventitial Fibroblast | 1.22E-152 | 1.320675502 | 0.695 | 0.28 | 2.49E-148 |
| PNRC1 | Adventitial Fibroblast | 1.45E-122 | 1.003578227 | 0.883 | 0.641 | 2.95E-118 |
| IL6 | Adventitial Fibroblast | 6.38E-110 | 1.625276577 | 0.452 | 0.142 | 1.30E-105 |
| GADD45B | Adventitial Fibroblast | 5.58E-95 | 1.17446996 | 0.697 | 0.365 | 1.14E-90 |
| ACTA2 | Airway Smooth Muscle | 0 | 3.46383406 | 0.999 | 0.082 | 0 |
| MYL9 | Airway Smooth Muscle | 0 | 3.017381895 | 0.999 | 0.179 | 0 |
| IGFBP7 | Airway Smooth Muscle | 0 | 2.321618285 | 0.999 | 0.331 | 0 |
| TPM1 | Airway Smooth Muscle | 0 | 1.741045717 | 0.981 | 0.273 | 0 |
| MFGE8 | Airway Smooth Muscle | 0 | 1.382008804 | 0.876 | 0.108 | 0 |
| C11orf96 | Airway Smooth Muscle | 0 | 1.35900598 | 0.724 | 0.11 | 0 |
| MAP1B | Airway Smooth Muscle | 0 | 1.045935725 | 0.727 | 0.04 | 0 |
| MT1A | Airway Smooth Muscle | 2.42E-110 | 1.028382373 | 0.418 | 0.133 | 4.92E-106 |
| CD55 | Alveolar Epithelial Type | 0 | 1.765846081 | 0.964 | 0.374 | 0 |
| ANXA3 | Alveolar Epithelial Type | 0 | 1.544484009 | 0.919 | 0.158 | 0 |
| CD9 | Alveolar Epithelial Type | 0 | 1.426255262 | 0.996 | 0.661 | 0 |
| SLC39A8 | Alveolar Epithelial Type | 0 | 1.409703442 | 0.923 | 0.115 | 0 |
| MYL9 | Alveolar Epithelial Type | 0 | 1.39117199 | 0.956 | 0.176 | 0 |
| RAB11FIP1 | Alveolar Epithelial Type | 0 | 1.375543319 | 0.899 | 0.189 | 0 |
| CLDN4 | Alveolar Epithelial Type | 0 | 1.292853547 | 0.817 | 0.107 | 0 |
| KLF6 | Alveolar Epithelial Type | 0 | 1.241313753 | 0.961 | 0.629 | 0 |
| ICAM1 | Alveolar Epithelial Type | 0 | 1.20581191 | 0.909 | 0.275 | 0 |

| | | | | | | |
|---|---|---|---|---|---|---|
| CAV2 | Alveolar Epithelial Type | 0 | 1.191728797 | 0.952 | 0.275 | 0 |
| APLP2 | Alveolar Epithelial Type | 0 | 1.136831369 | 0.948 | 0.492 | 0 |
| SLPI | Alveolar Epithelial Type | 0 | 1.878738268 | 0.996 | 0.176 | 0 |
| CXCL2 | Alveolar Epithelial Type | 0 | 1.723323158 | 0.839 | 0.225 | 0 |
| LAMP3 | Alveolar Epithelial Type | 0 | 1.329693634 | 0.92 | 0.02 | 0 |
| C11orf96 | Alveolar Epithelial Type | 0 | 1.222678159 | 0.793 | 0.074 | 0 |
| ABCA3 | Alveolar Epithelial Type | 0 | 1.16127674 | 0.879 | 0.025 | 0 |
| AREG | Alveolar Epithelial Type | 0 | 1.103382842 | 0.721 | 0.184 | 0 |
| SDC4 | Alveolar Epithelial Type | 0 | 1.070143991 | 0.871 | 0.178 | 0 |
| SDR16C5 | Alveolar Epithelial Type | 0 | 1.036025167 | 0.813 | 0.038 | 0 |
| LUM | Alveolar Fibroblast | 0 | 3.078860561 | 0.987 | 0.035 | 0 |
| COL6A2 | Alveolar Fibroblast | 0 | 1.538556201 | 0.91 | 0.086 | 0 |
| FHL1 | Alveolar Fibroblast | 0 | 1.523997458 | 0.94 | 0.281 | 0 |
| C1S | Alveolar Fibroblast | 0 | 1.500879306 | 0.856 | 0.059 | 0 |
| C1R | Alveolar Fibroblast | 0 | 1.288662061 | 0.819 | 0.097 | 0 |
| MACF1 | Alveolar Fibroblast | 0 | 1.241721333 | 0.803 | 0.162 | 0 |
| G0S2 | Alveolar Fibroblast | 0 | 1.21200183 | 0.602 | 0.07 | 0 |
| LTBP4 | Alveolar Fibroblast | 0 | 1.146861184 | 0.82 | 0.086 | 0 |
| LIMCH1 | Alveolar Fibroblast | 0 | 1.099595346 | 0.785 | 0.096 | 0 |
| DKK3 | Alveolar Fibroblast | 0 | 1.075098057 | 0.765 | 0.049 | 0 |
| SERPINA3 | Alveolar Fibroblast | 0 | 1.036220536 | 0.43 | 0.024 | 0 |
| CCL2 | Alveolar Fibroblast | 4.21E-165 | 1.352282783 | 0.424 | 0.155 | 8.57E-161 |
| MT1A | Artery | 0 | 1.911629805 | 0.654 | 0.124 | 0 |
| CXCL2 | Artery | 0 | 1.640122089 | 0.695 | 0.25 | 0 |
| EMP1 | Artery | 0 | 1.385755577 | 0.794 | 0.232 | 0 |
| CTNNAL1 | Artery | 0 | 1.359097047 | 0.861 | 0.135 | 0 |
| EPAS1 | Artery | 0 | 1.295336923 | 0.99 | 0.344 | 0 |
| MT2A | Artery | 0 | 1.250994712 | 0.993 | 0.758 | 0 |
| TFPI | Artery | 0 | 1.225280101 | 0.901 | 0.244 | 0 |
| TSPAN7 | Artery | 0 | 1.083194188 | 0.796 | 0.118 | 0 |
| IL33 | Artery | 0 | 1.040389236 | 0.767 | 0.078 | 0 |
| SRPX | Artery | 0 | 1.003528294 | 0.737 | 0.061 | 0 |
| IL6 | Artery | 3.11E-256 | 1.272206529 | 0.439 | 0.137 | 6.33E-252 |
| LTB | B | 0 | 1.703832645 | 0.913 | 0.098 | 0 |
| RPS27 | B | 0 | 1.377221664 | 1 | 0.995 | 0 |
| RPL39 | B | 0 | 1.138428334 | 1 | 0.979 | 0 |
| RPL13A | B | 0 | 1.12874843 | 1 | 0.995 | 0 |
| RPS23 | B | 0 | 1.081575606 | 1 | 0.986 | 0 |
| RPS8 | B | 0 | 1.065513987 | 1 | 0.984 | 0 |
| RPL23A | B | 0 | 1.038917475 | 1 | 0.983 | 0 |
| RPSA | B | 0 | 1.025482632 | 1 | 0.906 | 0 |
| RPL37 | B | 0 | 1.024511068 | 1 | 0.973 | 0 |
| RPS18 | B | 0 | 1.015328931 | 1 | 0.993 | 0 |
| RCSD1 | B | 0 | 1.01495427 | 0.799 | 0.133 | 0 |
| RPL34 | B | 0 | 1.001440921 | 1 | 0.992 | 0 |

| | | | | | | |
|---|---|---|---|---|---|---|
| KRT15 | Basal | 0 | 1.884304498 | 0.784 | 0.02 | 0 |
| ERRFI1 | Basal | 0 | 1.167132207 | 0.78 | 0.139 | 0 |
| LAMB3 | Basal | 0 | 1.110547199 | 0.837 | 0.048 | 0 |
| MYC | Basal | 2.98E-290 | 1.05906622 | 0.697 | 0.132 | 6.07E-286 |
| IER3 | Basal | 8.88E-254 | 1.260804702 | 0.729 | 0.165 | 1.81E-249 |
| CD9 | Basal | 8.60E-170 | 1.053711092 | 0.97 | 0.664 | 1.75E-165 |
| CXCL1 | Basal | 3.14E-169 | 1.954447191 | 0.537 | 0.121 | 6.39E-165 |
| TSC22D3 | Basophil/Mast 1 | 0 | 1.038745298 | 0.973 | 0.656 | 0 |
| AREG | Basophil/Mast 1 | 2.18E-228 | 1.245803697 | 0.553 | 0.208 | 4.43E-224 |
| BIRC3 | Basophil/Mast 2 | 0 | 1.799216945 | 0.924 | 0.218 | 0 |
| SRGN | Basophil/Mast 2 | 0 | 1.795134864 | 1 | 0.792 | 0 |
| AREG | Basophil/Mast 2 | 0 | 1.755138551 | 0.815 | 0.21 | 0 |
| NFKBIA | Basophil/Mast 2 | 0 | 1.701406796 | 1 | 0.771 | 0 |
| CREM | Basophil/Mast 2 | 0 | 1.505479187 | 0.88 | 0.189 | 0 |
| CPM | Basophil/Mast 2 | 0 | 1.253602618 | 0.904 | 0.212 | 0 |
| CSF1 | Basophil/Mast 2 | 0 | 1.174825666 | 0.799 | 0.131 | 0 |
| TNFAIP3 | Basophil/Mast 2 | 0 | 1.140770708 | 0.931 | 0.25 | 0 |
| TNFRSF9 | Basophil/Mast 2 | 0 | 1.085641938 | 0.73 | 0.01 | 0 |
| PTGS2 | Basophil/Mast 2 | 0 | 1.02255169 | 0.609 | 0.057 | 0 |
| FOSB | Basophil/Mast 2 | 1.08E-306 | 1.07249491 | 0.897 | 0.278 | 2.19E-302 |
| GADD45B | Basophil/Mast 2 | 2.20E-251 | 1.215040961 | 0.929 | 0.363 | 4.49E-247 |
| SDCBP | Basophil/Mast 2 | 1.05E-237 | 1.061626093 | 0.978 | 0.639 | 2.13E-233 |
| DDIT4 | Basophil/Mast 2 | 3.41E-237 | 1.026575298 | 0.895 | 0.328 | 6.94E-233 |
| PLIN2 | Basophil/Mast 2 | 2.34E-198 | 1.108371384 | 0.857 | 0.348 | 4.77E-194 |
| SELE | Bronchial Vessel 1 | 0 | 1.84341532 | 0.343 | 0.01 | 0 |
| MYC | Bronchial Vessel 1 | 0 | 1.25981007 | 0.719 | 0.132 | 0 |
| IGFBP7 | Bronchial Vessel 1 | 3.15E-290 | 1.726165669 | 0.965 | 0.334 | 6.41E-286 |
| EMP1 | Bronchial Vessel 1 | 1.72E-205 | 1.165592937 | 0.802 | 0.241 | 3.51E-201 |
| SOCS3 | Bronchial Vessel 1 | 3.26E-188 | 1.384362615 | 0.879 | 0.375 | 6.64E-184 |
| IL6 | Bronchial Vessel 1 | 1.12E-91 | 1.775003887 | 0.455 | 0.142 | 2.28E-87 |
| CCL2 | Bronchial Vessel 1 | 6.37E-47 | 1.071184335 | 0.398 | 0.159 | 1.30E-42 |
| EMP1 | Bronchial Vessel 2 | 1.57E-183 | 1.706785826 | 0.928 | 0.242 | 3.21E-179 |
| SOCS3 | Bronchial Vessel 2 | 9.15E-132 | 1.636335778 | 0.945 | 0.376 | 1.86E-127 |
| MT2A | Bronchial Vessel 2 | 4.07E-75 | 1.465991051 | 0.991 | 0.763 | 8.28E-71 |
| SOCS2 | Bronchial Vessel 2 | 1.43E-71 | 1.010244019 | 0.57 | 0.167 | 2.92E-67 |
| CCL2 | Bronchial Vessel 2 | 2.07E-41 | 1.262389577 | 0.468 | 0.16 | 4.21E-37 |
| EPAS1 | Capillary | 0 | 1.569223972 | 0.987 | 0.279 | 0 |
| AKAP12 | Capillary | 0 | 1.235216151 | 0.575 | 0.097 | 0 |
| MT2A | Capillary | 0 | 1.192902856 | 0.996 | 0.734 | 0 |
| CD59 | Capillary | 0 | 1.006696798 | 0.977 | 0.619 | 0 |
| HLA-E | Capillary | 0 | 1.001252428 | 1 | 0.873 | 0 |
| HLA-E | Capillary Aerocyte | 0 | 1.630291389 | 1 | 0.88 | 0 |
| APP | Capillary Aerocyte | 0 | 1.307051117 | 0.919 | 0.252 | 0 |
| SERPINE1 | Capillary Aerocyte | 0 | 1.206177658 | 0.57 | 0.078 | 0 |
| PDLIM1 | Capillary Aerocyte | 0 | 1.201891905 | 0.976 | 0.596 | 0 |

| | | | | | | |
|---|---|---|---|---|---|---|
| ESAM | Capillary Aerocyte | 0 | 1.15382022 | 0.918 | 0.203 | 0 |
| RHOB | Capillary Aerocyte | 0 | 1.148835301 | 0.93 | 0.468 | 0 |
| IFNGR1 | Capillary Aerocyte | 0 | 1.104567853 | 0.906 | 0.404 | 0 |
| CAV2 | Capillary Aerocyte | 0 | 1.052730529 | 0.899 | 0.242 | 0 |
| JUN | Capillary Aerocyte | 0 | 1.00881753 | 0.979 | 0.593 | 0 |
| EPAS1 | Capillary Intermediate 1 | 0 | 1.456091416 | 1 | 0.353 | 0 |
| SERPINE1 | Capillary Intermediate 1 | 0 | 1.371532474 | 0.798 | 0.104 | 0 |
| CX3CL1 | Capillary Intermediate 1 | 0 | 1.318659672 | 0.887 | 0.135 | 0 |
| HLA-E | Capillary Intermediate 1 | 0 | 1.274783513 | 1 | 0.886 | 0 |
| AKAP12 | Capillary Intermediate 1 | 0 | 1.256393851 | 0.838 | 0.144 | 0 |
| ESAM | Capillary Intermediate 1 | 0 | 1.224291981 | 0.992 | 0.243 | 0 |
| PDLIM1 | Capillary Intermediate 1 | 0 | 1.156240857 | 0.995 | 0.618 | 0 |
| APP | Capillary Intermediate 1 | 0 | 1.056021484 | 0.978 | 0.29 | 0 |
| ARHGAP29 | Capillary Intermediate 1 | 0 | 1.008913591 | 0.944 | 0.216 | 0 |
| RHOB | Capillary Intermediate 1 | 6.02E-285 | 1.085276589 | 0.992 | 0.494 | 1.22E-280 |
| CSF3 | Capillary Intermediate 1 | 5.33E-284 | 1.082588688 | 0.525 | 0.096 | 1.09E-279 |
| EMP1 | Capillary Intermediate 1 | 1.19E-271 | 1.032287386 | 0.812 | 0.239 | 2.43E-267 |
| ICAM1 | Capillary Intermediate 1 | 1.56E-226 | 1.136090767 | 0.803 | 0.279 | 3.17E-222 |
| MT2A | Capillary Intermediate 1 | 3.61E-196 | 1.0293594 | 1 | 0.761 | 7.35E-192 |
| EPAS1 | Capillary Intermediate 2 | 2.92E-239 | 1.260948699 | 0.963 | 0.354 | 5.95E-235 |
| IL32 | CD4+ Memory/Effector T | 0 | 1.367355189 | 0.958 | 0.353 | 0 |
| LTB | CD4+ Memory/Effector T | 0 | 1.208570503 | 0.657 | 0.08 | 0 |
| BTG1 | CD4+ Memory/Effector T | 0 | 1.12216113 | 0.992 | 0.707 | 0 |
| TSC22D3 | CD4+ Memory/Effector T | 0 | 1.106162868 | 0.967 | 0.648 | 0 |
| CCL5 | CD4+ Memory/Effector T | 0 | 1.011757492 | 0.731 | 0.145 | 0 |
| LTB | CD4+ Naive T | 0 | 1.661789742 | 0.958 | 0.095 | 0 |
| RPS27 | CD4+ Naive T | 0 | 1.517082546 | 1 | 0.995 | 0 |
| RPS12 | CD4+ Naive T | 0 | 1.295506121 | 1 | 0.985 | 0 |
| RPS15A | CD4+ Naive T | 0 | 1.256920807 | 1 | 0.987 | 0 |
| RPL34 | CD4+ Naive T | 0 | 1.250350415 | 1 | 0.992 | 0 |
| RPL39 | CD4+ Naive T | 0 | 1.241588509 | 1 | 0.979 | 0 |
| RPS6 | CD4+ Naive T | 0 | 1.234368294 | 1 | 0.987 | 0 |
| RPL32 | CD4+ Naive T | 0 | 1.204709159 | 1 | 0.992 | 0 |
| RPS3A | CD4+ Naive T | 0 | 1.181221239 | 1 | 0.985 | 0 |
| RPL30 | CD4+ Naive T | 0 | 1.175999212 | 1 | 0.983 | 0 |
| RPL31 | CD4+ Naive T | 0 | 1.16513278 | 1 | 0.965 | 0 |
| RPS18 | CD4+ Naive T | 0 | 1.148329926 | 1 | 0.993 | 0 |
| RPL13A | CD4+ Naive T | 0 | 1.139367342 | 1 | 0.995 | 0 |
| RPL4 | CD4+ Naive T | 0 | 1.127375171 | 1 | 0.893 | 0 |
| RPL36A | CD4+ Naive T | 0 | 1.121459321 | 0.915 | 0.417 | 0 |
| RPL37 | CD4+ Naive T | 0 | 1.111473185 | 1 | 0.973 | 0 |
| GAS5 | CD4+ Naive T | 0 | 1.104400027 | 0.95 | 0.42 | 0 |
| RPL3 | CD4+ Naive T | 0 | 1.103919365 | 1 | 0.992 | 0 |
| EEF1G | CD4+ Naive T | 0 | 1.100430572 | 1 | 0.942 | 0 |
| RPL23A | CD4+ Naive T | 0 | 1.089333354 | 1 | 0.983 | 0 |

| | | | | | | |
|---|---|---|---|---|---|---|
| RPL35A | CD4+ Naive T | 0 | 1.086961826 | 1 | 0.985 | 0 |
| RPS27A | CD4+ Naive T | 0 | 1.082878265 | 1 | 0.995 | 0 |
| RPSA | CD4+ Naive T | 0 | 1.079851771 | 1 | 0.906 | 0 |
| RPL38 | CD4+ Naive T | 0 | 1.064240747 | 1 | 0.945 | 0 |
| RPS8 | CD4+ Naive T | 0 | 1.035175388 | 1 | 0.984 | 0 |
| RPL11 | CD4+ Naive T | 0 | 1.023419135 | 1 | 0.993 | 0 |
| RPL10A | CD4+ Naive T | 0 | 1.017963561 | 1 | 0.961 | 0 |
| RPL5 | CD4+ Naive T | 0 | 1.015173456 | 1 | 0.936 | 0 |
| RPS4X | CD4+ Naive T | 0 | 1.008160883 | 1 | 0.978 | 0 |
| RPS23 | CD4+ Naive T | 0 | 1.000084552 | 1 | 0.986 | 0 |
| CCL5 | CD8+ Memory/Effector T | 0 | 1.65924011 | 0.966 | 0.158 | 0 |
| RPS27 | CD8+ Memory/Effector T | 0 | 1.156428402 | 1 | 0.995 | 0 |
| IL32 | CD8+ Memory/Effector T | 0 | 1.027832655 | 0.961 | 0.37 | 0 |
| CCL5 | CD8+ Naive T | 0 | 2.014546049 | 0.96 | 0.147 | 0 |
| IL32 | CD8+ Naive T | 0 | 1.098806478 | 0.947 | 0.362 | 0 |
| SAA1 | Ciliated | 0 | 2.252977849 | 0.446 | 0.057 | 0 |
| SAA2 | Ciliated | 0 | 2.002001417 | 0.397 | 0.031 | 0 |
| ODF3B | Ciliated | 0 | 1.994019763 | 0.977 | 0.163 | 0 |
| CCDC146 | Ciliated | 0 | 1.64055031 | 0.952 | 0.014 | 0 |
| LCN2 | Ciliated | 0 | 1.638736568 | 0.632 | 0.071 | 0 |
| CCDC170 | Ciliated | 0 | 1.617381413 | 0.951 | 0.021 | 0 |
| DNAH5 | Ciliated | 0 | 1.483915487 | 0.93 | 0.013 | 0 |
| EFHC1 | Ciliated | 0 | 1.479297433 | 0.96 | 0.077 | 0 |
| LRRC23 | Ciliated | 0 | 1.435576718 | 0.932 | 0.029 | 0 |
| RSPH9 | Ciliated | 0 | 1.388353707 | 0.902 | 0.007 | 0 |
| CLDN4 | Ciliated | 0 | 1.326561191 | 0.924 | 0.101 | 0 |
| SRI | Ciliated | 0 | 1.313381775 | 0.971 | 0.406 | 0 |
| CRNDE | Ciliated | 0 | 1.237282143 | 0.912 | 0.08 | 0 |
| NUCB2 | Ciliated | 0 | 1.216717569 | 0.893 | 0.216 | 0 |
| DMKN | Ciliated | 0 | 1.188792664 | 0.905 | 0.076 | 0 |
| CDS1 | Ciliated | 0 | 1.150885811 | 0.886 | 0.032 | 0 |
| AKAP9 | Ciliated | 0 | 1.149054831 | 0.937 | 0.267 | 0 |
| MLF1 | Ciliated | 0 | 1.114118609 | 0.902 | 0.115 | 0 |
| STK33 | Ciliated | 0 | 1.100667595 | 0.852 | 0.009 | 0 |
| DYNC2H1 | Ciliated | 0 | 1.078888935 | 0.871 | 0.029 | 0 |
| PPIL6 | Ciliated | 0 | 1.068052926 | 0.842 | 0.012 | 0 |
| CCDC113 | Ciliated | 0 | 1.041213556 | 0.827 | 0.005 | 0 |
| ALCAM | Ciliated | 0 | 1.026016414 | 0.89 | 0.139 | 0 |
| S100A8 | Classical Monocyte | 0 | 3.15071556 | 0.981 | 0.261 | 0 |
| S100A9 | Classical Monocyte | 0 | 3.14335825 | 0.994 | 0.426 | 0 |
| CTSS | Classical Monocyte | 0 | 1.377304718 | 0.998 | 0.519 | 0 |
| CSTA | Classical Monocyte | 0 | 1.231830039 | 0.943 | 0.263 | 0 |
| LST1 | Classical Monocyte | 0 | 1.108474501 | 0.975 | 0.294 | 0 |
| AP1S2 | Classical Monocyte | 0 | 1.031417561 | 0.825 | 0.218 | 0 |
| RPL39 | Classical Monocyte | 0 | 1.000918679 | 1 | 0.978 | 0 |

| | | | | | | |
|---|---|---|---|---|---|---|
| SLPI | Club | 0 | 2.671090349 | 0.987 | 0.213 | 0 |
| CLDN4 | Club | 0 | 1.056889215 | 0.808 | 0.108 | 0 |
| ATP1B1 | Club | 0 | 1.045449346 | 0.814 | 0.221 | 0 |
| SOX4 | Club | 0 | 1.012977507 | 0.665 | 0.136 | 0 |
| AREG | Club | 1.45E-144 | 1.081117553 | 0.541 | 0.211 | 2.95E-140 |
| SERPINB3 | Differentiating Basal | 0 | 2.835092522 | 0.706 | 0.012 | 0 |
| CLDN4 | Differentiating Basal | 0 | 1.667690272 | 0.94 | 0.114 | 0 |
| KRT15 | Differentiating Basal | 0 | 1.146520815 | 0.615 | 0.022 | 0 |
| SERPINB4 | Differentiating Basal | 0 | 1.048640523 | 0.426 | 0.006 | 0 |
| MDK | Differentiating Basal | 1.69E-247 | 1.261617891 | 0.804 | 0.14 | 3.45E-243 |
| SLPI | Differentiating Basal | 2.04E-200 | 1.991293603 | 0.909 | 0.221 | 4.15E-196 |
| RPLP0 | Differentiating Basal | 3.71E-143 | 1.170810819 | 1 | 0.916 | 7.56E-139 |
| CXCL1 | Differentiating Basal | 2.44E-124 | 1.16682453 | 0.589 | 0.122 | 4.97E-120 |
| EREG | EREG+ Dendritic | 7.08E-235 | 1.016147962 | 0.599 | 0.043 | 1.44E-230 |
| G0S2 | EREG+ Dendritic | 6.84E-87 | 1.705843756 | 0.514 | 0.08 | 1.39E-82 |
| IER3 | EREG+ Dendritic | 1.20E-81 | 1.133486894 | 0.711 | 0.168 | 2.45E-77 |
| AREG | EREG+ Dendritic | 2.17E-79 | 1.523660327 | 0.803 | 0.214 | 4.41E-75 |
| NAMPT | EREG+ Dendritic | 1.16E-78 | 1.109432692 | 0.965 | 0.375 | 2.36E-74 |
| SRGN | EREG+ Dendritic | 7.32E-67 | 1.190812698 | 1 | 0.793 | 1.49E-62 |
| IL1B | EREG+ Dendritic | 9.64E-64 | 1.194330871 | 0.676 | 0.169 | 1.96E-59 |
| PLAUR | EREG+ Dendritic | 3.24E-63 | 1.112441964 | 0.859 | 0.297 | 6.61E-59 |
| C15orf48 | EREG+ Dendritic | 1.34E-60 | 1.186129022 | 0.577 | 0.13 | 2.72E-56 |
| HLA-DQA1 | EREG+ Dendritic | 1.49E-47 | 1.003512303 | 0.944 | 0.453 | 3.04E-43 |
| LTBP1 | Fibromyocyte | 0 | 1.195961055 | 0.827 | 0.035 | 0 |
| GEM | Fibromyocyte | 6.67E-280 | 1.453617765 | 0.816 | 0.049 | 1.36E-275 |
| DKK3 | Fibromyocyte | 3.68E-258 | 1.543171677 | 0.878 | 0.062 | 7.48E-254 |
| ACTA2 | Fibromyocyte | 5.21E-241 | 3.067765367 | 0.99 | 0.09 | 1.06E-236 |
| COL6A2 | Fibromyocyte | 2.34E-160 | 1.224129253 | 0.908 | 0.101 | 4.77E-156 |
| MYL9 | Fibromyocyte | 9.99E-124 | 2.09425671 | 0.99 | 0.187 | 2.03E-119 |
| FILIP1L | Fibromyocyte | 4.14E-98 | 1.401994947 | 0.857 | 0.159 | 8.43E-94 |
| FHL1 | Fibromyocyte | 3.02E-85 | 1.656235034 | 0.98 | 0.293 | 6.16E-81 |
| IGFBP7 | Fibromyocyte | 1.23E-67 | 1.643721605 | 0.99 | 0.337 | 2.50E-63 |
| TPM1 | Fibromyocyte | 4.02E-59 | 1.153090286 | 0.929 | 0.28 | 8.19E-55 |
| GADD45B | Fibromyocyte | 5.46E-11 | 1.156686377 | 0.633 | 0.367 | 1.11E-06 |
| LCN2 | Goblet | 0 | 3.342277345 | 0.994 | 0.08 | 0 |
| SERPINB3 | Goblet | 0 | 3.064383002 | 0.912 | 0.013 | 0 |
| CFB | Goblet | 0 | 1.365219525 | 0.862 | 0.075 | 0 |
| CLDN10 | Goblet | 0 | 1.331012713 | 0.843 | 0.013 | 0 |
| ASS1 | Goblet | 0 | 1.320424954 | 0.893 | 0.077 | 0 |
| SERPINB4 | Goblet | 0 | 1.194991617 | 0.535 | 0.007 | 0 |
| CDC42EP5 | Goblet | 0 | 1.07067372 | 0.855 | 0.041 | 0 |
| CREB3L1 | Goblet | 0 | 1.011198103 | 0.774 | 0.041 | 0 |
| BIK | Goblet | 0 | 1.008297567 | 0.83 | 0.028 | 0 |
| SPINT1 | Goblet | 2.24E-262 | 1.030034356 | 0.931 | 0.114 | 4.55E-258 |
| MDK | Goblet | 1.16E-260 | 2.032325437 | 0.981 | 0.141 | 2.36E-256 |

| | | | | | | |
|---|---|---|---|---|---|---|
| CLDN4 | Goblet | 9.88E-252 | 1.532100344 | 0.937 | 0.116 | 2.01E-247 |
| SLPI | Goblet | 1.87E-185 | 3.184910521 | 1 | 0.222 | 3.81E-181 |
| C3 | Goblet | 4.54E-170 | 1.032222497 | 0.748 | 0.099 | 9.23E-166 |
| CXCL1 | Goblet | 2.50E-157 | 1.491889238 | 0.799 | 0.122 | 5.08E-153 |
| HEBP2 | Goblet | 1.34E-152 | 1.105661522 | 0.899 | 0.186 | 2.72E-148 |
| ATP1B1 | Goblet | 1.13E-140 | 1.260680085 | 0.95 | 0.228 | 2.31E-136 |
| C15orf48 | Goblet | 2.98E-126 | 1.775894637 | 0.736 | 0.13 | 6.07E-122 |
| NUCB2 | Goblet | 1.11E-119 | 1.411097533 | 0.874 | 0.228 | 2.26E-115 |
| XBP1 | Goblet | 2.53E-111 | 1.6625422 | 0.981 | 0.423 | 5.14E-107 |
| CD9 | Goblet | 2.20E-66 | 1.07506251 | 0.981 | 0.665 | 4.47E-62 |
| HLA-DQA1 | IGSF21+ Dendritic | 1.60E-156 | 1.561736414 | 0.979 | 0.452 | 3.26E-152 |
| LST1 | Intermediate Monocyte | 3.34E-163 | 1.75360999 | 1 | 0.314 | 6.80E-159 |
| NAMPT | Intermediate Monocyte | 8.36E-100 | 1.110519647 | 0.941 | 0.374 | 1.70E-95 |
| SAT1 | Intermediate Monocyte | 5.97E-98 | 1.293106599 | 1 | 0.847 | 1.22E-93 |
| CTSS | Intermediate Monocyte | 1.03E-88 | 1.109284975 | 1 | 0.533 | 2.10E-84 |
| SCG2 | Ionocyte | 0 | 1.547379095 | 0.682 | 0.002 | 0 |
| FAM24B | Ionocyte | 4.18E-104 | 1.166699655 | 0.636 | 0.019 | 8.50E-100 |
| BIK | Ionocyte | 1.04E-83 | 1.100839019 | 0.727 | 0.03 | 2.12E-79 |
| PFN2 | Ionocyte | 1.54E-57 | 1.058895427 | 0.864 | 0.062 | 3.13E-53 |
| TPD52 | Ionocyte | 8.30E-45 | 1.508468881 | 0.955 | 0.108 | 1.69E-40 |
| CLDN4 | Ionocyte | 1.05E-37 | 1.333097135 | 0.955 | 0.118 | 2.14E-33 |
| SOX4 | Ionocyte | 1.70E-28 | 1.175933859 | 0.955 | 0.143 | 3.45E-24 |
| ATP1B1 | Ionocyte | 1.40E-20 | 1.416787075 | 0.955 | 0.23 | 2.85E-16 |
| SMS | Ionocyte | 3.64E-16 | 1.022358214 | 0.864 | 0.24 | 7.41E-12 |
| CD9 | Ionocyte | 2.47E-15 | 1.917322806 | 1 | 0.666 | 5.02E-11 |
| APLP2 | Ionocyte | 1.70E-12 | 1.091222178 | 0.955 | 0.499 | 3.46E-08 |
| FST | Lipofibroblast | 0 | 1.953409514 | 0.857 | 0.009 | 0 |
| ALDH1A3 | Lipofibroblast | 0 | 1.447865029 | 0.943 | 0.021 | 0 |
| MEDAG | Lipofibroblast | 0 | 1.388733329 | 0.829 | 0.008 | 0 |
| GFPT2 | Lipofibroblast | 0 | 1.109336779 | 0.857 | 0.012 | 0 |
| MLLT11 | Lipofibroblast | 1.04E-303 | 1.296283525 | 0.771 | 0.015 | 2.12E-299 |
| TFPI2 | Lipofibroblast | 1.23E-132 | 1.736232812 | 0.943 | 0.051 | 2.50E-128 |
| C1S | Lipofibroblast | 1.22E-102 | 2.259195612 | 0.971 | 0.074 | 2.49E-98 |
| RARRES1 | Lipofibroblast | 3.83E-99 | 1.75237774 | 0.943 | 0.068 | 7.79E-95 |
| GEM | Lipofibroblast | 7.46E-89 | 1.220392749 | 0.771 | 0.049 | 1.52E-84 |
| C3 | Lipofibroblast | 1.75E-84 | 2.484340643 | 1 | 0.1 | 3.55E-80 |
| SRPX | Lipofibroblast | 3.17E-84 | 1.568588364 | 0.914 | 0.076 | 6.46E-80 |
| COL6A2 | Lipofibroblast | 9.79E-76 | 2.104969852 | 0.971 | 0.102 | 1.99E-71 |
| CYP1B1 | Lipofibroblast | 1.06E-71 | 1.639201113 | 0.771 | 0.063 | 2.15E-67 |
| C1R | Lipofibroblast | 6.54E-71 | 2.074512958 | 0.971 | 0.11 | 1.33E-66 |
| EFEMP1 | Lipofibroblast | 1.59E-65 | 1.35614908 | 0.971 | 0.113 | 3.24E-61 |
| MYC | Lipofibroblast | 1.02E-55 | 2.200312507 | 0.943 | 0.136 | 2.08E-51 |
| LUM | Lipofibroblast | 2.02E-54 | 1.185253536 | 0.657 | 0.054 | 4.11E-50 |
| ELL2 | Lipofibroblast | 1.05E-47 | 1.032427079 | 0.829 | 0.115 | 2.13E-43 |
| CREM | Lipofibroblast | 6.15E-43 | 1.861637551 | 0.971 | 0.195 | 1.25E-38 |

| | | | | | | |
|---|---|---|---|---|---|---|
| MT1A | Lipofibroblast | 1.06E-40 | 1.618671558 | 0.886 | 0.136 | 2.15E-36 |
| C11orf96 | Lipofibroblast | 2.42E-39 | 1.888095111 | 0.771 | 0.116 | 4.93E-35 |
| UAP1 | Lipofibroblast | 8.36E-37 | 1.034420083 | 0.771 | 0.124 | 1.70E-32 |
| PHLDA1 | Lipofibroblast | 1.84E-36 | 2.030816244 | 0.971 | 0.239 | 3.75E-32 |
| CCL2 | Lipofibroblast | 7.29E-29 | 1.10487582 | 0.829 | 0.16 | 1.48E-24 |
| NAMPT | Lipofibroblast | 4.14E-27 | 1.452543835 | 1 | 0.376 | 8.43E-23 |
| MT2A | Lipofibroblast | 2.29E-24 | 2.325726527 | 1 | 0.763 | 4.65E-20 |
| B4GALT1 | Lipofibroblast | 1.14E-23 | 1.155291092 | 0.886 | 0.238 | 2.33E-19 |
| CDKN1A | Lipofibroblast | 4.60E-23 | 1.06836998 | 0.8 | 0.205 | 9.35E-19 |
| JUNB | Lipofibroblast | 1.61E-21 | 1.734778428 | 1 | 0.752 | 3.28E-17 |
| MCL1 | Lipofibroblast | 2.78E-21 | 1.480060197 | 1 | 0.655 | 5.66E-17 |
| ZFP36L1 | Lipofibroblast | 3.05E-21 | 1.463558667 | 0.943 | 0.391 | 6.21E-17 |
| PNRC1 | Lipofibroblast | 1.47E-20 | 1.467478275 | 1 | 0.643 | 2.99E-16 |
| SOCS3 | Lipofibroblast | 7.26E-20 | 1.23767289 | 0.943 | 0.378 | 1.48E-15 |
| CEBPB | Lipofibroblast | 7.75E-18 | 1.389968049 | 1 | 0.704 | 1.58E-13 |
| DUSP1 | Lipofibroblast | 1.14E-15 | 1.224461088 | 1 | 0.867 | 2.32E-11 |
| CXCL2 | Lipofibroblast | 1.29E-14 | 1.494818885 | 0.771 | 0.26 | 2.63E-10 |
| IGFBP7 | Lymphatic | 0 | 2.380256735 | 1 | 0.333 | 0 |
| TFPI | Lymphatic | 0 | 1.487724254 | 0.961 | 0.253 | 0 |
| AKAP12 | Lymphatic | 0 | 1.462761825 | 0.85 | 0.146 | 0 |
| EFEMP1 | Lymphatic | 0 | 1.268983583 | 0.694 | 0.109 | 0 |
| PPFIBP1 | Lymphatic | 0 | 1.159310744 | 0.872 | 0.144 | 0 |
| THBD | Lymphatic | 6.79E-242 | 1.126838808 | 0.835 | 0.251 | 1.38E-237 |
| CD9 | Lymphatic | 2.99E-203 | 1.234843127 | 0.983 | 0.664 | 6.08E-199 |
| CD59 | Lymphatic | 7.44E-195 | 1.028783841 | 0.991 | 0.657 | 1.51E-190 |
| MARCO | Macrophage | 0 | 2.201537743 | 0.97 | 0.078 | 0 |
| CCL20 | Macrophage | 0 | 2.029166059 | 0.385 | 0.117 | 0 |
| FTH1 | Macrophage | 0 | 1.963256098 | 1 | 0.999 | 0 |
| CD68 | Macrophage | 0 | 1.809531415 | 0.979 | 0.142 | 0 |
| IL1B | Macrophage | 0 | 1.715364915 | 0.52 | 0.068 | 0 |
| S100A11 | Macrophage | 0 | 1.691016679 | 1 | 0.892 | 0 |
| LGALS3 | Macrophage | 0 | 1.677982675 | 0.997 | 0.608 | 0 |
| ALOX5AP | Macrophage | 0 | 1.520971685 | 0.983 | 0.356 | 0 |
| CTSC | Macrophage | 0 | 1.394627856 | 0.953 | 0.274 | 0 |
| OLR1 | Macrophage | 0 | 1.359594282 | 0.939 | 0.038 | 0 |
| TREM1 | Macrophage | 0 | 1.357226106 | 0.91 | 0.062 | 0 |
| HLA-DQA1 | Macrophage | 0 | 1.299102429 | 0.969 | 0.306 | 0 |
| CXCL5 | Macrophage | 0 | 1.287616358 | 0.411 | 0.016 | 0 |
| SNX10 | Macrophage | 0 | 1.269985376 | 0.909 | 0.083 | 0 |
| CXCL3 | Macrophage | 0 | 1.158748812 | 0.633 | 0.113 | 0 |
| PSAP | Macrophage | 0 | 1.147320751 | 0.998 | 0.732 | 0 |
| CTSB | Macrophage | 0 | 1.14205641 | 0.982 | 0.348 | 0 |
| CTSS | Macrophage | 0 | 1.141720974 | 0.993 | 0.402 | 0 |
| LTA4H | Macrophage | 0 | 1.132272517 | 0.895 | 0.184 | 0 |
| VIM | Macrophage | 0 | 1.066901338 | 1 | 0.837 | 0 |

| | | | | | | |
|---|---|---|---|---|---|---|
| CSTA | Macrophage | 0 | 1.019709732 | 0.909 | 0.103 | 0 |
| PRG4 | Mesothelial | 0 | 1.947154995 | 0.862 | 0.006 | 0 |
| PAPPA | Mesothelial | 0 | 1.211065752 | 0.724 | 0.005 | 0 |
| ALDH1A3 | Mesothelial | 6.10E-237 | 1.979881986 | 0.897 | 0.021 | 1.24E-232 |
| STEAP1 | Mesothelial | 4.50E-230 | 1.043967231 | 0.759 | 0.015 | 9.15E-226 |
| CLDN1 | Mesothelial | 3.29E-119 | 1.246674656 | 0.828 | 0.036 | 6.70E-115 |
| TFPI2 | Mesothelial | 1.35E-91 | 1.793702169 | 0.862 | 0.051 | 2.75E-87 |
| C1S | Mesothelial | 3.41E-83 | 2.019559727 | 0.966 | 0.074 | 6.94E-79 |
| MEST | Mesothelial | 2.50E-81 | 1.115955184 | 0.724 | 0.04 | 5.09E-77 |
| RARRES1 | Mesothelial | 3.13E-81 | 1.932162483 | 0.931 | 0.068 | 6.37E-77 |
| CFB | Mesothelial | 3.14E-77 | 2.554189363 | 0.931 | 0.076 | 6.39E-73 |
| CCDC71L | Mesothelial | 3.26E-65 | 1.373014066 | 0.793 | 0.062 | 6.63E-61 |
| C3 | Mesothelial | 2.63E-60 | 2.805293274 | 0.931 | 0.1 | 5.36E-56 |
| C1R | Mesothelial | 3.12E-57 | 1.819000063 | 0.966 | 0.11 | 6.36E-53 |
| COL6A2 | Mesothelial | 1.74E-54 | 1.652506981 | 0.931 | 0.102 | 3.54E-50 |
| EFEMP1 | Mesothelial | 4.55E-49 | 1.27150229 | 0.931 | 0.113 | 9.27E-45 |
| CCL2 | Mesothelial | 2.99E-37 | 1.876436883 | 0.966 | 0.16 | 6.09E-33 |
| ERRFI1 | Mesothelial | 1.05E-35 | 1.091746088 | 0.897 | 0.143 | 2.13E-31 |
| PIM1 | Mesothelial | 1.79E-26 | 1.092363919 | 0.793 | 0.158 | 3.63E-22 |
| MDK | Mesothelial | 1.18E-25 | 1.102407455 | 0.759 | 0.143 | 2.41E-21 |
| RPS4Y1 | Mesothelial | 1.07E-20 | 1.174701108 | 0.966 | 0.368 | 2.18E-16 |
| RPS12 | Mesothelial | 9.03E-17 | 1.038334979 | 1 | 0.985 | 1.84E-12 |
| MT2A | Mesothelial | 8.76E-16 | 1.508219114 | 1 | 0.763 | 1.78E-11 |
| RPL12 | Mesothelial | 2.47E-15 | 1.014091818 | 1 | 0.964 | 5.03E-11 |
| GAS5 | Mesothelial | 4.39E-14 | 1.033018436 | 0.931 | 0.427 | 8.93E-10 |
| SAA1 | Mucous | 0 | 3.740597151 | 0.904 | 0.059 | 0 |
| LCN2 | Mucous | 0 | 3.71595707 | 0.986 | 0.075 | 0 |
| SAA2 | Mucous | 0 | 3.263902439 | 0.866 | 0.032 | 0 |
| SLPI | Mucous | 0 | 3.001902828 | 0.996 | 0.218 | 0 |
| SERPINA3 | Mucous | 0 | 2.725180975 | 0.723 | 0.027 | 0 |
| CXCL1 | Mucous | 0 | 2.121790394 | 0.796 | 0.119 | 0 |
| PI3 | Mucous | 0 | 2.112982356 | 0.627 | 0.012 | 0 |
| MMP7 | Mucous | 0 | 2.039034049 | 0.823 | 0.019 | 0 |
| RARRES1 | Mucous | 0 | 1.938771313 | 0.896 | 0.062 | 0 |
| C3 | Mucous | 0 | 1.768759158 | 0.951 | 0.094 | 0 |
| MDK | Mucous | 0 | 1.453983303 | 0.955 | 0.137 | 0 |
| CFB | Mucous | 0 | 1.402986853 | 0.868 | 0.071 | 0 |
| PDZK1IP1 | Mucous | 0 | 1.340861311 | 0.811 | 0.067 | 0 |
| CRISP3 | Mucous | 0 | 1.221590871 | 0.418 | 0.001 | 0 |
| NCOA7 | Mucous | 0 | 1.079051637 | 0.798 | 0.174 | 0 |
| CCL20 | Mucous | 6.66E-221 | 1.471254462 | 0.684 | 0.173 | 1.36E-216 |
| XBP1 | Mucous | 1.37E-188 | 1.05928154 | 0.886 | 0.421 | 2.79E-184 |
| CCL17 | Myeloid Dendritic Type 1 | 0 | 2.681912668 | 0.489 | 0.007 | 0 |
| CSF2RA | Myeloid Dendritic Type 1 | 1.01E-238 | 1.446571977 | 0.908 | 0.101 | 2.06E-234 |
| SERPINB9 | Myeloid Dendritic Type 1 | 4.79E-137 | 1.251191502 | 0.771 | 0.116 | 9.74E-133 |

| | | | | | | |
|---|---|---|---|---|---|---|
| RGS10 | Myeloid Dendritic Type 1 | 1.05E-99 | 1.103478553 | 0.916 | 0.236 | 2.14E-95 |
| HLA-DQA1 | Myeloid Dendritic Type 1 | 4.01E-89 | 1.891895134 | 0.992 | 0.453 | 8.16E-85 |
| C15orf48 | Myeloid Dendritic Type 1 | 5.97E-75 | 2.174083729 | 0.626 | 0.13 | 1.22E-70 |
| G0S2 | Myeloid Dendritic Type 1 | 4.33E-60 | 2.267628586 | 0.45 | 0.08 | 8.81E-56 |
| BIRC3 | Myeloid Dendritic Type 1 | 1.08E-32 | 1.308175361 | 0.618 | 0.224 | 2.20E-28 |
| AREG | Myeloid Dendritic Type 1 | 2.37E-19 | 1.66027885 | 0.473 | 0.215 | 4.83E-15 |
| CCL17 | Myeloid Dendritic Type 2 | 0 | 2.0634761 | 0.256 | 0.007 | 0 |
| HLA-DQA1 | Myeloid Dendritic Type 2 | 3.57E-164 | 1.762493144 | 0.989 | 0.452 | 7.27E-160 |
| AREG | Myeloid Dendritic Type 2 | 3.47E-39 | 1.014500968 | 0.523 | 0.214 | 7.07E-35 |
| LUM | Myofibroblast | 0 | 1.948272991 | 0.944 | 0.051 | 0 |
| COL6A2 | Myofibroblast | 0 | 1.375075631 | 0.863 | 0.1 | 0 |
| ACTA2 | Myofibroblast | 0 | 1.334396257 | 0.806 | 0.089 | 0 |
| DKK3 | Myofibroblast | 0 | 1.26977861 | 0.875 | 0.06 | 0 |
| RARRES1 | Myofibroblast | 0 | 1.269306754 | 0.694 | 0.066 | 0 |
| C1S | Myofibroblast | 0 | 1.259900004 | 0.847 | 0.071 | 0 |
| GEM | Myofibroblast | 0 | 1.169919284 | 0.669 | 0.047 | 0 |
| LTBP2 | Myofibroblast | 0 | 1.002560274 | 0.746 | 0.025 | 0 |
| FILIP1L | Myofibroblast | 9.21E-212 | 1.113113693 | 0.823 | 0.158 | 1.87E-207 |
| FHL1 | Myofibroblast | 1.26E-171 | 1.415063639 | 0.931 | 0.292 | 2.56E-167 |
| CCL2 | Myofibroblast | 1.47E-16 | 1.356867396 | 0.339 | 0.16 | 2.99E-12 |
| CCL5 | Natural Killer | 0 | 1.417388492 | 0.763 | 0.126 | 0 |
| CCL5 | Natural Killer T | 0 | 1.931881874 | 0.994 | 0.168 | 0 |
| KLRC2 | Natural Killer T | 0 | 1.136054204 | 0.693 | 0.029 | 0 |
| BTG1 | Natural Killer T | 2.36E-150 | 1.133920057 | 0.997 | 0.719 | 4.80E-146 |
| TSC22D3 | Natural Killer T | 4.84E-134 | 1.116378879 | 0.994 | 0.661 | 9.86E-130 |
| SCG2 | Neuroendocrine | 0 | 2.839338022 | 0.909 | 0.002 | 0 |
| SCG5 | Neuroendocrine | 0 | 2.29019825 | 0.909 | 0.003 | 0 |
| NEFL | Neuroendocrine | 0 | 1.626390742 | 0.364 | 0.001 | 0 |
| TUBB2B | Neuroendocrine | 0 | 1.151666149 | 0.727 | 0.004 | 0 |
| MB | Neuroendocrine | 8.24E-78 | 1.023829455 | 0.545 | 0.009 | 1.68E-73 |
| DNAJC12 | Neuroendocrine | 1.22E-77 | 1.018068258 | 0.636 | 0.013 | 2.48E-73 |
| MEG3 | Neuroendocrine | 5.81E-65 | 1.960754926 | 0.909 | 0.031 | 1.18E-60 |
| BEX2 | Neuroendocrine | 1.66E-24 | 1.487052024 | 0.818 | 0.073 | 3.38E-20 |
| PAM | Neuroendocrine | 5.45E-09 | 1.060749077 | 0.636 | 0.113 | 0.000110998 |
| VAMP2 | Neuroendocrine | 4.70E-08 | 1.034107247 | 1 | 0.432 | 0.000957604 |
| EIF4A2 | Neuroendocrine | 9.98E-08 | 1.097580045 | 1 | 0.559 | 0.002031893 |
| CIRBP | Neuroendocrine | 1.34E-07 | 1.153357668 | 1 | 0.746 | 0.00273753 |
| LST1 | Nonclassical Monocyte | 0 | 2.207635324 | 1 | 0.307 | 0 |
| CTSS | Nonclassical Monocyte | 0 | 1.374506188 | 1 | 0.529 | 0 |
| NAP1L1 | Nonclassical Monocyte | 0 | 1.261563474 | 0.98 | 0.478 | 0 |
| SAT1 | Nonclassical Monocyte | 0 | 1.200929686 | 1 | 0.845 | 0 |
| PSAP | Nonclassical Monocyte | 0 | 1.062619445 | 0.999 | 0.789 | 0 |
| G0S2 | OLR1+ Classical Monocy | 0 | 2.217297121 | 0.85 | 0.078 | 0 |
| SERPINB9 | OLR1+ Classical Monocy | 0 | 1.292378192 | 0.903 | 0.115 | 0 |
| ATP13A3 | OLR1+ Classical Monocy | 0 | 1.151570772 | 0.894 | 0.1 | 0 |

| | | | | | | |
|---|---|---|---|---|---|---|
| C15orf48 | OLR1+ Classical Monocy | 5.00E-258 | 2.248304193 | 0.865 | 0.129 | 1.02E-253 |
| IL1B | OLR1+ Classical Monocy | 4.04E-237 | 2.085705497 | 0.942 | 0.167 | 8.23E-233 |
| TNFRSF1B | OLR1+ Classical Monocy | 2.52E-224 | 1.288720633 | 0.865 | 0.15 | 5.12E-220 |
| ACSL1 | OLR1+ Classical Monocy | 6.18E-213 | 1.262036867 | 0.908 | 0.17 | 1.26E-208 |
| IER3 | OLR1+ Classical Monocy | 7.82E-197 | 1.312593618 | 0.87 | 0.167 | 1.59E-192 |
| GK | OLR1+ Classical Monocy | 1.67E-183 | 1.04648259 | 0.787 | 0.141 | 3.40E-179 |
| BCL2A1 | OLR1+ Classical Monocy | 1.47E-177 | 1.496522943 | 0.937 | 0.221 | 2.99E-173 |
| SOD2 | OLR1+ Classical Monocy | 7.96E-177 | 2.417967238 | 1 | 0.335 | 1.62E-172 |
| MARCKS | OLR1+ Classical Monocy | 1.96E-161 | 1.417300987 | 0.942 | 0.265 | 3.99E-157 |
| IFNGR2 | OLR1+ Classical Monocy | 6.29E-154 | 1.189193662 | 0.918 | 0.258 | 1.28E-149 |
| NAMPT | OLR1+ Classical Monocy | 7.29E-151 | 1.724893232 | 0.99 | 0.374 | 1.48E-146 |
| TNFAIP3 | OLR1+ Classical Monocy | 1.18E-147 | 1.264031264 | 0.908 | 0.254 | 2.40E-143 |
| S100A8 | OLR1+ Classical Monocy | 6.52E-140 | 1.992230888 | 0.918 | 0.282 | 1.33E-135 |
| SRGN | OLR1+ Classical Monocy | 2.62E-131 | 1.801445048 | 1 | 0.793 | 5.33E-127 |
| CD44 | OLR1+ Classical Monocy | 5.66E-129 | 1.415068876 | 0.995 | 0.529 | 1.15E-124 |
| UPP1 | OLR1+ Classical Monocy | 5.28E-127 | 1.009667986 | 0.918 | 0.303 | 1.08E-122 |
| S100A9 | OLR1+ Classical Monocy | 4.21E-118 | 2.225300076 | 0.947 | 0.442 | 8.57E-114 |
| SAT1 | OLR1+ Classical Monocy | 1.38E-112 | 1.425404055 | 1 | 0.847 | 2.81E-108 |
| WTAP | OLR1+ Classical Monocy | 1.32E-110 | 1.056550962 | 0.865 | 0.294 | 2.69E-106 |
| TYMP | OLR1+ Classical Monocy | 1.30E-103 | 1.057055136 | 0.966 | 0.426 | 2.65E-99 |
| TNIP3 | OLR1+ Classical Monocy | 6.43E-102 | 1.189542507 | 0.488 | 0.084 | 1.31E-97 |
| PLAUR | OLR1+ Classical Monocy | 7.87E-97 | 1.043533931 | 0.879 | 0.297 | 1.60E-92 |
| CEBPB | OLR1+ Classical Monocy | 2.71E-86 | 1.073387628 | 0.99 | 0.703 | 5.51E-82 |
| PNRC1 | OLR1+ Classical Monocy | 3.19E-82 | 1.044880784 | 0.976 | 0.642 | 6.49E-78 |
| FTH1 | OLR1+ Classical Monocy | 2.88E-81 | 1.16973151 | 1 | 0.999 | 5.87E-77 |
| NDUFA4L2 | Pericyte | 0 | 1.795066288 | 0.951 | 0.018 | 0 |
| IGFBP7 | Pericyte | 0 | 1.691507961 | 0.999 | 0.322 | 0 |
| MYL9 | Pericyte | 0 | 1.410912303 | 0.971 | 0.168 | 0 |
| COL4A1 | Pericyte | 0 | 1.332323038 | 0.876 | 0.124 | 0 |
| COL4A2 | Pericyte | 0 | 1.22262789 | 0.851 | 0.124 | 0 |
| EFEMP1 | Pericyte | 0 | 1.120212673 | 0.798 | 0.096 | 0 |
| STOM | Pericyte | 0 | 1.056087623 | 0.944 | 0.434 | 0 |
| COL6A2 | Pericyte | 0 | 1.035821438 | 0.831 | 0.084 | 0 |
| PAG1 | Pericyte | 0 | 1.006654809 | 0.746 | 0.157 | 0 |
| DNAJB9 | Plasma | 8.89E-194 | 1.25056027 | 0.898 | 0.175 | 1.81E-189 |
| HERPUD1 | Plasma | 1.34E-151 | 2.419982595 | 0.989 | 0.379 | 2.72E-147 |
| HSP90B1 | Plasma | 1.42E-124 | 1.947741479 | 0.984 | 0.514 | 2.89E-120 |
| SPCS3 | Plasma | 1.05E-121 | 1.102438496 | 0.877 | 0.271 | 2.14E-117 |
| SSR3 | Plasma | 1.80E-113 | 1.002058917 | 0.92 | 0.339 | 3.66E-109 |
| XBP1 | Plasma | 1.41E-103 | 1.585231182 | 0.941 | 0.423 | 2.88E-99 |
| BIRC3 | Plasma | 9.79E-103 | 1.164765868 | 0.802 | 0.223 | 1.99E-98 |
| TSC22D3 | Plasma | 2.62E-71 | 1.359799408 | 0.93 | 0.662 | 5.33E-67 |
| IRF4 | Plasmacytoid Dendritic | 0 | 1.049825978 | 0.759 | 0.01 | 0 |
| IRF7 | Plasmacytoid Dendritic | 3.68E-216 | 2.072052305 | 0.993 | 0.156 | 7.49E-212 |
| LTB | Plasmacytoid Dendritic | 3.20E-137 | 1.314870555 | 0.759 | 0.105 | 6.51E-133 |

| | | | | | | |
|---|---|---|---|---|---|---|
| UGCG | Plasmacytoid Dendritic | 8.75E-122 | 1.017679144 | 0.883 | 0.186 | 1.78E-117 |
| CCDC50 | Plasmacytoid Dendritic | 3.51E-116 | 1.112789308 | 0.839 | 0.182 | 7.15E-112 |
| PLP2 | Plasmacytoid Dendritic | 2.61E-99 | 1.157287125 | 0.949 | 0.303 | 5.32E-95 |
| HERPUD1 | Plasmacytoid Dendritic | 1.62E-82 | 1.311617096 | 0.978 | 0.379 | 3.30E-78 |
| AREG | Plasmacytoid Dendritic | 5.03E-40 | 1.339158755 | 0.65 | 0.214 | 1.02E-35 |
| C2orf88 | Platelet/Megakaryocyte | 0 | 1.264712452 | 0.625 | 0.008 | 0 |
| KIF2A | Platelet/Megakaryocyte | 3.33E-47 | 1.265350565 | 0.7 | 0.089 | 6.79E-43 |
| PLA2G12A | Platelet/Megakaryocyte | 1.18E-41 | 1.098241085 | 0.675 | 0.093 | 2.41E-37 |
| MAX | Platelet/Megakaryocyte | 8.44E-40 | 1.755674779 | 0.875 | 0.186 | 1.72E-35 |
| CA2 | Platelet/Megakaryocyte | 1.65E-35 | 1.448175531 | 0.75 | 0.136 | 3.35E-31 |
| RGS10 | Platelet/Megakaryocyte | 1.68E-30 | 2.079505257 | 0.85 | 0.237 | 3.42E-26 |
| LIMS1 | Platelet/Megakaryocyte | 2.65E-30 | 1.815790956 | 0.85 | 0.234 | 5.39E-26 |
| ODC1 | Platelet/Megakaryocyte | 3.49E-29 | 1.534518794 | 0.725 | 0.168 | 7.10E-25 |
| MPP1 | Platelet/Megakaryocyte | 5.98E-28 | 1.538108224 | 0.725 | 0.168 | 1.22E-23 |
| CCL5 | Platelet/Megakaryocyte | 1.72E-26 | 1.478776116 | 0.775 | 0.172 | 3.50E-22 |
| TLK1 | Platelet/Megakaryocyte | 2.91E-25 | 1.016967049 | 0.575 | 0.109 | 5.92E-21 |
| NCOA4 | Platelet/Megakaryocyte | 1.07E-21 | 1.974010353 | 0.725 | 0.235 | 2.18E-17 |
| ACTN1 | Platelet/Megakaryocyte | 2.99E-18 | 1.086182096 | 0.7 | 0.226 | 6.09E-14 |
| RNF11 | Platelet/Megakaryocyte | 1.18E-17 | 1.209985329 | 0.625 | 0.186 | 2.39E-13 |
| TPM4 | Platelet/Megakaryocyte | 2.89E-17 | 1.737158971 | 0.85 | 0.452 | 5.89E-13 |
| NAP1L1 | Platelet/Megakaryocyte | 3.40E-16 | 1.638362131 | 0.875 | 0.484 | 6.93E-12 |
| DAB2 | Platelet/Megakaryocyte | 2.48E-14 | 1.021696239 | 0.625 | 0.217 | 5.05E-10 |
| MYL12A | Platelet/Megakaryocyte | 3.24E-13 | 1.355213784 | 1 | 0.939 | 6.60E-09 |
| KRT15 | Proliferating Basal | 0 | 3.407924464 | 0.979 | 0.024 | 0 |
| SERPINB3 | Proliferating Basal | 0 | 2.169756874 | 0.723 | 0.014 | 0 |
| SERPINB4 | Proliferating Basal | 0 | 1.355115912 | 0.617 | 0.007 | 0 |
| PHGDH | Proliferating Basal | 9.43E-179 | 1.053567324 | 0.872 | 0.043 | 1.92E-174 |
| CLDN4 | Proliferating Basal | 2.39E-75 | 1.314475006 | 0.957 | 0.117 | 4.87E-71 |
| HMGA1 | Proliferating Basal | 4.68E-68 | 1.467549437 | 0.979 | 0.162 | 9.53E-64 |
| MDK | Proliferating Basal | 2.11E-64 | 1.671577677 | 0.915 | 0.142 | 4.29E-60 |
| IMPDH2 | Proliferating Basal | 4.39E-63 | 1.249016769 | 0.936 | 0.16 | 8.93E-59 |
| PLP2 | Proliferating Basal | 1.75E-39 | 1.153943634 | 0.979 | 0.304 | 3.57E-35 |
| RPLP0 | Proliferating Basal | 2.32E-29 | 1.217851725 | 1 | 0.917 | 4.72E-25 |
| EEF1G | Proliferating Basal | 1.08E-26 | 1.029309195 | 1 | 0.943 | 2.20E-22 |
| RPS4X | Proliferating Basal | 1.14E-26 | 1.239131761 | 1 | 0.978 | 2.32E-22 |
| RPL10A | Proliferating Basal | 2.88E-26 | 1.119468718 | 1 | 0.961 | 5.86E-22 |
| RPSA | Proliferating Basal | 3.36E-26 | 1.066625617 | 1 | 0.907 | 6.85E-22 |
| GAS5 | Proliferating Basal | 3.06E-25 | 1.035709113 | 0.957 | 0.427 | 6.22E-21 |
| RPS18 | Proliferating Basal | 3.23E-25 | 1.094736461 | 1 | 0.993 | 6.58E-21 |
| RPL3 | Proliferating Basal | 1.10E-23 | 1.051620079 | 1 | 0.992 | 2.24E-19 |
| MARCO | Proliferating Macrophag | 1.14E-173 | 1.482066785 | 0.996 | 0.276 | 2.33E-169 |
| ANP32B | Proliferating Macrophag | 4.03E-138 | 1.012402884 | 0.942 | 0.347 | 8.21E-134 |
| CTSC | Proliferating Macrophag | 1.16E-107 | 1.035996913 | 0.987 | 0.425 | 2.36E-103 |
| IL1B | Proliferating Macrophag | 5.19E-37 | 1.1376636 | 0.465 | 0.169 | 1.06E-32 |
| CCL5 | Proliferating NK/T | 1.57E-63 | 1.111912828 | 0.8 | 0.171 | 3.20E-59 |

| | | | | | | |
|---|---|---|---|---|---|---|
| ANP32B | Proliferating NK/T | 2.10E-63 | 1.111784203 | 0.905 | 0.348 | 4.28E-59 |
| KRT15 | Proximal Basal | 0 | 3.189724773 | 0.962 | 0.022 | 0 |
| SERPINB3 | Proximal Basal | 0 | 2.821699795 | 0.873 | 0.013 | 0 |
| SERPINB4 | Proximal Basal | 0 | 1.446209984 | 0.694 | 0.006 | 0 |
| CLDN4 | Proximal Basal | 6.88E-238 | 1.118066253 | 0.949 | 0.116 | 1.40E-233 |
| MDK | Proximal Basal | 9.98E-201 | 1.287877716 | 0.917 | 0.141 | 2.03E-196 |
| SLPI | Proximal Basal | 3.95E-149 | 1.705549977 | 1 | 0.222 | 8.05E-145 |
| GAS5 | Proximal Basal | 2.07E-107 | 1.32646858 | 0.981 | 0.426 | 4.22E-103 |
| RPL10A | Proximal Basal | 4.53E-98 | 1.293657293 | 1 | 0.961 | 9.21E-94 |
| RPS4X | Proximal Basal | 4.58E-97 | 1.333222904 | 1 | 0.978 | 9.31E-93 |
| RPLP0 | Proximal Basal | 4.86E-96 | 1.243300112 | 1 | 0.916 | 9.88E-92 |
| RPL15 | Proximal Basal | 1.53E-93 | 1.020477184 | 1 | 0.992 | 3.11E-89 |
| RPL3 | Proximal Basal | 6.82E-91 | 1.187600401 | 1 | 0.992 | 1.39E-86 |
| EEF1G | Proximal Basal | 1.51E-89 | 1.056147458 | 1 | 0.943 | 3.07E-85 |
| RPL7A | Proximal Basal | 3.75E-89 | 1.015896181 | 1 | 0.965 | 7.64E-85 |
| RPS18 | Proximal Basal | 1.11E-87 | 1.145426549 | 1 | 0.993 | 2.26E-83 |
| RPL12 | Proximal Basal | 1.06E-85 | 1.001940741 | 1 | 0.964 | 2.16E-81 |
| RPS6 | Proximal Basal | 2.32E-85 | 1.049517565 | 1 | 0.987 | 4.73E-81 |
| RPL13A | Proximal Basal | 1.25E-83 | 1.048634389 | 1 | 0.995 | 2.55E-79 |
| CD9 | Proximal Basal | 6.64E-80 | 1.099586533 | 1 | 0.665 | 1.35E-75 |
| CCDC146 | Proximal Ciliated | 0 | 1.8005813 | 0.966 | 0.032 | 0 |
| LRRC23 | Proximal Ciliated | 0 | 1.782314594 | 1 | 0.046 | 0 |
| CCDC170 | Proximal Ciliated | 0 | 1.75272191 | 0.989 | 0.038 | 0 |
| RSPH9 | Proximal Ciliated | 0 | 1.686258303 | 0.966 | 0.023 | 0 |
| DNAH5 | Proximal Ciliated | 0 | 1.359674554 | 0.92 | 0.03 | 0 |
| CDS1 | Proximal Ciliated | 0 | 1.21844205 | 0.989 | 0.048 | 0 |
| TMEM231 | Proximal Ciliated | 0 | 1.185144322 | 0.955 | 0.025 | 0 |
| PPIL6 | Proximal Ciliated | 0 | 1.167121636 | 0.898 | 0.028 | 0 |
| DYNC2H1 | Proximal Ciliated | 0 | 1.121533562 | 0.92 | 0.044 | 0 |
| DAW1 | Proximal Ciliated | 0 | 1.102150159 | 0.898 | 0.017 | 0 |
| DZIP3 | Proximal Ciliated | 0 | 1.097537092 | 0.898 | 0.044 | 0 |
| CCDC113 | Proximal Ciliated | 0 | 1.095573378 | 0.943 | 0.02 | 0 |
| STK33 | Proximal Ciliated | 0 | 1.08470662 | 0.886 | 0.024 | 0 |
| LRP11 | Proximal Ciliated | 0 | 1.083206231 | 0.898 | 0.039 | 0 |
| RFX3 | Proximal Ciliated | 0 | 1.0370727 | 0.909 | 0.044 | 0 |
| RP1 | Proximal Ciliated | 0 | 1.011050967 | 0.75 | 0.019 | 0 |
| DNAL1 | Proximal Ciliated | 7.23E-291 | 1.072831619 | 0.932 | 0.056 | 1.47E-286 |
| CLUAP1 | Proximal Ciliated | 4.32E-245 | 1.275572043 | 0.943 | 0.07 | 8.79E-241 |
| PFN2 | Proximal Ciliated | 1.75E-221 | 1.012363236 | 0.852 | 0.061 | 3.56E-217 |
| EFHC1 | Proximal Ciliated | 3.26E-215 | 1.678651304 | 1 | 0.094 | 6.64E-211 |
| CRNDE | Proximal Ciliated | 1.20E-204 | 1.581214001 | 0.989 | 0.096 | 2.45E-200 |
| C11orf74 | Proximal Ciliated | 1.51E-200 | 1.100813243 | 0.875 | 0.073 | 3.06E-196 |
| DMKN | Proximal Ciliated | 6.56E-184 | 1.13442815 | 0.943 | 0.092 | 1.34E-179 |
| MLF1 | Proximal Ciliated | 6.23E-135 | 1.203599814 | 0.943 | 0.13 | 1.27E-130 |
| ODF3B | Proximal Ciliated | 6.04E-126 | 1.992655448 | 1 | 0.178 | 1.23E-121 |

| | | | | | | |
|---|---|---|---|---|---|---|
| AZIN1 | Proximal Ciliated | 2.90E-116 | 1.00919294 | 0.966 | 0.157 | 5.90E-112 |
| NUCB2 | Proximal Ciliated | 1.57E-93 | 1.789429382 | 0.977 | 0.229 | 3.19E-89 |
| SLPI | Proximal Ciliated | 2.83E-75 | 1.092712484 | 1 | 0.223 | 5.76E-71 |
| SRI | Proximal Ciliated | 2.14E-65 | 1.23820179 | 0.989 | 0.417 | 4.36E-61 |
| CALM2 | Proximal Ciliated | 1.76E-49 | 1.074403239 | 1 | 0.914 | 3.58E-45 |
| C6orf58 | Serous | 0 | 5.154386519 | 0.875 | 0.004 | 0 |
| CRISP3 | Serous | 3.89E-261 | 1.456832454 | 0.458 | 0.004 | 7.93E-257 |
| DMBT1 | Serous | 1.62E-140 | 2.845846984 | 1 | 0.038 | 3.30E-136 |
| SERPINA3 | Serous | 3.55E-135 | 1.856884395 | 0.917 | 0.032 | 7.23E-131 |
| LCN2 | Serous | 5.55E-33 | 1.247056433 | 0.75 | 0.082 | 1.13E-28 |
| SLPI | Serous | 3.05E-31 | 5.020015739 | 1 | 0.224 | 6.20E-27 |
| NUCB2 | Serous | 2.48E-21 | 1.09353425 | 0.958 | 0.229 | 5.05E-17 |
| XBP1 | Serous | 2.24E-15 | 1.320433056 | 1 | 0.424 | 4.56E-11 |
| SLPI | Signaling Alveolar Epithe | 0 | 1.752039566 | 0.999 | 0.216 | 0 |
| MMP9 | TREM2+ Dendritic | 0 | 1.785258525 | 0.415 | 0.003 | 0 |
| C15orf48 | TREM2+ Dendritic | 1.84E-190 | 1.694985342 | 0.868 | 0.13 | 3.74E-186 |
| HLA-DQA1 | TREM2+ Dendritic | 1.87E-97 | 1.621903302 | 0.994 | 0.453 | 3.80E-93 |
| CTSB | TREM2+ Dendritic | 5.96E-95 | 1.375040707 | 0.994 | 0.489 | 1.21E-90 |
| TYMP | TREM2+ Dendritic | 6.22E-93 | 1.186156945 | 0.994 | 0.426 | 1.27E-88 |
| CD68 | TREM2+ Dendritic | 4.61E-89 | 1.072619875 | 1 | 0.328 | 9.38E-85 |
| CTSS | TREM2+ Dendritic | 4.17E-69 | 1.220293139 | 0.975 | 0.533 | 8.48E-65 |
| PSAP | TREM2+ Dendritic | 2.39E-66 | 1.260881602 | 1 | 0.791 | 4.86E-62 |
| FTH1 | TREM2+ Dendritic | 4.28E-57 | 1.047049261 | 1 | 0.999 | 8.72E-53 |
| NUPR1 | TREM2+ Dendritic | 7.99E-38 | 1.010904675 | 0.698 | 0.342 | 1.63E-33 |
| G0S2 | TREM2+ Dendritic | 2.17E-21 | 1.078858268 | 0.277 | 0.08 | 4.41E-17 |
| ACTA2 | Vascular Smooth Muscle | 0 | 3.244966786 | 0.994 | 0.085 | 0 |
| MYL9 | Vascular Smooth Muscle | 0 | 2.464883348 | 0.996 | 0.182 | 0 |
| IGFBP7 | Vascular Smooth Muscle | 0 | 2.389330509 | 0.994 | 0.333 | 0 |
| TPM1 | Vascular Smooth Muscle | 0 | 1.463259554 | 0.942 | 0.276 | 0 |
| COL6A2 | Vascular Smooth Muscle | 0 | 1.367266843 | 0.853 | 0.097 | 0 |
| MFGE8 | Vascular Smooth Muscle | 0 | 1.080051905 | 0.817 | 0.111 | 0 |
| NTN4 | Vascular Smooth Muscle | 0 | 1.028672235 | 0.695 | 0.098 | 0 |
| TGFB1I1 | Vascular Smooth Muscle | 0 | 1.009101859 | 0.774 | 0.115 | 0 |
| FILIP1L | Vascular Smooth Muscle | 1.96E-303 | 1.016281355 | 0.729 | 0.156 | 4.00E-299 |
| IGFBP7 | Vein | 0 | 1.444665206 | 0.98 | 0.327 | 0 |
| MT2A | Vein | 0 | 1.421660934 | 0.99 | 0.759 | 0 |
| SELE | Vein | 0 | 1.373857296 | 0.264 | 0.007 | 0 |
| MT1A | Vein | 0 | 1.278258068 | 0.57 | 0.129 | 0 |
| PLAT | Vein | 0 | 1.167429437 | 0.674 | 0.062 | 0 |
| LIFR | Vein | 0 | 1.138642244 | 0.814 | 0.117 | 0 |
| CD59 | Vein | 0 | 1.127939989 | 0.993 | 0.653 | 0 |
| CYP1B1 | Vein | 0 | 1.124874509 | 0.545 | 0.055 | 0 |
| EPAS1 | Vein | 0 | 1.06778236 | 0.979 | 0.348 | 0 |
| RGS5 | Vein | 0 | 1.053452403 | 0.713 | 0.076 | 0 |
| SRPX | Vein | 0 | 1.038424785 | 0.758 | 0.064 | 0 |

| | | | | | | |
|---|---|---|---|---|---|---|
| CXCL2 | Vein | 5.04E-303 | 1.421884137 | 0.675 | 0.253 | 1.03E-298 |
| CSF3 | Vein | 1.46E-255 | 1.085046807 | 0.39 | 0.095 | 2.96E-251 |
| CCL2 | Vein | 9.19E-239 | 1.175490734 | 0.501 | 0.155 | 1.87E-234 |
| IL6 | Vein | 6.08E-197 | 1.239169963 | 0.435 | 0.139 | 1.24E-192 |
